# Structure and dynamics of the essential endogenous mycobacterial polyketide synthase Pks13

**DOI:** 10.1101/2023.01.27.525930

**Authors:** Sun Kyung Kim, Miles Sasha Dickinson, Janet Finer-Moore, Ziqiang Guan, Robyn M. Kaake, Ignacia Echeverria, Jen Chen, Ernst H. Pulido, Andrej Sali, Nevan J. Krogan, Oren S. Rosenberg, Robert M. Stroud

## Abstract

*Mycobacterium tuberculosis* is currently the leading cause of death by any bacterial infection^1^. The mycolic acid layer of the cell wall is essential for viability and virulence, and the enzymes responsible for its synthesis are therefore front line targets for antimycobacterial drug development^2,3^. Polyketide synthase 13 (Pks13) is a module comprised of a closely symmetric parallel dimer of chains, each encoding several enzymatic and transport functions, that carries out the condensation of two different very long chain fatty acids to produce mycolic acids that are essential components of the mycobacterial cell wall. Consequently individual enzymatic domains of Pks13 are targets for antimycobacterial drug development^4^. To understand this machinery, we sought to determine the structure and domain trajectories of the dimeric multi-enzyme Pks13, a 2×198,426 Dalton complex, from protein purified endogenously from mycobacteria under normal growth conditions, to capture it with normal substrates bound trapped ‘in action’.

Structures of the multi-domain assembly revealed by cryogenic electron microscopy (cryoEM) define the ketosynthase (KS), linker, and acyltransferase (AT) domains, each at atomic resolution (1.8Å), with bound substrates defined at 2.4Å and 2.9Å resolution. Image classification reveals two distinct structures with alternate locations of the N-terminal acyl carrier protein (termed ACP1a, ACP1b) seen at 3.6Å and 4.6Å resolution respectively. These two structures suggest plausible intermediate states, related by a ~60Å movement of ACP1, on the pathway for substrate delivery from the fatty acyl-ACP ligase (FadD32) to the ketosynthase domain. The linking sequence between ACP1 and the KS includes an 11 amino acid sequence with 6 negatively charged side chains that lies in different positively charged grooves on the KS in ACP1a versus ACP1b structures. This charge complementarity between the extended chain and the grooves suggests some stabilization of these two distinct orientations. Other domains are visible at lower resolution and indicate flexibility relative to the KS-AT core. The chemical structures of three bound endogenous long chain fatty acid substrates with their proximal regions defined in the structures were determined by electrospray ionization mass spectrometry.

The domain proximities were probed by chemical cross-linking and identified by mass spectrometry. These were incorporated into integrative structure modeling to define multiple domain configurations that transport the very long fatty acid chains throughout the multistep Pks13 mediated synthetic pathway.

## Main text

*Mycobacterium tuberculosis* (*Mtb*) coevolved with humans and is currently the world’s most lethal bacterium, causing over 1.5 million deaths in 2018^1^. In *Mtb* and other mycobacteria including the non-pathogenic *Mycobacterium smegmatis (Ms)*, the outer layer of the cell envelope is a waxy, ~80 Å thick “mycomembrane”^5–7^ primarily comprised of 2-alkyl, 3-hydroxy long chain fatty acids termed mycolic acids. This outer layer is essential for mycobacterial viability as it is the principal permeability barrier against environmental stressors including innate immune molecules and antimycobacterial drugs. Because of their central role in mycobacterial physiology, the reactions in the mycolic acid synthesis pathway are targets for antimycobacterial drugs^3^. Drugs that target earlier parts of the pathway in *Mtb* include the front-line drug isoniazid and thioamides such as ethionamide. While most strains of *Mtb* are susceptible to the standard regimen antibiotics, approximately a half million people are diagnosed with multi-drug resistant TB each year^1^. Hence steps in the pathway of mycolic acid synthesis are pursued for new effective small molecule antimycobacterial drugs that can evade currently evolved drug resistance in *Mtb*. ^4,8^.

The final steps of mycolic acid synthesis ligate two very long chain fatty acid chains together in a condensation step carried out by a cytoplasmic multi-domain polyketide synthase 13 (Pks13)^9^ to form an α-alkyl β-ketoacyl thioester (Fig S1). Each Pks13 polypeptide chain begins with an acyl carrier protein toward the N-terminus (ACP1) (Fig 1A,B) that is activated to attach a prosthetic 4’-phosphopantetheinyl extension (Ppant), which has a terminal SH group, to S38 of the ACP1 domain. One of the substrates of Pks13 is generated from a long-chain fatty acid R1, containing aliphatic chains between 48 and 62 CH_2_ groups in length. The long-chain-fatty acid AMP ligase FadD32 activates R1 by ligation of the chain to AMP to produce the substrate of Pks13, meromycolyl-AMP^10–13^. This activated R1 is the equivalent of a ‘starter unit’ in a modular PKS. The terminal thiol of the Ppant group on ACP1 displaces the AMP to form a thioester with the R1 meromycolyl chain. This process also requires FadD32 in a role analogous to the loading domain in other PKSs^10,12,14^(Fig1B). In the amino acid sequence of Pks13 the ACP1 domain is followed by a ketosynthase domain (KS) (Fig 1A). The KS active site cysteine C267 reacts with the R1 substrate from the ACP1 by trans-thioesterification.

**Fig 1.**
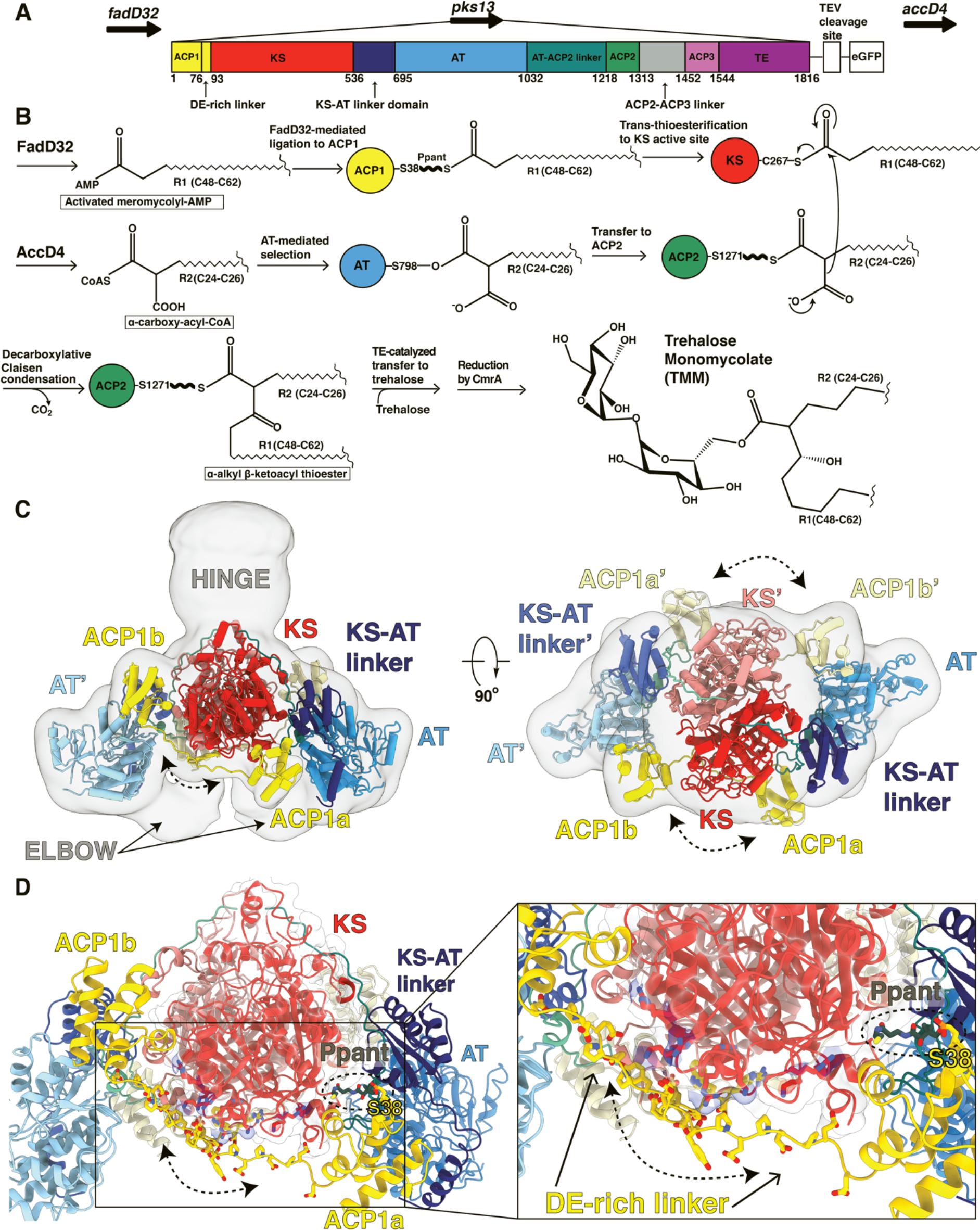
Overall architecture of the mycobacterial Pks13 showing two alternate positions of the ACP1 domain. (A) Part of the *Ms* operon containing the *fadD32 - pks13 - accD4* locus (top). Pks13 is comprised of 1816 amino acids with nine domains, and five domains of recognized function with lengths shown to scale in sequence. Colors are preserved throughout the Figures. An N-terminal acyl carrier protein (ACP1, yellow) is followed by an aspartate glutamate-rich linker (DE-rich linker), a ketosynthase (KS, red), a ketosynthase-acyltransferase linker domain (KS-AT linker domain, navy blue), an acyltransferase (AT, blue), AT-ACP2 linker (teal), two acyl carrier protein domains (ACP2, ACP3 domains, green and light purple) separated by a linker (grey), and the final thioesterase domain (TE, purple). ACP1 and ACP2 have the necessary -DSL- motif that needs to be phosphopantetheinylated to receive substrate. ACP3 lacks the -DSL- motif and hence no function can yet be ascribed to it. The C-terminus of Pks13 was tagged with a TEV-cleavage site and eGFP. (B) Schematic of the reaction carried out by the functional domains beginning with AMPylation of the meromycolyl substrate by FadD32, and transfer by FadD32 of the meromycolyl-AMP to the Ppant arm (bold wave line) of the N-terminal ACP (ACP1). This is followed by trans-thioesterification to the KS active site thiol for subsequent decarboxylative Claisen condensation. AccD4 carboxylates the second substrate to α-carboxy-acyl-CoA, which is then attached to the active site serine of AT. The α-carboxy-acyl branch is subsequently transferred to ACP2 and delivered to the KS active site for the decarboxylative Claisen condensation that ligates the two long substrate chains together. The α-alkyl β-ketoacyl thioester product is released from ACP2 by the TE that ligates it to trehalose to yield trehalose monomycolate (TMM) for subsequent export across the plasma membrane. (C) A composite structure of the Pks13 homodimer showing the relative positions of ACP1a and ACP1b fitted into a low-resolution composite cryo-EM map constructed from the focused reconstructions and incorporating the ACP1a density from that focused map, and the density from the ACP1b map. The two protomers are distinguished from each other by darker shades of domain colors yellow, red, and blue for one protomer, and lighter colors of the same for the other protomer. Domains in the lighter-colored protomer are labeled with the prime (‘) designation. The left Elbow region connects the lighter colored KS and AT domains. The right Elbow connects the darker colored domains. The Elbow and Hinge regions’ resolutions (~10Å) indicate flexibility and don’t allow model building. (D) The two positions of the ACP1 (ACP1a and ACP1b) are followed by acidic residues in the DE-rich linker (yellow) that electrostatically interact with basic residues on the KS (blue). Ppant attached to S38 is visible in ACP1a and labeled.

The second substrate is derived from a long-chain fatty acid R2 containing aliphatic chains between 24 and 26 CH_2_ groups in length. R2 is first esterified by coenzyme A (CoA) and carboxylated by an acyl-CoA-carboxylase AccD4, to yield an α-carboxy-acyl-CoA. This product is analogous to the ‘extender unit’ of modular PKSs. It is notable that the genes that encode FadD32 and AccD4 flank the gene for Pks13 (Fig 1A).

A 159 amino acid linker domain follows KS in the Pks13 chain and leads into the acyl transferase domain (AT). Serine S798 of the AT domain displaces CoA to form a serine ester with the carboxyl of R2. The AT is followed in the sequence by a 186-residue AT-ACP2 linker to a second Ppant-activated ACP (ACP2). R2 is captured by the sulfhydryl of the Ppant prosthetic group on ACP2 for transport to the KS active center where it undergoes a decarboxylative Claisen condensation with the R1 meromycolyl chain, with release of CO_2_. The condensed product results in a mycolic α−alkyl β-ketoacyl thioester attached to the Ppant arm of ACP2. This intermediate is transferred from ACP2 to a thioesterase (TE) domain toward the C-terminus of Pks13, which ligates the condensed product to trehalose. Subsequently the reductase CmrA reduces the ketone on the ligated intermediate to generate a mature trehalose mono mycolic acid (TMM) (Fig 1B) (Fig S1A)^16^. A third ACP-like domain (ACP3) is also encoded in the Pks13 sequence though it does not contain the active site serine necessary for attachment of the Ppant prosthetic group and therefore cannot perform the carrier function. It remains of unknown function in Pks13 (Fig 1A).

Though Pks13 is essential for *Mtb* there are no drugs yet that target individual enzymatic domains of Pks13. The general mechanism for how long-chain substrates are chaperoned and moved between domains has so far not been accessible for antimycobacterial target strategies^15^. We therefore sought to define the mechanism by which Pks13 processes these large hydrophobic substrates by determining the structure of the endogenous Pks13 purified from its *in vivo* state, with native long-chain substrates bound, rather than purified by overexpression in an exogenous host. This approach allowed us to visualize the multi domain enzyme in action and to uncover new functional sites in the substrate pathway that are unique to mycolic acid producing bacteria.

## Results

### Pks13 solubilized from native mycobacterial membranes is a stable dimeric complex

*Ms* Pks13 shares 71% amino acid sequence identity with that of *Mtb*, carries out the same enzymatic reaction, and so is a model for Pks13 in the human pathogen *Mtb*^9^. The *pks13* locus was tagged at the C-terminal end of the endogenous gene in *Ms*^16^ so as not to impair endogenous expression or function, with a TEV-cleavable Green Fluorescent Protein ‘GFP’ (Fig 1A). The GFP fusion was used to enable native purification using anti-GFP nanobody beads. Cleavage of Pks13 from GFP using TEV protease yielded full length Pks13 with an additional five-amino acid linker (-SKSTS-) followed by six amino acids from the TEV cleavage site at its C-terminus.

Detergent was required for solubilization and purification, suggesting that Pks13 is attached to the mycobacterial plasma membrane, as suggested based on live cell imaging^16^. Various detergents were evaluated for optimal solubilization of Pks13 using fluorescence size exclusion chromatography (methods). The non-denaturing detergent N-dodecyl-β-D-maltoside (β-DDM) was selected based on efficiency of solubilization. Following affinity purification using anti-GFP nanobody beads and on-bead cleavage by TEV, Pks13 was run through a sizing column, which gave two peaks (Fig S2A). The protein in each peak was comprised of Pks13 as seen on SDS-PAGE, and was predominantly dimeric Pks13 as shown by ‘blue native’-PAGE (Fig S2B,C). Liquid Chromatography with tandem mass spectrometry (LC-MS-MS) confirmed the identity with 78.1% sequence coverage throughout the entire Pks13 (Fig S2D). Therefore, the higher molecular weight peak1 resolved in SEC is a larger multimer that readily dissociates into a dimer.

Cryogenic electron microscopy (cryoEM) was used to image particles both from peak1 and peak 2 fractions (Table S1). We added a benzofuran inhibitor of the terminal thioesterase (TE) domain ^17^ ‘TAM16’ to part of the material, with the aim of stalling the throughput of the overall reaction, and producing a more homogeneous stable structure. With and without this inhibitor produced the same structures with marginally better resolution with inhibitor validating the concept. The peak 1 fraction revealed multiple conformational states of the dimeric protein, whose structures were determined by cryoEM (Fig S3, S4). Focused three-dimensional classifications were used to resolve domains and substructures within the complex (Table S2). Resolutions of the classified structures and domains were estimated by map-to-model comparisons using Q scores and by Fourier shell correlations (FSC) between maps constructed from independent sets of images (Table S2, S3). The most ordered parts of the structures that include the KS, KS-AT linker, and the AT domain were at 1.8Å atomic resolution. The proximal parts of the bound substrates attached to the KS and AT were seen at 2.4Å and 2.9Å resolution respectively (Fig2C, 3A) (Fig S5). Two positions were seen for the most N-terminal acyl carrier protein (ACP1a, ACP1b) at 3.6Å, 4.6Å respectively (Fig 1C,D) (Fig S4).

**Fig 2.**
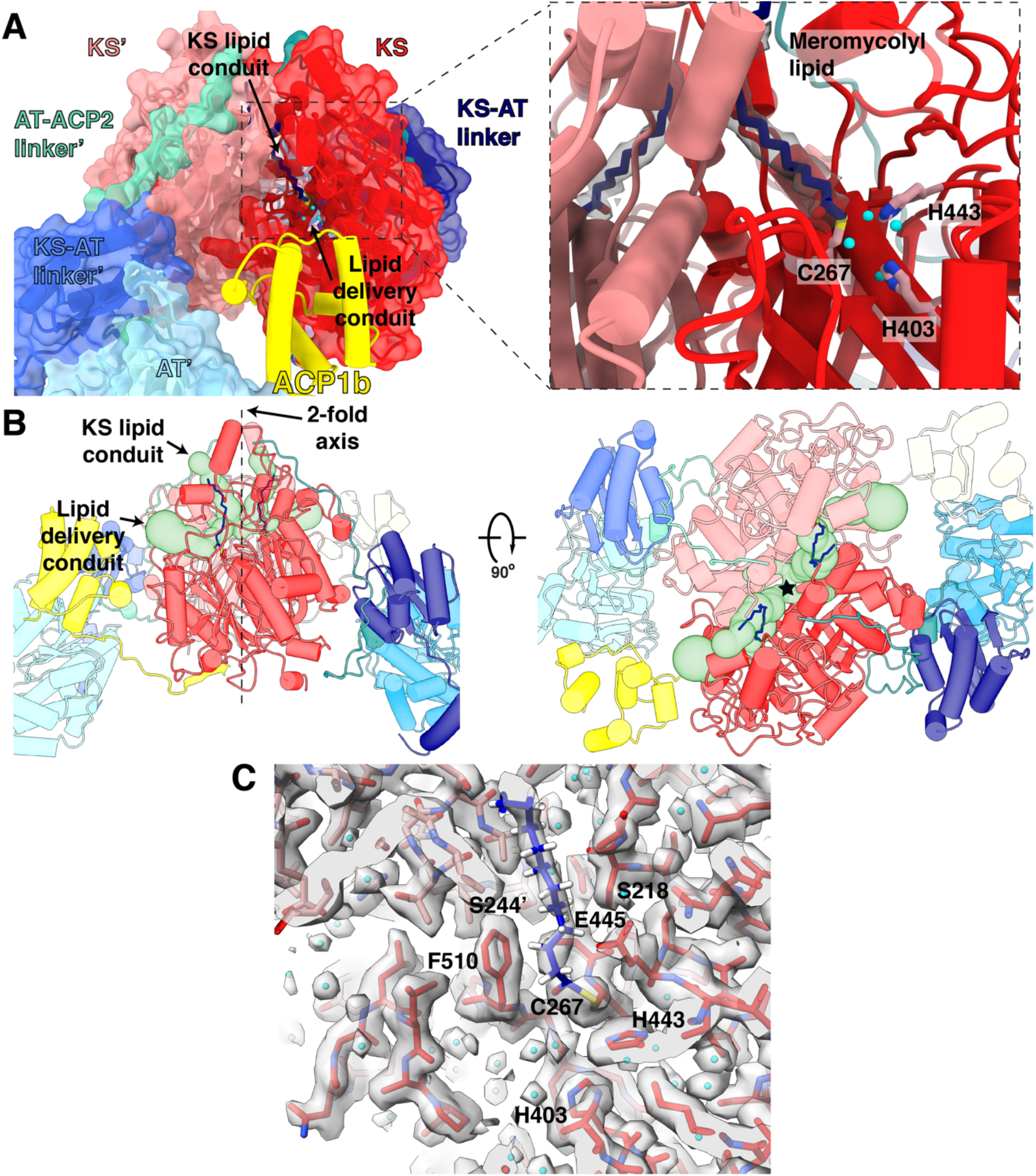
Meromycolyl substrate in the KS active site. (A) A view of the ACP1b (yellow) near the KS (red) of the same protomer near the putative lipid delivery conduit. Inset: Close up view of the KS active site. The meromycolate substrate is a thioester adduct with C267 and density for the lipid is shown within 2 Å about the carbon atoms. Active site residues C267, H443, and H403 involved in trans-thioesterification are shown in pink. Water molecules within hydrogen bonding distance of active site residues are shown as cyan spheres. (B) Front (left) and top (right) views of the ACP1b conformer structure with two branches of a hydrophobic tunnel (KS lipid conduit and lipid delivery conduit) shown as green surface models. The approximate location of the 2-fold axis is indicated as a dotted line and labeled in the front view (left) and by a star in the top view (right). (C) Density for the catalytic site and surrounding regions and water molecules, and the proximal portion of the meromycolate substrate attached to C267.

**Fig 3.**
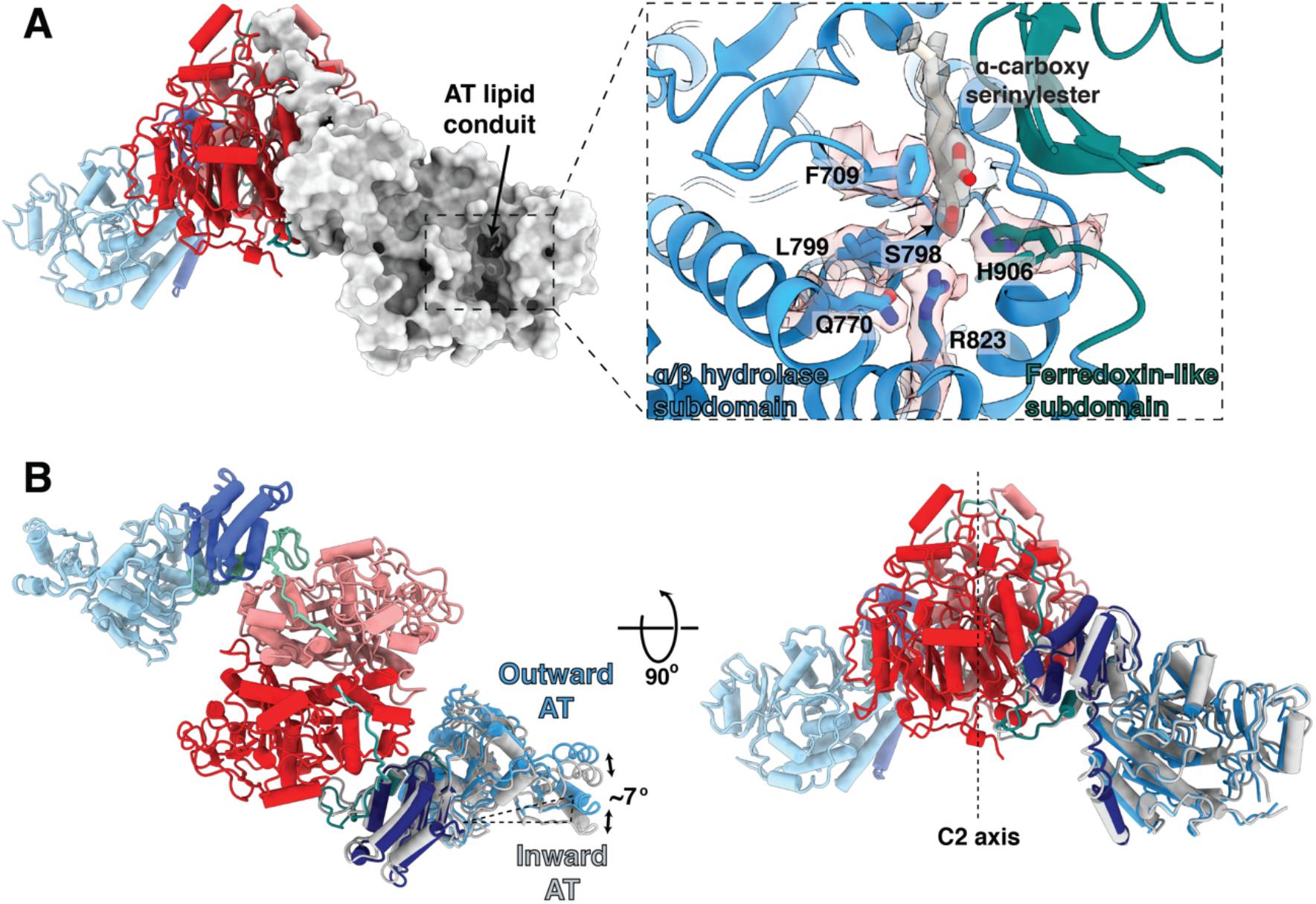
Substrate-bound AT active site, and alternate inward and outward positions of the AT domain. (A) Cartoon and surface representations viewed perpendicular to the vertical two-fold axis showing the lipid conduit in the AT domain. The inset shows an expanded view of the density for the α-carboxy lipid substrate in the AT lipid conduit between the αβ hydrolase subdomain (blue) and the ferredoxin-like subdomain (dark cyan). The α-carboxy lipid is attached as an ester linkage at S798. Surrounding active site residues, including the catalytic base H906, are labeled and shown with corresponding density. Density and atomic structure for the entire region is shown in stereo in Fig S5. (B) Two perpendicular views of the AT conformers show the movement ~7° outward from their hinge point with respect to the KS dimeric core. The inward conformation is shown in gray, elaborated in Movie S1.

### The ketosynthase dimer is bound to meromycolyl substrate chains in a pre-condensation state

Pks13 is a parallel homodimer of the multi-domain chains driven by the dimeric KS interface^18,19^ (Fig 1C). The interface area between the KS domains is 2400Å^2^, for a buried surface area of 4800Å^2^ consistent with its role as a tight and specific dimeric interaction. The calculated solvation free energy gain upon interface formation is Δ^i^G of −22.2 kcal/mol^20^. Substrate density protruding from each of the catalytic C267 residues in the KS dimer core (Fig 2) extends outward through one branch of a hydrophobic tunnel towards solvent, close to the two-fold axis labeled ‘KS lipid conduit’. Covalently bound fatty acid chains in the complex were identified by hydrolysis and mass spectrometry. Following identification of the attached chains (see below) we determined the chain attached to C267 is a statistical proportion of the meromycolic chain belonging to the α_2_ mycolic acid (C_55_H_106_O_2_), and the α’ mycolic acid (C_40_H_78_O_2_) ^21^. These two alternative chains are activated by FadD32 AMPylation, and transferred onto the thiol of the Ppant arm attached to S38 of ACP1 with release of AMP^22^ (Fig 1B), and subsequently translocated into the KS active site by trans-thioesterification (Fig 1B). The α2 or α’ substrates seen attached to the KS active site are therefore in the pre-condensation state.

Multiple ordered water molecules are seen within hydrogen-bonding distances of the conserved KS catalytic triad C267, H403 and H443^10–12,18,23^ (Fig 2). The active center and the first 18 CH_2_ groups of the substrate acyl chain embedded within the KS are well-defined in density at 2.4Å resolution (Fig 2C, Table S3). The remaining 22, and 37 carbons of the distal parts of these chains reside outside the KS domain and are not visible in density indicating that they are less well ordered. They may possibly be sequestered by the detergent in our preparation.

### ACP1 alternates between two positions that suggest a pathway for meromycolyl transport to the KS

Two stable alternative positions of the ACP1 (ACP1a and ACP1b) with respect to the KS-AT dimeric structure are well defined and separated in focused 3D image classifications of the structures. They are superimposed in a composite density map that superposes the image reconstructions of the focused image classification and refinements to display their relative positions (Fig 1C). These two states of Pks13 were each refined to ~ 2.4 Å average resolution, with lower resolutions for the ACP1a (3.6Å resolution) and ACP1b (4.6Å resolution) regions (Table S3)(Fig 1C,D)(Fig S3,S4, Movie S1).

ACP1a and ACP1b each have continuous density for residues 76-92 that connect ACP1 to the downstream KS. This 17-amino acid linking sequence (E76-R92) is negatively charged with nine acidic amino acids (E76, E78, E80, E82, D85, E86, D87, E88, D89), termed the ‘DE-rich linker’. 11 amino acids in this sequence (E76-E86) occupy either of two positively charged grooves along the ‘guiding surfaces’ of the KS (Fig 1D)(Fig S6,S7). These positively charged groove surfaces are in stark contrast to the otherwise predominantly negatively charged electrostatic surface of the KS (Fig S7). The DE-rich linker adopts an extended chain configuration perhaps assisted by its proline content (P79, P81, and P83) (Fig 1C,D).

The distance between ACP1a and ACP1b is approximately 60 Å with ~180° rotation of ACP1. ACP1a makes contacts with the KS, KS-AT linker domain, and the AT-ACP2 linker via its helix 2 and 3, and the 2-3 loop joining them (Fig S8). This position is not seen in other PKSs or fatty acid synthases. The total buried surface area of 1125 Å^2^ is consistent with it being a transient association (Fig 1C,D, S8)^24^. Analysis of the interface using PISA suggests that it is an unusually hydrophobic interface with a ‘predicted solvation free energy’ of formation ΔG^i^ of −5.5 kcal/mol^20^. There is one hydrogen bond in the interface, between the backbone amide of R39 of ACP1a and the carbonyl oxygen of A645 in the KS-AT linker domain, and there is a notable pi-pi stacking interaction between guanidinium groups of R63 on the ACP1a and R496 on the KS (Fig S8A,B). These interactions add specificity to this ACP1a-KS-AT linker orientation. The Ppant arm attached to S38 of ACP1a is clearly resolved in density maps, and shows that it is not bound to a fatty acid chain (Fig S8C).

The second ACP1 position, ACP1b, is much closer than ACP1a to the second branch of the KS hydrophobic tunnel that forms a putative ‘delivery conduit’ into the KS active site (Fig 2A,B). ACP1b contacts the KS from the same protomer with a total of 495 Å^2^ buried surface area and a single hydrogen bond between the backbone amide of T59 and the carbonyl oxygen of P476 (Fig S9B). The interface with KS is otherwise hydrophobic, with −G^i^ of −4.0 kcal/mol^20^. ACP1b also contacts the AT’ from the opposite protomer with a buried surface area of 129.5 Å^2^. This interface is more hydrophilic than the interface with KS and includes two hydrogen bonds that may add specificity to the overall interaction (Fig S9A)^20^. S38 of ACP1b which receives the Ppant arm, is 27 Å away from the KS active site C267 of the same protomer (Fig S10), where the meromycolyl substrate is seen attached in the structure. The Ppant arm seen in ACP1a is not visible on ACP1b due to limited resolution there, indicating that it is then too flexible to be resolved.

The negatively charged nature of the DE-rich linker, though not the sequence itself, and the two positively charged grooves are conserved among Pks13 homologs from other mycobacterial species (Fig S11). This suggests a role of this linker and its interactions in guiding the dynamic trajectory of ACP1 for meromycolate delivery from FadD32 to the active site of the KS domain.

The well-populated image classes for ACP1a versus ACP1b (Fig S3) for each position in an equilibrium sample on the em grids is evidence that the free energy difference between these two ACP1 states is small (see discussion), consistent with ACP1 function in readily alternating between these two positions to facilitate forward translocation of the meromycolyl chain.

### The acyltransferase domain reveals the binding site for the α-branch substrate chain

The second much shorter ‘α-branch’ substrate chain (the α-carboxy-acyl-CoA (C_24_-C_26_) is brought in from AccD4 to the AT active site where the catalytic serine (S798) displaces CoA^25^ to form an ester with the R2 substrate. In the AT structure, this endogenous substrate is bound to S798 with density at 2.9Å resolution for the first nine carbons of the acyl chain from the carboxylated serine ester. This proximal end of the substrate occupies a hydrophobic tunnel termed the AT lipid conduit (Fig 3A). This chain is analogous to the ‘extender unit’ of modular Pkss ^10,26^ (Fig 1B) (Fig S1). The α-carboxy-acyl chain would subsequently be transferred from the AT to the Ppant arm thiol bound to S1271 of the downstream ACP2 (aa. 1218-1313). From there the ACP2 thioester and R2 chain would be inserted into the KS active site to undergo decarboxylative Claisen condensation with the meromycolyl-cysteine adduct at C267 (Fig 2A). Decarboxylation of the α-carboxy-acyl chain either generates a nucleophilic carbanion that attacks the KS-bound electrophilic thioester, or undergoes a concerted reaction to yield the α-alkyl β-ketoacyl thioester product, the direct precursor of mycolic acid^9^ (Fig 1B)(Fig S1). The R stereochemistry at position 2 that bears the carboxyanion is conserved in all mycolic acids showing that the Pks13 reaction is stereospecific^13^.

The native α-carboxylated substrate of the AT domain lies in a cleft between the α/β hydrolase and ferredoxin-like subdomains of the AT (Fig 3A, Fig S12, Movie S1). The carboxylate of the carboxyacyl serine ester intermediate is stabilized by anion-*π* stacking^27^ with the phenyl ring of F709 (Fig 3A, Fig S5A,B). The *ε*N of the catalytic base H906 is hydrogen bonded to both the O*γ* of S798 (2.8Å) and to the carbonyl oxygen of the substrate ester (3.5Å), supporting its action as both the catalytic base and as a stabilizing force for the carbonyl oxyanion intermediate during the reaction. The imidazole of H906 is pinioned by hydrogen bonds from the *δ*N-H to the carbonyl oxygens of G959 and L960. The N*ε*2 of H906, O*γ* of S905, and NH2 of positively charged R823 are all oriented to stabilize the oxyanion formed during the catalytic reaction (Fig S5A,B).

When compared with the structure of a 52 kDa AT-containing fragment of *Mtb* H37Rv^25^ in complex with an α-carboxy palmitoyl substrate mimetic, Pks13AT had an RMSD of 1.00 Å versus the fragment, however the long fatty acid substrate density lies in a different solvent-exposed hydrophobic tunnel (‘lipid conduit’ in Fig 3A, Fig S12 Movie S1), and the Pks13 active site differs in that the amidinium cation of R823 does not form a bidentate salt bridge with the α-carboxy group of the α-fatty acid^25^ but is a key part of the oxyanion binding site formed during the reactions. These differences between the isolated AT and our multi-domain structures may reflect conformational restraints introduced by interactions of the AT domain with the KS and ACP1 domains in the native Pks13 environment. Such restraints may modulate access for substrate translocation to the ACP2.

KS and AT domains are connected by a flexible “elbow” that incorporates 59 unmodeled amino acids between residues 529 and 588 with its general location defined in the density (Fig 1C, Fig S13). An amphipathic helix (residues 588-608) found only in mycobacterial Pks13s leads into the KS-AT linker domain (Fig S13). Two discrete alternate conformations of the AT domain against the KS domain are seen in image classes and each were subjected to focused refinement of the individual AT domains. The two positions of each AT are rotated by 7° relative to the KS-AT linker domain (Fig 3B, Movie S1). However, this change cannot yet be correlated with any particular structural change in Pks13. The AT-ACP2 linker (residues 1032-1074) is well defined as an extended chain along the surface of the KS domain. It remains possible that this conformation change may accompany repositioning of some of the other domains that are C-terminal to the AT-ACP2 linker.

### Regions following the post-AT linker are outlined by low resolution envelope densities, without specific assignments

The ‘Hinge’ region (Fig 1C) and two lenticular densities ‘on top’ of the 2-fold axis of the KS-AT domains (Fig 4) account for some or all of the remaining domains of the structure: these domains comprise a 143 amino acid linker (1075-1217), ACP2, a 138 amino acid linker, ACP3, and the TE domain (Fig 4A,B). These two three dimensional image classes resolved at low (~20Å) resolution were not rotated by 180° to average them since any asymmetric single image class identified for the overall dimeric assembly has two-fold equivalent position with identical interactions of the lenticular densities. Taking account of the two-fold symmetry of the dimer, this pair of lenticular shaped densities have two alternate positions ~40° apart around the ‘Hinge’ region. This implies flexibility around the Hinge region to allow this movement in transferring substrates between the domains, and accounts for the lower resolution in these regions. This is consistent with the need for the ACP2, and TE domains to access and act in concert to transport the product between multiple different sites within the entire Pks13.

**Fig 4.**
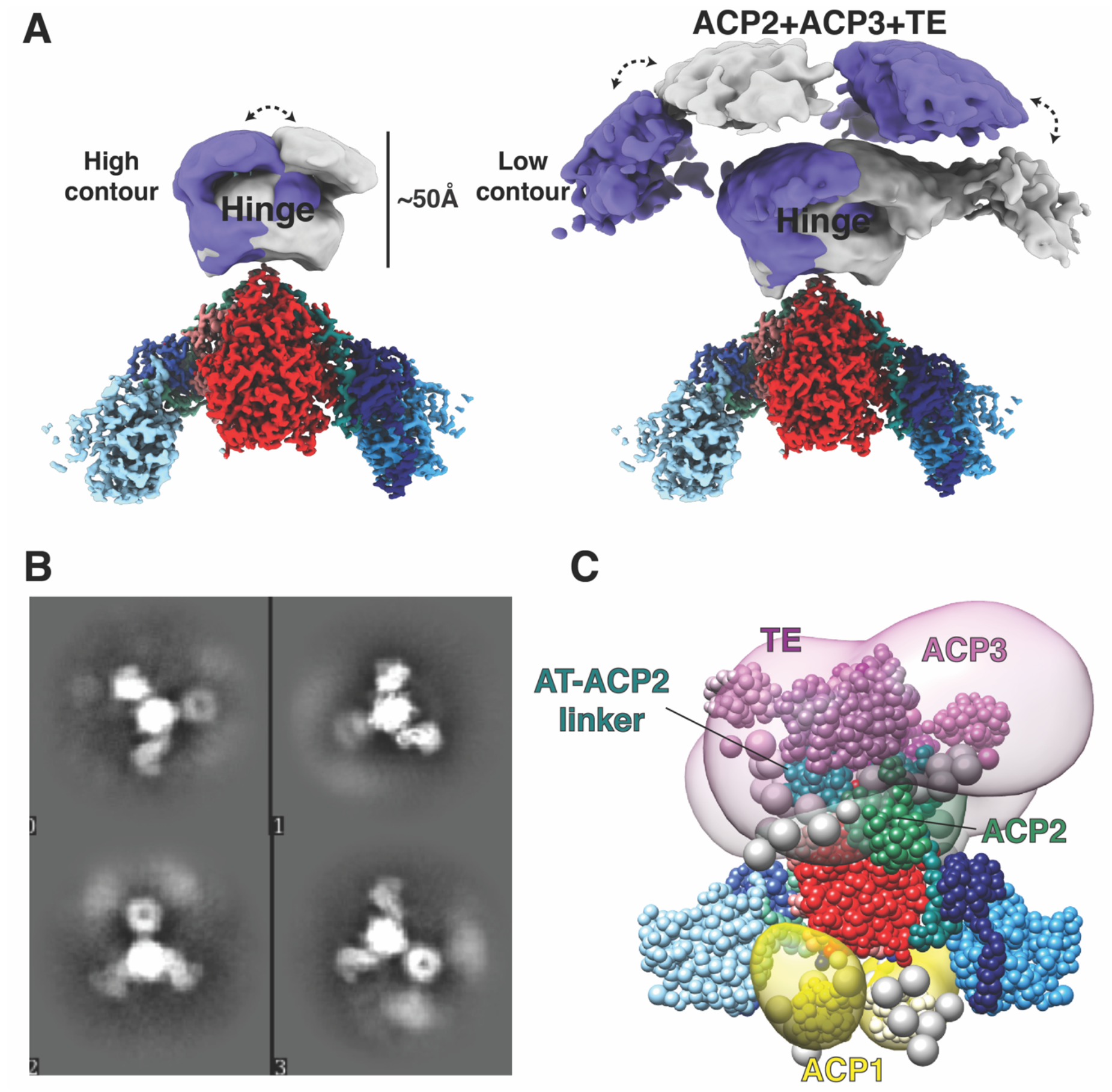
C-terminal domains are visualized at low resolution in 3D classification in two alternate positions. (A) The Hinge and C-terminal domains are visible in the cryo-EM map above the KS-AT domains. Two conformations of the C-terminal domain densities were recovered from 3D classification, shown in purple and grey respectively. These image classes are two-fold related but shown as alternative structures to emphasize the movement of the Hinge and the associated domains. (B) 2D classes showing smeared density attributed to the ACP2, ACP3 and TE domains that have multiple positions around the Hinge domain. (C) Integrative structure model of the Pks13 dimer based on the cryoEM structure and chemical cross-linking. The model represents the centroid of the ensemble of good scoring models obtained through Monte Carlo sampling. Regions with available atomic structures were modeled as rigid bodies, while regions lacking atomic structure were modeled as coarse-grained flexible beads. The structural ensemble is presented as a 3D localization probability density whose surface is rendered transparent for visual clarity, using the same color scheme for domains as Fig. 1. Complete analysis of all classes is shown in Fig S20. Two-fold symmetry was applied to compute the model.

### Identification of the bound substrate chains

The proximal regions of ligands at the active sites of the KS and AT domains are well defined in the density at 2.4Å and 2.9Å resolution respectively (Table S3). As the bound fatty acid chains emerge from the protein tunnels that emanate from the active sites the density for the hydrophobic distal regions is more diffuse consistent with less specific binding of the aliphatic chains outside of the tunnel regions. The substrates attached to Pks13 were identified by hydrolysis followed by (normal phase) liquid chromatography coupled with electrospray ionization high resolution mass spectrometry (NPLC-ESI-HRMS). Their predicted and observed mass-to-charge ratios (m/z) accurate to three decimal places are listed in Table S4. Three very long chain fatty acids series were identified: 1) the longest series is centered around C_55_H_106_O_2_ with exact mass 798.819 Da. (Fig S14) and corresponds to R1. It is consistent with being the “alpha 2” mycolate (C_51_-C_57_) found in *Ms*^28,29^, seen attached to C267 of the KS structure (Fig2). 2) A series centered around C_40_H_78_O_2_ with exact mass 590.600 Da.(Fig S15) corresponds to an alternative R1, and is termed the “alpha-prime” mycolate of length C_36_-C_42_ with C_40_ being most represented. It contains an unsaturated double bond followed by a methyl substituent in the chain. The proximal portions of the C_55_ or the alternate C_40_ fatty acyl chains are seen at 2.4Å resolution attached to C267 in the KS structure. They are in the state prior to the Claisen condensation (Fig 2) (Fig S1). 3) A series centered around C_24_H_48_O_2_ with exact mass 368.365 Da. (Fig S16) corresponds to R2, a C_22_-C_26_ fully saturated chain with C_24_ as most represented (termed lignoceric acid) (Fig 1B). This fatty acid is consistent with the α-carboxy acyl chain attached to the AT active site as seen in the density (Fig 3) (Fig S5). The mass however indicates that it lacks the α-carboxy group expected in the chain that is seen in the density maps. The carboxyl group may have been released by the hydrolysis from Pks13, or during mass spectrometry. The residue-specific attachment sites of fatty acids to the peptide chain, previously established for *Mtb*^10^ were not further reverified experimentally due to the difficulty of obtaining mass spectra of these long chain fatty acids attached to peptides.

### Proximity of domain locations and integrative restraints on trajectories

The diffuse densities for regions of Pks13 that are C-terminal to the AT-ACP2 linker in linear sequence suggest structural flexibility and a heterogenous localization of these domains, as required to transfer substrates. To determine the localization of the flexible domains in solution under native conditions, we performed cross-linking mass spectrometry (XL-MS). We used a disuccinimidyl sulfoxide (DSSO)-based XL-MS^3^ approach ^30,31^ wherein DSSO covalently links primary amines of the epsilon nitrogens of lysine (K) or the N-terminus of the protein. Following enzymatic digestion, resulting peptides are separated by liquid chromatography (LC) and analyzed by multi-stage MS: 1) intact peptide ions are measured at the MS1 stage; 2) using a lower energy collision-induced dissociation (CID) the DSSO backbone is cleaved while leaving peptide backbones intact, and the resulting fragment ions are measured at the MS2 stage; 3) each resulting ion is isolated in the instrument and the peptide backbone is fragmented using standard higher-energy C-trap dissociation (HCD) for peptide sequencing at the MS3 stage. Using this approach we can distinguish between inter-linked and ‘dead-end’ or mono-linked peptides^32^. Inter-linked peptides are the result of two linked K residues or the N-term that are between two separate peptide species (α and β). Mono-linked peptides are formed when a K or N-term residue reacts with DSSO, and before a second residue can form a bridge, the DSSO hydrolyzes resulting in a dead-end. The spacer arm that bridges linked residues is 10.3 Å. When estimating distances between α-carbons of inter-linked lysine amino groups or the N-terminal amine we allow for rotational freedom, therefore we classify a cross-link as satisfied if the Cα–Cα distance spanned by the cross-linked residues is less than 35 Å^33^.

We cross-linked natively purified Pks13 with increasing amounts of DSSO at 37°C and at 4°C (Fig S17). All bands associated with cross-linked products were excised and analyzed by XL-MS^3^ (Fig S17). In total, we identified 1435 inter-links and 2300 dead-end links including redundant counts, with a false positive rate of 1% as calculated by the identification of decoys (Table S5, Fig S18A). This included a small number of linkages between identified contaminating proteins (29 inter-links and 53 mono-links) (Table S5). In total, the number of redundant Pks13 inter-links corresponded to 96 unique K-K linkages, 63 of which could be mapped to the ACP1a or ACP1b configurations of Pks13 with 50.8% of those found within the expected α carbon distance of 35 Å, and a high number of long-distance violations (>50 Å) (Table S6, Fig S18B). Among the 96 unique linkages we identified 8 that could only come from oligomeric structures between at least two dimers of Pks13, all of which when mapped to either the ACP1a or the ACP1b configurations of Pks13 were beyond expected distances. Suspecting that some of the highly cross-linked samples were the product of aggregation or higher order oligomeric structures, we re-analyzed the data but removed the data files that were associated with highly cross-linked samples (Table S6 and Fig S17). Using this filtered XL-MS^3^ dataset, we identified 604 redundant inter-links which corresponded to 57 unique linkages (Table S7; Fig S18C). This corresponded to 33 linkages that mapped to ACP1a or ACP1b configurations, with 67% of those within the expected distances, and only one long-distance violation (Table S7; Fig S18D). Most of the violations can be accounted for by small movements or flexibility in the structure. Therefore, in order to model the domains not seen in ACP1a or ACP1b configurations, and to capture flexibility in the in-solution structure, we used the 57 unique linkages (Fig S19) from the filtered XL-MS dataset for integrative modeling.

Lysine 677 forms several cross-links to other domains that are present in the protein sample but defined only as diffuse domains in the structure (Fig 4), and the cross-links then support that the ACP2, ACP3, and the TE domain come close to the AT domain as required for ACP2 to transport the α-chain to the KS active site to evoke the final condensation. While the majority of the 34 cross-links formed within the atomistic portions of the Pks13 dimer structure are consistent with known distances in the dimeric structure, several are too far apart (FigS18, Table S6,7). These can only be consistent with higher order assemblies of Pks13 dimers as indicated in the dimer-of dimers seen as peak 1 in the size exclusion profile (Fig S2).

A structural model of the Pks13 dimer was computed by integrative modeling^34–38^ based on the 57 chemical inter-links (Fig. S19) and structural models of the components of the Pks13 dimer. These components include the current atomic structural model based on the cryo-EM map at 1.8 Å resolution, *de novo* AlphaFold2 predictions of domains unresolved in the cryo-EM structure^39,40^, and flexible linker regions (Fig. S20A). The cross-links were obtained from populations of structurally dynamic protein complexes as they exist in solution. A model of the Pks13 dimer was computed by satisfying this input information to the best possible degree using IMP^41^. The resulting ensemble of acceptable models satisfies 91% of the cross-links (Fig. S20B,C). The unsatisfied (over 35Å) cross-links span mostly residues in the KS and AT domains, indicating that the rigid representation of the system is not adequate to capture the full range of conformations in solution. The integrative structure localizes the AT-ACP2 linker and ACP2 domain relatively precisely to the ‘Hinge’ region of the cryo-EM density map and suggests large uncertainty in the configuration of the ACP3 and TE domains within an arch-like volume above the ‘Hinge’ (Fig 4) or around the KS and AT domains (Fig. S20A). This uncertainty can in principle arise from both the actual structural dynamics of the domains, and relative lack of input information; it is difficult to deconvolute the two possibilities. Furthermore, the model indicates that the distances between the active sites of the ACP1(S38)-KS(C267), KS(267)-ACP2(S1271), and ACP2(S1271)-TE(S1616) domains are highly variable, but that these active sites come in proximity (Fig. S20D).

## Discussion

### Comparison of the KS-AT arrangement with other structures of isolated KS-AT

There are other structures of KS-AT paired domains from type I PKSs and FASs. Pks13 AT domains are arched ‘downward’ from the KS dimer by some 15° relative to the arrangement seen in type I PKS 6-deoxyerythronolide B synthase (DEBS) 2 module 3^33^, DEBS 3 module 5^19,42–45^, or porcine and human FASs^46–48^ (Fig S21A-D). When compared with the density envelope for type I pikromycin PKS module 5 (PikAIII), docking of KS and AT domains separately into their density shows that the PikAIII AT domains are rotated to arch upwards by ~90° and rotated about their longer axis^49,50^ (Fig S21E). There is a unique specificity for these relative positions perhaps restricted by downstream domains as they may come together around the defining KS dimer two-fold axis.

### The alternate ACP1 positions

Based on the interface analysis using PISA the ACP1a position is consistent with an interface with an ΔG^i^ of −5.5 kcal/Mole. The ACP1b interface though slightly smaller has a similar ΔG^i^. Likewise the number of particles seen on the EM grids is ~3-4 times higher for ACP1a than for ACP1b indicating a very small energy difference between the two states. Applying the free energy relationship between two states b and a at equilibrium ΔG_0_=RTln ([b]/[a]) = ~0.8kcal/Mol difference. Since both ACP1a, and ACP1b are well ordered structures with specific hydrophobic interfaces we suggest that these may be intermediate positions on the pathway to deliver the meromycolate to the KS active site via the delivery conduit. The absence of clear density for the incoming meromycolate on the Ppant of ACP1a suggests that the structure is of an intermediate state following delivery of the R1 α- or α’- chain to the KS active site, and preceding the next FadD32 catalyzed attachment of the acyl-adenylate intermediate to S38 of the ACP1a^12^.

ACP1b is much closer to the delivery conduit in the KS than is ACP1a. The meromycolyl substrates in the KS active site and the lack of the meromycolate density on the S38 of ACP1b suggest that this structure may represent the post-delivery position of the ACP1 retreating from the loaded KS domain. The serine S38 attached to the Ppant arm is removed ~17Å from the position required to deliver the meromycolate (Fig S19). A conserved active site loop in KS is proposed to be a gate that opens to accept substrate ^51^. In Pks13, this loop is in a closed position, consistent with the structure we determine being in a post-delivery state.

### Comparison of ACP:KS binding modes

The KS delivery conduit observed in the Pks13 structure near the ACP1b position is the same tunnel where both trans-acting and cis-acting ACPs bind to their cognate KSs. Trans-acting ACPs from type II PKSs, AntF^52^ and Iga10^53^, and from *E. coli* fatty acid synthase, AcpP^51,54^, have been shown to interact similarly with their cognate KS domains via helix II, with the Ppant-binding serine residue at the N-terminus of helix II pointing towards the substrate delivery conduit (Fig S22A-D). The Pks13 ACP1b is unlikely to adopt this pose to deliver substrate to the KS considering the length constraints of the ACP-KS linking DE-rich linker, since it would entail an approximate 180° rotation about an axis connecting the ACP1b and KS active sites (Fig S22A-D).

During substrate elongation in type I modular PKSs DEBS module 1^55^ and Lsd14^56^, cis-acting ACPs bind near the delivery conduit at the KS dimer interface in a cleft between KS dimer, and KS-AT linker and AT domains of the same protomer (Fig S22E,F). In this position, loop 1 and helix II of ACP interact with the KS dimer, with loop 1 mediating most of the interaction. In both structures, the Ppant arm attached to the ACP stretches into the conduit next to the catalytic C267 of the opposite KS protomer. In Pks13, ACP1b, which resides at the N-terminus of the same polypeptide chain as the KS and delivers the meromycolate chain onto the KS of the same protomer, similarly binds near the cleft between the KS-KS’ dimer, the KS-AT linker’ and AT’ domains (Fig S9, S22E,F). A twist of ACP1b about its connection to the N terminal end of the DE-rich linker would position ACP1 in this cleft with S38 approximately 19 Å, the approximate length of the Ppant arm, from C267 of KS, allowing the Ppant thioester to deliver substrate to the KS active site (Fig S23A). In this docked position, helix III of ACP1 interacts with the KS accepting substrate, and loop 1 interacts with the KS’, KS-AT linker’, and AT’ domains of the opposite protomer. The docked ACP1b orientation in Pks13 is rotated ~90° from the DEBS module 1 ACP binding mode (Fig S23B). The fact that the AT’ domain that is closest to the KS active center is derived from the contralateral polypeptide chain in the Pks13 dimer suggests that the chemistry may alternate between domains of the two protomers of the Pks13 dimer to process a single condensation. Indeed, successive catalytic steps in DEBS PKS have been shown to distribute between protomers^57^.

In the cryoEM structure of pikromycin PKS module 5 (PikAIII)^49,50^, ACP4 from the previous module 4, fused with a flexible linker to the docking domains of module 5, binds on the ‘bottom’ of the KS domain (as viewed in Fig 1C left) between the docking domain helices and the AT domain of the opposite protomer. ACP4 would deliver the growing polyketide from the previous module to the KS active site, hence it is functionally analogous to ACP1 in Pks13, but whereas PikAIII ACP4 operates in trans, ACP1 operates in cis and its position is constrained by the ACP1-KS linker. It was proposed^49^ that the ACP4 serine is next to the side active-site entrance, but in Pks13 this site is sterically occluded by AT. Furthermore, the electrostatic interactions between KS and the DE-rich linker guide ACP1 to the two different sites (i.e. ACP1a and ACP1b) seen in the Pks13 structure and to the docked position of ACP1b at the delivery conduit in KS (Fig 1D, S23).

ACP5 of PikAIII, which transfers the extender unit from the AT active site to the KS active site, binds at the ‘top’ of the KS viewed from the Fig 1C orientation, near the KS lipid conduit^50^. ACP5 has a function analogous to the ACP2 in Pks13. Density for ACP2 is not defined as such in our EM map, but the KS lipid conduit is accessible to the low-resolution lenticular shaped densities we ascribe to the ACP2 and domains C-terminal to it. Thus, ACP2 could deliver the second R2 substrate (C24-C26) through this tunnel. In the Pks13 structure, this KS lipid conduit tunnel is occupied by the meromycolyl acyl chain, therefore it would need to widen to accept the incoming R2 to undergo the condensation with R1 in the KS domain. This mechanism is eminently reasonable, as the product of the condensation must be removed by ACP2 through a topologically accessible pathway, without restriction.

### Bound substrates

Endogenously bound substrates were determined by hydrolysis to release them from the protein, followed by mass spectrometry. The fatty acid chains we identified for R1, R2 are as expected for covalent adducts in the reaction of Pks13 domains, at a particular state in the reaction. The mass associated with each fatty acid chain defines the chemistry without ambiguity. The mass of the R1 alpha-prime C40H78O2 species that is added to C267 indicates an unsaturated C-C double bond with an adjacent methyl substituent^28^. This mass could have also been consistent with a cyclopropane in the aliphatic chain however cyclopropanes are not produced in *M.smegmatis* hence the assignment to the methyl substituent next to the double bond is assured. The “alpha-prime” mycolate is found in *Ms*, (though not in *Mtb)*^21,29^.

The requirement for detergent to release Pks13, we presume from the membrane, raises the question as to where the membrane association is encoded. We surmise that the very hydrophobic meromycolate chains might be chaperoned in part by the lipid bilayer, and so might provide the link that anchors Pks13 to the plasma membrane during mycolic acid synthesis.

### Use of TAM16 as inhibitor of TE

The intent of using a TE inhibitor was to stall the throughput to final product, and thereby to make the sample more homogeneous structurally. The structure reported here is that of the TE-inhibited species that had a marginally better quality. However, the structure was also determined from the non-inhibited sample. Both structures were otherwise indistinguishable, which argues that the substrate bound states of KS and AT are stable states that do not proceed to the Claisen condensation in isolated Pks13 without some other input. Conceivably this could be driven forward by the arrival of new substrates from FadD32, and AccD4 for example.

### Conclusion

A cartoon of the chemical scheme deduced from the structural arrangement is shown in Fig 5. The overall structure of Pks13 is a parallel dimeric assembly of a series of domains that operate in ordered sequence between domains in *cis* or in *trans* to assemble mycolic acids by condensation of two long chain fatty acids. These unusually long chain reactants are moved by carrier proteins (ACPs) between domains of the dimer. Within the dimer individual enzymes are arranged across the parallel two-fold axis, and by their proximities can operate *in trans* between domains on either side of the dimer. The catalytic functions of several isolated domains of PKSs have been described independently at the levels of atomic structure and reaction chemistry ^18,58–61^ though not previously seen in the context of their native environment.

**Fig 5.**
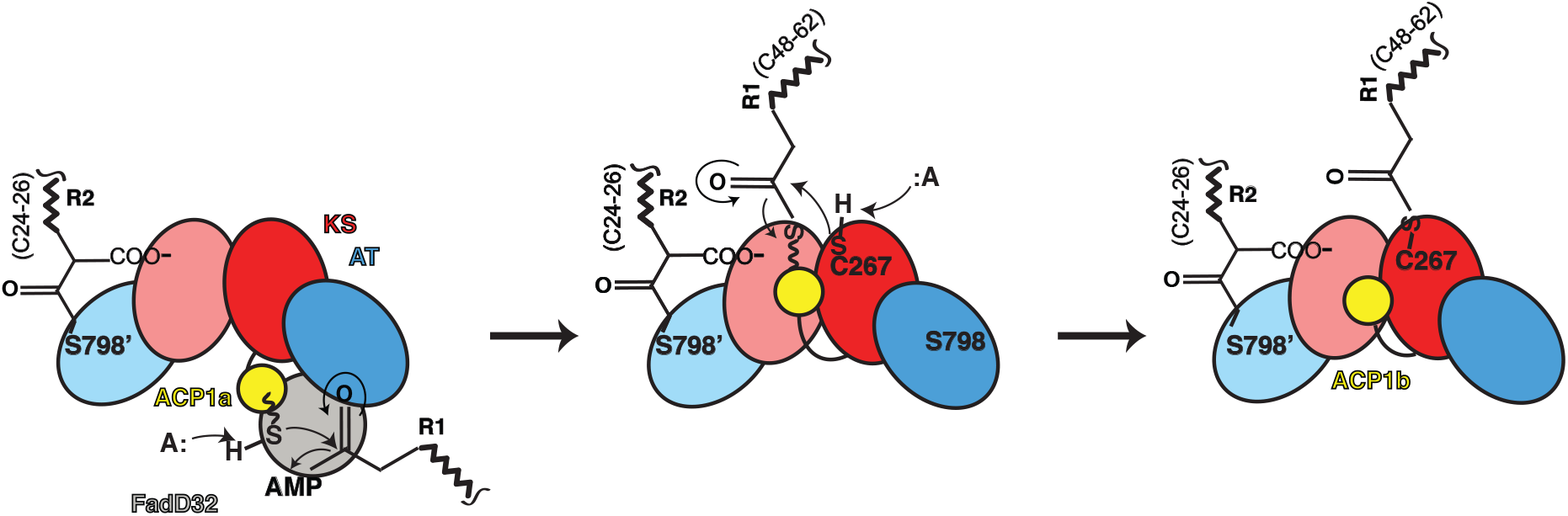
Proposed pathway to the structures determined for substrates bound Pks13. The chemical changes during the mechanism of Pks13 are schematized for just one side of the two-fold symmetric Pks13 for clarity, and incorporating the two structures (ACP1a, ACP1b) we determined. The steps may occur in trans between alternate sides as suggested by the relative proximities of the ACPs and the active sites of KS, and AT of the contralateral protomer. Only the ACP1a, ACP1b, KS, AT domains are cartooned to describe this initial part of the reaction up to the structure we determined (third panel). The left panel shows an early stage of the catalytic cycle in which the R2 fatty acid substrate chain of length (CH_2_)_24-26_ is attached to the AT domain as seen in the structure (Fig 3. The Ppant arm depicted as a wavy line ending in a sulfhydryl and seen on ACP1a (yellow, left panel) links to the adenylated meromycolyl substrate (R1 (CH_2_)_48-62_) from FadD32. ACP1 delivers the attached meromycolyl substrate to the KS active site C267 through trans thioesterification as seen in the structure (right hand panel). This is the stable state reflected in the structure with native substrates bound. In the subsequent steps (not shown, schematized in Fig S1) the carboxy-acyl substrate R2 is transferred to ACP2. Decarboxylative Claisen condensation proceeds between R1 and R2 where the enolate generated by decarboxylation attacks the meromycolyl thioester bound to the KS producing an α-alkyl β-ketothioester. Finally, the TE domain passes the condensed α-alkyl β-ketothioester product to trehalose and is released from Pks13. The product is subsequently reduced by CmrA to form trehalose monomycolate (TMM). TMM is subsequently transported across the plasma membrane by the transporter MmpL3^13^.

Extracting the Pks13 complex from its native context is significant in identifying the *in vivo* states of this protein, as it acts on substrates at their normal concentrations in the mycobacterial cell, rather than a more typical structure generated by adding desired substrate/inhibitor in excess. The substrates covalently bound to the KS and AT were chemically identified and seen liganded to their active sites in density maps. While the entire dimeric structure of Pks13 was imaged by cryoEM and reveals the arrangement of ACP1, KS, AT, and the linking domains between KS and AT, the domains C-terminal to these that include ACP2, ACP3 and TE are resolved in general location only in the EM maps and indicate their necessary hinge bending to fulfill the dynamic aspects of transfer between active centers. Low resolution envelope density and cross-linking mass spectrometry indicates that these domains make large-scale, coordinated motions about the ACP1-KS-AT core.

Mycolic acids are synthesized from very long-chain hydrophobic intermediates, that depend on a chaperoned pathway, in which Pks13 provides a means to control and react on very long chain hydrophobic molecules. In mycobacteria, the acyl-AMP ligase FadD32 and carboxylase AccD4 are encoded within a gene cluster comprised of *fadD32-pks13-accD4*, consistent with their concerted role in the condensation process (Fig 1A). This locus is found exclusively among the mycolic acid producing mycobacteria, and Corynebacterineae with high sequence identity. All three proteins are essential to *Mtb* and therefore are high value drug targets^9,62^.

First-line antimycobacterials against *Mtb* target earlier portions of the mycolic acid synthetic pathway, consistent with their essential role in *Mtb*. Strategies to inhibit the active sites of the ACP1, KS, AT, TE, and the MmpL3 transporter of mycolic acids^63,64^ have been proposed but not yet secured by drugs. Sacchetini *et al*.^17^ using structure-based design, developed a benzofuran class inhibitor of the thioesterase (TE) domain of Pks13, ‘TAM16’, with highly potent *in vitro* bactericidal activity against drug-susceptible and drug-resistant clinical isolates of *Mtb*. The TE domain lies ‘above’ the structure close to the two-fold symmetry axis of Pks13. In isolation, TE domains often form two-fold symmetric dimers^17,58^ consistent with their position around the overall two-fold axis. The structure of the entire dimeric Pks13 adds a new dimension to antimycobacterial drug discovery prospects.

## Acknowledgements

We thank James Sacchettini, Texas A&M University for the inhibitor TAM16 and, Christopher Sassetti, University of Massachusetts Worcester for the Pks13-TEV-eGFP MSMEG strain and both for their insights, Adrian Keatinge-Clay and James Sacchettini for their reading of the manuscript, counsel and advice and Gary Ashley, ProLynx, Inc. for figure S1B. We acknowledge Paul Thomas, Matt Harrington, Joshua Baker-LePain and the Wynton HPC for computational support; and David Bulkley, Zanlin Yu and Glen Gilbert for their maintenance of the UCSF EM Core. We thank Jim Wilkins and Kathy Li for performing initial mass spectrometry for lipid identification in the UCSF mass spectrometry core.

## Funding

Research was supported by P01 AI095208 (Sacchettini), by GM24485 (Stroud) AI48366 (Guan), GM083960 & GM109824 (Sali), U19 A1135990 (Krogan) and R01 AI128214 (Rosenberg). MSD acknowledges an NSF graduate fellowship.

## Author contributions

OR, SK, and RMS conceived the project, SK and JC prepared the protein, MSD prepared EM grids, performed EM imaging and image processing. MSD interpreted density maps and determined the five structures reported. SK, MSD and JFM refined and interpreted the structures. ZG determined the lipid chains attached to PKS13 by mass spectrometry at ‘the lipidomics project’, EHP RK and NK determined the DSSO cross-links formed by mass spectrometry. IE and AS performed integrative structure determination. RMS and OR supervised the project. SK, MSD, JFM, ZG, OR and RMS wrote the manuscript with contributions from all authors.

## Competing interests

The Krogan Laboratory has received research support from Vir Biotechnology, F. Hoffmann-La Roche, and Rezo Therapeutics. Nevan Krogan has financially compensated consulting agreements with the Icahn School of Medicine at Mount Sinai, New York, Maze Therapeutics, Interline Therapeutics, Rezo Therapeutics, GEn1E Lifesciences, Inc. and Twist Bioscience Corp. He is on the Board of Directors of Rezo Therapeutics and is a shareholder in Tenaya Therapeutics, Maze Therapeutics, Rezo Therapeutics, and Interline Therapeutics.

## Data and Materials

The atomic coordinates for five structures of *pks13* have been deposited in the Protein Data Bank with the accession code 7UK4, 8CUY, 8CV1, 8CUZ, 8CV0. The corresponding maps have been deposited in the Electron Microscopy Data Bank with the accession code EMD-26574, EMD-27002, EMD-27005, EMD-27003, EMD-27004.

