## Supplementary Information for "Structure and dynamics of the essential endogenous mycobacterial polyketide synthase Pks13"

**This PDF file includes:**

Materials and Methods

Figures S1-S23

Tables S1-S9

Caption for Movie S1

References

**Other Supplementary Materials for this manuscript include the following:**

Movie S1

### Methods

#### Pks13 fusion generation and culturing

The Pks13-eGFP fusion strain was generated by chromosomally tagging the C-terminus of *Pks13* gene with a TEV cleavage site followed by eGFP in the *Ms recBCD*-mutant strain as described previously[16]. The Pks13-eGFP protein was expressed under the native promoter in *Ms* when the culture was grown in 7H9 media supplemented with 1% (v/v) 50% glucose, 1% (v/v) 50% glycerol, and 0.05% (v/v) Tween 80 at 37° C to $OD_{600}$ ~ 2. Cell pellets were washed three times with 1x PBS, frozen, and reduced to particles by cryo-milling[65].

#### Fluorescence size exclusion chromatography

1 g of lysed cell powder (finely ground *M. smegmatis* frozen cell pellet resulting in cell lysis) was resuspended in 12 mL of solubilization buffer (50 mM Tris HCl pH 7.5, 150 mM NaCl), and a Roche cOmplete™ULTRA EDTA-free protease inhibitor cocktail tablet. 1 μl of benzonase (26.4 units $\mu L^{-1}$; EMD Millipore) was added to help de-clump the lysate, which was stirred at 4° C for 1 hr. To seek a mode of solubilization non-denaturing detergents n-dodecyl-β-D-maltoside (β -DDM), glyco-diosgenin (GDN), digitonin, n-decyl-β-D-maltopyranoside (DM) that are often used for structure determination, and denaturing detergents Fos-choline 12 (FC12), Triton X100 that are denaturing but harsher in solubilizing from membranes were separately added to aliquoted lysate to final 1% (w/v) and solubilized at 4°C for 2 hrs. Unsolubilized materials were pelleted at 55 k rpm for 20 min, and samples were filtered with 0.22 μm filter prior to injection onto the column. A Superose 6 Increase 5/150GL column (GE Healthcare) was run at 0.05 ml/min where 40 μL of each detergent-solubilized sample was loaded with the mobile phase (50 mM Tris HCl pH 7.5, 150 mM NaCl, Roche cOmplete™ULTRA EDTA-free protease inhibitor cocktail, 0.15 mM β-DDM). β-DDM solubilized achieved the most optimal solubilization of Pks13, and thus was used for purification of the protein.

**Protein purification for cryoEM studies**

10 g of cryo-milled lysed cell powder (finely ground *M. smegmatis* frozen cell pellet resulting in cell lysis) was resuspended in 100 mL buffer (50 mM Tris pH 7.5, 150 mM NaCl, 1% β-DDM, one tablet of Roche cOmplete™ ULTRA EDTA-free protease inhibitor cocktail). 10 μl of benzonase (26.4 units $\mu L^{-1}$; EMD MilliporeSigma) was added and the solution was stirred at 4°C for 2 hrs to solubilize membrane-tethered Pks13. The lysate containing Pks13 was spun at 100 kg for 30 min at 4° C. Supernatant containing Pks13 was incubated with 5 mL of anti-GFP nanobody beads at 4°C for 1.5 hr. to allow GFP-tagged Pks13 to bind to the anti-GFP nanobody beads. Anti-GFP nanobody beads were prepared by first expressing and purifying the GFP nanobody protein from *E. coli* BL21(DE3) and conjugating to NHS-Activated Sepharose 4 Fast Flow (Cytiva) following the manufacturer's protocol for conjugation. Pks13-GFP bound anti-GFP nanobody beads were washed three times with 30 mL of buffer (50 mM Tris pH 7.5, 150 mM NaCl, 0.2% β-DDM), with three-minute shaking for each wash to remove non-specifically bound protein(s). Washed anti-GFP nanobody beads were resuspended in 18 mL of buffer (50 mM Tris pH 7.5, 150 mM NaCl, 0.2% β-DDM) and incubated with ~1:10 TEV:Pks13 weight ratio of TEV enzyme overnight with gentle shaking at 4° C to cleave off the C-terminal GFP tag from Pks13. TEV-cleaved Pks13 protein sample was concentrated and purified on a Superose 6 increase 10/300GL column (GE Healthcare) with buffer (50 mM Tris pH 7.5, 150 mM NaCl, and Roche cOmplete™ ULTRA EDTA-free protease inhibitor cocktail), at a 0.3 ml/min flow rate (Fig S2). Fractions containing the Pks13 first peak were pooled and concentrated to 1.5 mg/ml (4μM). The TAM16 inhibitor of the terminal thioesterase[17] was prepared as a 50mM stock in DMSO, and added to the 4 μM protein sample to a final 40 μM concentration with the aim of stabilizing and ordering the thioesterase domains. Inhibitor and Pks13 protein were incubated together at 4° C for one hour before cryo-EM grid preparation.

### Cryo-EM sample preparation

Full length pks13 was concentrated to 1.5 mg $mL^{-1}$ for cryo-EM grid preparation. 4.5 μL of sample was applied to freshly glow discharged holey carbon on gold R1.2/1.3 300 mesh Quantifoil grids and blotted for 9 s with Whatman 1 filter paper at max humidity and $10^oC$ in a FEI Mark IV Vitrobot, before vitrification in liquid nitrogen-cooled liquid ethane.

### Image acquisition

Grids were loaded onto an FEI Titan Krios G3 (at UCSF) operating at 300 kV, equipped with a K3 BioQuantum imaging system, using a 20 eV energy slit at UCSF. Imaging was performed in nanoprobe mode using a 70 um C2 aperture, with a ~1.3 um parallel illuminated area, without an objective aperture. The nominal EFTEM magnification was 105,000x, resulting in a super-resolution pixel size on the specimen of 0.4175 Å $pix^{-1}$. The dose rate was 8 $e^{-}$ $pix^{-1}$ $s^{-1}$ with a total exposure time of 5.9 s, fractionated into 117 frames, using correlated double sampling. Movies were acquired semi-automatedly with 'SerialEM', using 3x3 hole beam-shift image-shift, over a nominal underfocus range of 0.7 to 1.5 µm. The super-resolution movies were drift corrected and dose weighted using UCSF MotionCor2[66] and twice Fourier binned to a pixel size of 0.835 Å $pix^{-1}$.

### Image processing

7567 dose weighted images were imported into cryoSPARC v2.12[67] and CTF estimation was performed in patches. $4.4x10^6$ particles were picked *ab initio* and extracted using a 386 pixel box and subjected to 2D classification from which $1.2x10^6$ particles were retained. The particle set was then used to calculate an *ab initio* 3D volume which displayed high resolution features and quasi-C2 symmetry. The initial volume, low pass filtered to 30 Å was used as a reference for global angular refinement, without imposition of symmetry. Sequential refinement of beam tilt, defocus, Cs, trefoil and tetrafoil resulted in a nominal 2.0 Å reconstruction by gold standard Fourier shell correlation using a 0.143 criterion. Resolution-limited refinement in cisTEM[68] also produced a 2.0 Å reconstruction.

The same procedure was also repeated with imposition of C2 symmetry resulting in a nominal 1.8 Å reconstruction, and these particle data were exported to Relion 3.0[69] for symmetry expansion-focused classification. The C2 map was filtered and segmented in UCSF Chimera[70,71] for selection of domains to focus on. These maps were used to calculate cosine-edged masks using "relion_mask_create". The particle stack was C2 expanded using "relion_particle_symmetry_expand", and classified against the C2 map, using masks for the ACPs, AT and KS-AT domains, without angular searches, varying the regularisation parameter tau from 8 to 20. The particle subsets that provided the best visual features of each respective domain from each classification were selected and further refined using local angular searches and masking over the region of interest. Half-maps from the refinements were used to calculate density modified maps using DeepEMhancer[72] and phenix.resolve_cryo_em[73,74], in order to assist with model building and interpretation.

**Model building and refinement**

Pks13 KS and AT core domains were built *de novo* using a Phyre2-generated homology model[75] as a guide. ACP1a was built into the density guided by aromatic side chain densities for residues W9, W13, F62, and W90. ACP1b was similarly built into density. The negatively charged DE-rich linker subdomain lies against positive charges of arginines and lysines from the KS domain, constrained by favorable electrostatic interactions. In both ACP1a and ACP1b the negatively charged DE-rich linker lies in a net positively charged groove with electrostatic attraction (Fig 1D, E). Few of these side chains form specific interactions, and the sequences are not conserved though the general charged nature is. The closest of these interactions are between Glu88 in the linker that is salt-bridged to Arg384 in the KS (<3.5Å) in ACP-1, and to Lys388 of KS in ACP1b (Fig 1E).

Model building was performed in Coot[76] using a combination of sharpened and density-modified volumes, but all models were finally refined against the gold-standard output volumes from cryoSPARC and RELION using phenix_real_space_refine[77]. All map and model Figures were prepared using University of California, San Francisco (UCSF) Chimera[70] and ChimeraX[71]. MOLE [78] was used to calculate the tunnels reported in the models.

**Analysis of fatty acids in PKS13 by alkaline hydrolysis and LC/MS**

Chains attached to PKS13 were determined by mass spectrometry. Negative ion electrospray ionization (ESI) mass spectra of fatty acids released by hydrolysis are shown (Fig S14, S15, S16), along with their chemical structures and corresponding molecular formulae of the major species. Fatty acids are primarily detected as the deprotonated $[M-H]^-$ ions, along with lower levels of the chloride adduct $[M+Cl]^-$ ions. Hence the fatty acids are determined in the negative ion mode, being 1 Dalton less than the neutral species as listed in the figures.

To remove the fatty acids from the protein, the PKS13 protein sample was subjected to mild alkaline hydrolysis in 3.8 mL chloroform/methanol /0.4 N KOH (1:2:0.8, v/v) at room temperature for 1 hr. The system was converted to a two-phase Bligh/Dyer mixture consisting of chloroform/methanol/water (2:2:1.8, v/v/v) by adding appropriate volumes of chloroform and water. The lower phase was dried under a stream of nitrogen and stored at -20 °C before further analysis. Lipid analysis by normal phase liquid chromatography coupled with electrospray ionization /mass spectrometry (NPLC-ESI/MS) was performed as described[79] using an Agilent 1200 Quaternary LC system (Santa Clara, CA) coupled to a high resolution TripleTOF5600 mass spectrometer (Sciex, Framingham, MA). An Ascentis® Si HPLC column (5 μm, 25 cm × 2.1 mm, Sigma-Aldrich) was used. Mobile phase A consisted of chloroform/methanol/aqueous ammonium hydroxide (800:195:5, v/v/v). Mobile phase B consisted of chloroform/methanol/water/ aqueous ammonium hydroxide (600:340:50:5, v/v/v/v.). The mobile phase C consisted of chloroform/methanol/water/aqueous ammonium hydroxide (450:450:95:5, v/v/v/v). The elution program was as follows: 100% mobile phase A was held isocratically for 2 min and then linearly increased to 100% mobile phase B for 14 min and held at 100% B for 11 min. The LC gradient was then changed to 100% mobile phase C for 3 min and held at 100% C for 3 min, and finally returned to 100% A over 0.5 min and held at 100% A for 5 min. Instrumental settings for negative ion ESI and MS/MS analysis of lipid species were as follows: ion spray voltage (IS) = -4500 V; current gas (CUR) = 20 psi (pressure); gas-1 (GS1) = 20 psi; de-clustering potential (DP) = -55 V; and focusing potential (FP) = -150 V. The MS/MS analysis used nitrogen as the collision gas. Data acquisition and analysis were performed using the Analyst TF1.5 software (Sciex, Framingham, MA).

Based on exact mass measurement, three fatty acid series were identified, 1) C51-C57 long (Fig S14); 2) C36-C42 in length (Fig S15); 3) C22-C26 length (Fig S16). 1) The C51-C57 series are consistent with $\alpha$2-meromycolate as the substrate for which the proximal portion is seen attached to the active site cysteine C267 of KS in the structure (Fig 2). It could theoretically also have been consistent with being released from attachment to the ACP1 Ppant extension prior to trans thioesterification and delivery to the KS. However, no modification is seen in the ACP1 structure arguing against this possibility. Therefore, we conclude that it is the species attached to the KS, hence the system has been captured prior to the Claisen condensation. 2) A very long chain (C36-C42) with a methyl branch followed by a trans double bond in the chain is consistent with the "alpha prime" mycolate found in *Ms* but not in *Mtb*[21]. This chain also is derived from the KS active center. 3) C22-26 fatty acids are fully saturated and most consistent with the $\alpha$-branch species seen in density attached to the AT active center at serine S798. An additional possibility is that some of this species might also have been removed from the ACP2 prior to the Claisen condensation though the ACP2 is not defined in the PKS13 structure. The mass spectra of C22-26 fatty acids do not contain the expected alpha-carboxyl group that is seen in the structure. The carboxyl may have been unstable at some part of the procedure hydrolysis of during mass spectrometry.

**Analysis of Pks13 complexes by in solution DSSO cross-linking and multi-stage mass spectrometry (XL-MS$^3$)**

Pks13 protein complexes were purified from *M. smegmatis* the same way as for cryoEM sample preparation except for the TAM16 addition. Purified Pks13 complexes were cross-linked with increasing amounts of DSSO solubilized in anhydrous DMSO (Table S8). The first replicate was prepared in duplicate, with both replicates cross-linked for 30 min at 1000 RPM, one at 37°C and one at 4°C, with the following molar ratios of DSSO to complex: 1:10; 1:50; 1:250; and 1:1000. Based on the similarity between DSSO cross-linking at 37°C and one at 4°C for Pks13, replicate 3 and 4 were cross-linked for 30 min at 1000 RPM at 37°C only with the following molar ratios: 1:50; 1:250; and 1:1000. Cross-linking reactions were quenched with 50mM Tris pH 8, and then mixed with 4x SDS-PAGE loading buffer to a final 1x concentration (1x SDS-PAGE loading buffer: 62.5mM Tris pH 6.8, 2% SDS, 75mM DTT, 7.5% Glycerol, and 0.02% Bromophenol Blue). Protein samples were then separated by SDS-PAGE on 4-20% Criterion TGX gels (BioRad), stained by AcquaStain (Bulldog Bio) MS safe blue protein stain, and bands excised for in gel digest (Fig S17). Gel pieces were cut to 1mm$^2$ cubes, dehydrated, rehydrated in 15mM TCEP, 25mM $NH_4HCO_3$ reducing buffer, and alkylated in 50mM chloroacetamide in the dark. Gel pieces were dehydrated and rehydrated in 0.5ng/uL trypsin buffer for protein digest. Peptides were extracted from the gel pieces with 50% acetonitrile (ACN), 5% formic acid (FA), dried, and resolubilized in 3% ACN, 2% FA. Resolubilized peptides were separated using an Easy-nLC 1200 (Thermo Fisher Scientific) on a 75 μm × 30 cm fused silica IntregraFrit capillary column (New Objective) packed in-house with 1.9-μm Reprosil-Pur C18 AQ reverse-phase resin (Dr. Maisch-GmbH), or a 15 cm-long column containing 1.7 μm BEH beads (Waters). Peptides were eluted from Reprosil-Pur C18 AQ columns at 300nL/min using the following linear gradient: 50%–8% B in 5 min, 8%–45% B in 35 min, 45%-100% B in 12 min, 100% B for 5 min, 100%-2% B in 2 min, and 2% B for 2 min (mobile phase buffer A: 100% $H_2O$, 0.1% FA; mobile phase buffer B: 80% ACN, 0.1% FA). Peptides were eluted from BEH columns at 300nL/min using the following linear gradient: 5%–22% B in 40 min, 22%–32% B in 5 min,

32%-100% B in 5 min, and 100% B for 10 min (mobile phase buffer A: 100% $H_2O$, 0.1% FA; mobile phase buffer B: 80% ACN, 0.1% FA). Each sample was analyzed in technical duplicate by two independent $MS^3$ methods on an Orbitrap Fusion Lumos (Thermo Fisher Scientific) operated in positive ion mode. In one method, a single acquisition cycle consisted of 9 scan events: 1) one full MS1 scan in the orbitrap (350–1200 m/z, 60,000 resolution, max injection time of 50 ms); 2) two data-dependent MS2 scans in the orbitrap (30,000 resolution, normalized AGC target at 200%, isolation window 1.6 m/z) with normalized collision energy set at 22% on the top two precursor ions; and 3) four MS3 scans in the ion trap (isolation window 2.5 m/z, standard AGC target, auto max injection time, rapid scan rate) with HCD collision energy set at 30% on the top 4 ions from each MS2 scan. In the second method, a single acquisition cycle was allowed 3 sec and consisted of: 1) full MS1 scan in the orbitrap (350–1200 m/z, 120,000 resolution, max injection time of 100 ms); 2) data-dependent MS2 scans in the orbitrap (30,000 resolution, normalized AGC target at 200%, isolation window 1.6 m/z) with normalized collision energy set at 22% on the top two precursor ions; and 3) four MS3 scans in the ion trap (isolation window 2.5 m/z, standard AGC target, auto max injection time, rapid scan rate) with HCD collision energy set at 30% on the top 4 ions from each MS2 scan. For both MS methods ions with charge state 4 to 8 were sampled for MS2 and dynamically excluded for 20 seconds (tolerance of 10 ppm), and ions with charge state 2 to 6 with precursor ions (5 m/z tolerance) excluded were selected for MS3. Cross-linked peptides were analyzed as described in Kaake et al[34].The proteomics data from each step of the analysis pipeline, including raw files, MS2 and MS3 extracted peak files (from MSConvert (ProteoWizard[80,81])), MS3 search files (from ProteinProspector v 6.2.17), and associated search and filtering parameters files, have been deposited to the ProteomeXchange Consortium via the PRIDE[82]partner repository (dataset identifier TBD). Annotated spectra for all inter-linked, dead-end, and single peptides can be found on the MSViewer[83]application through ProteinProspector (https://msviewer.ucsf.edu/prospector/cgi-bin/msform.cgi?form=msviewer) (search key TBD).

**Integrative structure modeling of the Pks13 dimer**

A structural model of the Pks13 in solution was computed by integrative modeling[35,38,84], based on atomic models of component domains obtained by cryo-EM and AlphaFold2[39,40]prediction as well as 57 unique DSSO cross-links. Integrative modeling proceeded through the standard four stages[34–37,85]: (1) gathering data; (2) representing subunits and translating data into spatial restraints; (3) structural sampling to produce an ensemble of models that satisfies the restraints; and (4) analyzing and validating the ensemble models as well as input information. The modeling protocol was scripted using the Python Modeling Interface package, a library for modeling macromolecular complexes based on our open-source *Integrative Modeling Platform* (IMP) package[41] (https://integrativemodeling.org). Files containing the input data, scripts, and output results are freely available at https://github.com/integrativemodeling/Pks13. The details about the approach are given in Table 9.

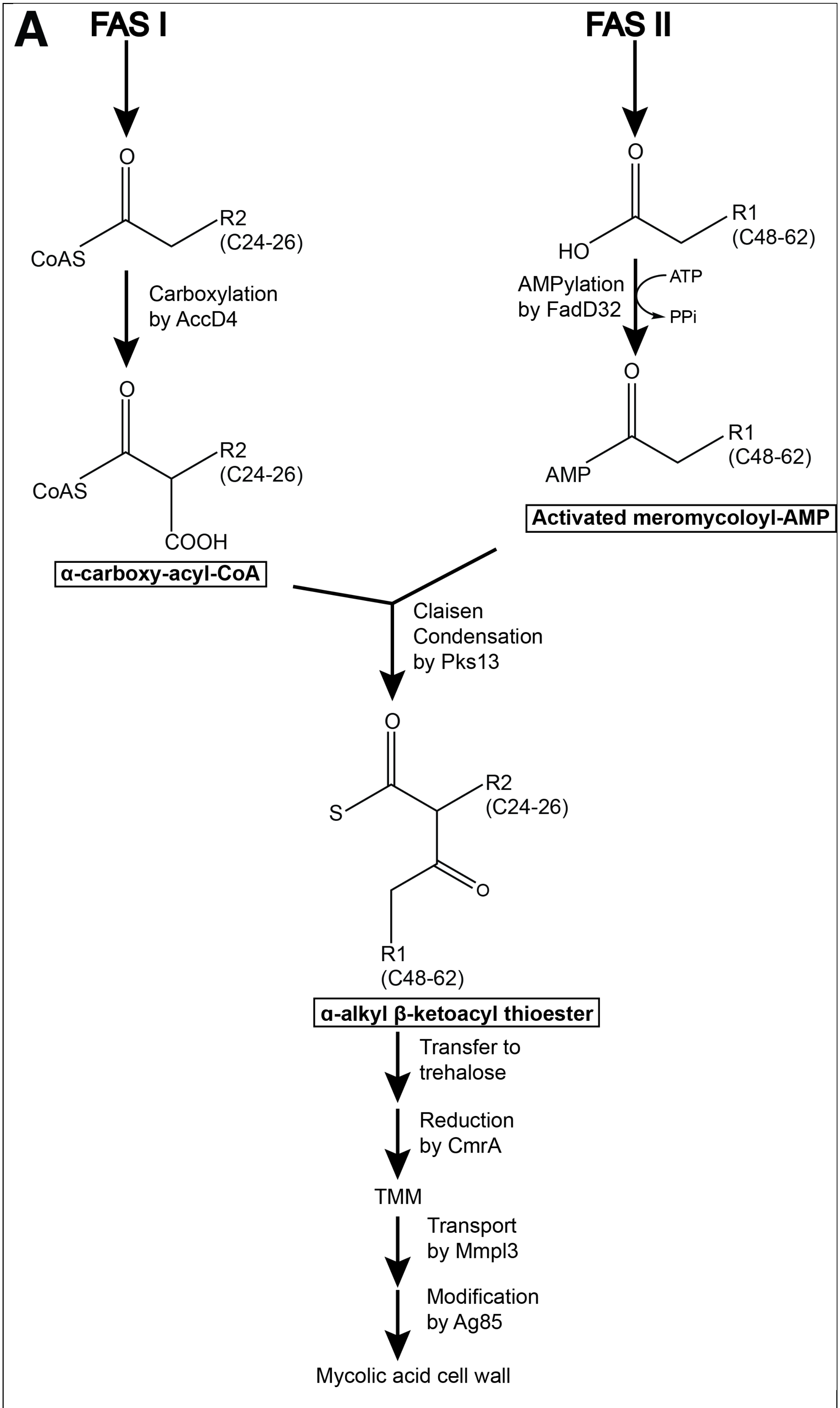
A
FAS I
FAS II
O
CoAS
R2
(C24-26)
Carboxylation
by AccD4
O
CoAS
R2
(C24-26)
COOH
α-carboxy-acyl-CoA
O
HO
R1
(C48-62)
AMPylation
by FadD32
ATP
PPi
O
AMP
R1
(C48-62)
Activated meromycoloyl-AMP
Claisen
Condensation
by Pks13
O
S
R2
(C24-26)
O
R1
(C48-62)
α-alkyl β-ketoacyl thioester
Transfer to
trehalose
Reduction
by CmrA
TMM
Transport
by Mmpl3
Modification
by Ag85
Mycolic acid cell wall

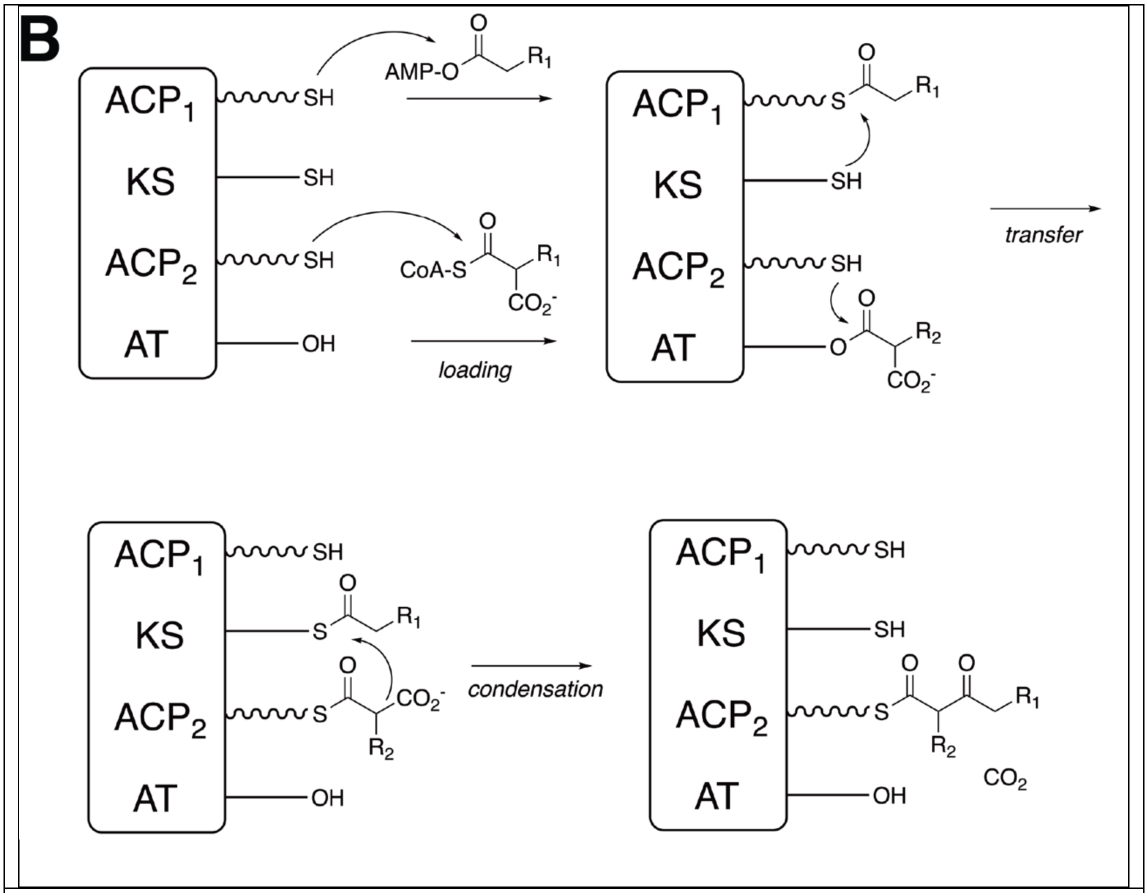

**Fig S1. An Overview of mycolic acid synthesis pathway in mycobacteria.**

(A) FAS I and FAS II synthesize the precursor fatty acids that are condensed by Pks13 to produce α-alkyl β-ketoacyl thioester, a direct precursor of mycolic acids. Long-chain acyl-CoA (C24-C26) is produced by FAS I and carboxylated by AccD4 to produce ɑ-carboxyacyl-CoA. FAS II, composed of multiple enzymes, produces the acyl backbone for the meromycolic chain. The mature C48-C62 meromycolic chain is activated by FadD32 to produce meromycolyl-AMP. Pks13 carries out decarboxylative Claisen condensation of the two fatty acids to produce α-alkyl β-ketoacyl thioester. ɑ-alkyl β-ketoacyl thioester is transferred to trehalose and reduced by CmrA to produce trehalose monomycolate (TMM). The product is transported across the plasma membrane by Mmpl3 and is further modified by the Ag85 complex to form the final building blocks of mycolic acid cell wall. (B) The chemical reactions carried out by the domains are indicated. The reactions can occur in trans between domains of alternate polypeptide chains.

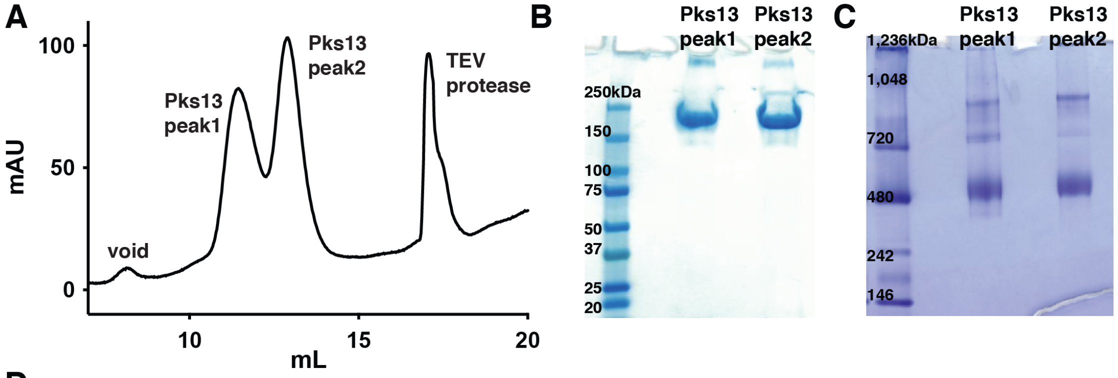

D

```
1    MTVNEMREWL RNWVANATGQ SADAIDESTP MVELGLSSRD AVAMASDIED LTGVTLTATV AFRHPTIESL ATVIIEGEPE
81   PEPYDEDEDW SRTRDVEDIA IVGVATRFPG DLNTPDEMWE ALLEGKDCVT DLPEDRWTEF LDEPRIAERV KKARTRGGYL
161  TDIKGFDSEF FALSKMEADN IDPQQRMALE LTWEALEHAR IPASSLRGES VGVYIGSSTN DYSFLAMSDP SIAHPYAITG
241  TASSIIANRV SYFYDFRGPS VAVDTACSSS LVATHQGVQA LRAGEADVAI VGGVNALVTP LVTVGFDEVG GVLAPDGRIK
321  SFSSDADGYA RSEGGGMLVL KRISDARRDG DQILAVIAGS AVNHDGRSNG LLAPNPDAQA EVLRKAYKDA GINPRDVDYI
401  EAHGTGTILG DPIEADALGR IVGKGRPADK PALLGAVKSN LGHLESAAGA ASLAKMTLAL ANDKLPPSIN YAGPNPYIDF
481  EKERLKVNDT VSDWPRYSGK AIAGVSGFGF GGANAHVVMR EVLAGDLVEP EPEPEPEAKP EKSEADAVYV GGVRMDEYGE
561  FIDEDEPAEG GDAYPSYDED SYELPGITEA AQRLLEQARE ELEAKEAEEP TKQLVPLAVS AFLTSRKRQA AAELADWIDS
641  PEGRASSLES IGRSLSRRNH GRSRAVVLAH DHDEAIKGLR ALAEGKQHPS VLSADGPVTN GPVWVLAGFG AQHRKMGKSL
721  YLRNEVFAEW INKVDALIQD ERGYSILELI LDDNVDYTDA TCEYPIEVVQ LVIFAIQIAL GELLRHHGAK PAAVVGQSLG
801  EAAASYFAGG LSLADATRTI CSRSHLMGEG EAMLFGEYIR LMALVEYSAD EIKTVFSDYP DLEVCVYAAP TQTVIGGPPD
881  QVDAIIARAE SEGKFARKFQ TKGASHTQQM DPLLGELAAE LQGIEPKPLT TGYFSTVHEG TFIRPGSAPI HDVDYWKKGL
961  RHSVYFTQGI RNAVDNGHTT FLELAPNPVA LMQVGLTTAS AGLHDAQLIA TLARKQDEVE SMISAMAQLY VHGHDLDFRT
1041 LFPRRSKGLA GALDFANIPP TRFKRKEHWL PAHFTGDSSA VMPGNHVATP DGRHVWEFVP RGKTDLAALV KAAAAQVLPD
1121 AKLAAFEQRA VPADNARLVT TLTRHPGGAT VQVHARVEES FTLVYDAIVA RANGAGVTAL PVAVGAGVAV SGDVAGEGAG
1201 ASVIEDDEPD AEILQDNLTA GAGMGADFQK WDPNSGETIG QRLGTIVGAA MGYEPEDLPW EVPLIELGLD SLMAVRIKNR
1281 VEYDFDLPPI QLTAVRDANL YNVEELIRYA IEHRDEVEQI AESQKGKTAE EIAAEQSELL GGASTVAELE AKLAEAGHPL
1361 AAKDSEDSEN SEDNAAGAAA AAEASAVEGL EIPPPPTDPT GPGGAPIPPP PSDPSGPAQA ASATDAPAGT VNKATAAAAA
1441 AKVLTQEAVT EALGADVPPR DAAERVTFAT WAIVTGKSPG GIFNELPTVS EETAKKMAER LSERAEGTIT VEDVLGAKTI
1521 EGLATIVREQ LEEGVVDGFV RTLRPPKEGS NAVPLFVFHP AGGSTVVYEP LMKRLPADVP VYGLERVEGS IEERAAEYVP
1601 KLLEMHKGPF VLAGWSLGGA LAYACAIGLK QSGADVRFVG LIDTVLPGEP IDQSKEGMRA RWDRYARFAE RTFNVEIPAI
1681 PYEELEKLDD EGQVKYVLEI VKESGVQIPG GIIEHQRTSY LDNRALDTVD IKPYDGHVTL YMADRYHDDA IVFEPAYATR
1761 KPDGGWGSFV SDLEVVHIGG EHIQAIDEPY IAKVGAHMSE ALNRIEAQAS KEDGAK
```

| Num Unique | % Cov | Best Disc Score | Best Expect Val |
|---|---|---|---|
| 242 | 78.1 | 14.32 | 6.6e-26 |

**Fig S2. Purification of native Pks13 from mycobacteria.** (A) Superose 6 size exclusion chromatography profile of Pks13 following protein extraction with β-DDM from cryo-milled mycobacteria, anti-GFP nanobody beads affinity purification, and on-bead GFP tag cleavage by TEV protease. Two peaks containing Pks13 are labeled 1 and 2, in addition to the void peak and peak containing TEV protease. Pks13 peaks were run on non-reducing SDS-PAGE (Biorad #4561096) (B), and blue native PAGE (C). In SDS-PAGE (B) the protein runs at ~198 kDa, as expected for a protomer of

Pks13. Blue native PAGE (C) indicates that majority of the protein is in a dimeric form, and higher oligomeric forms are present. Upon screening both peaks 1 and 2 on gold Quantifoil grids by cryoEM, peak 1 was selected for data collection since it produced more monodisperse particles that correspond to Pks13 dimers. We conclude that the Pks13 dimers associate loosely in solution, and that these readily dissociate to form monodisperse Pks13 dimers under the conditions in which the images were acquired. (D) 78.1% sequence coverage (highlighted red) by tryptic digested Pks13 fragments run on LC-MS-MS.

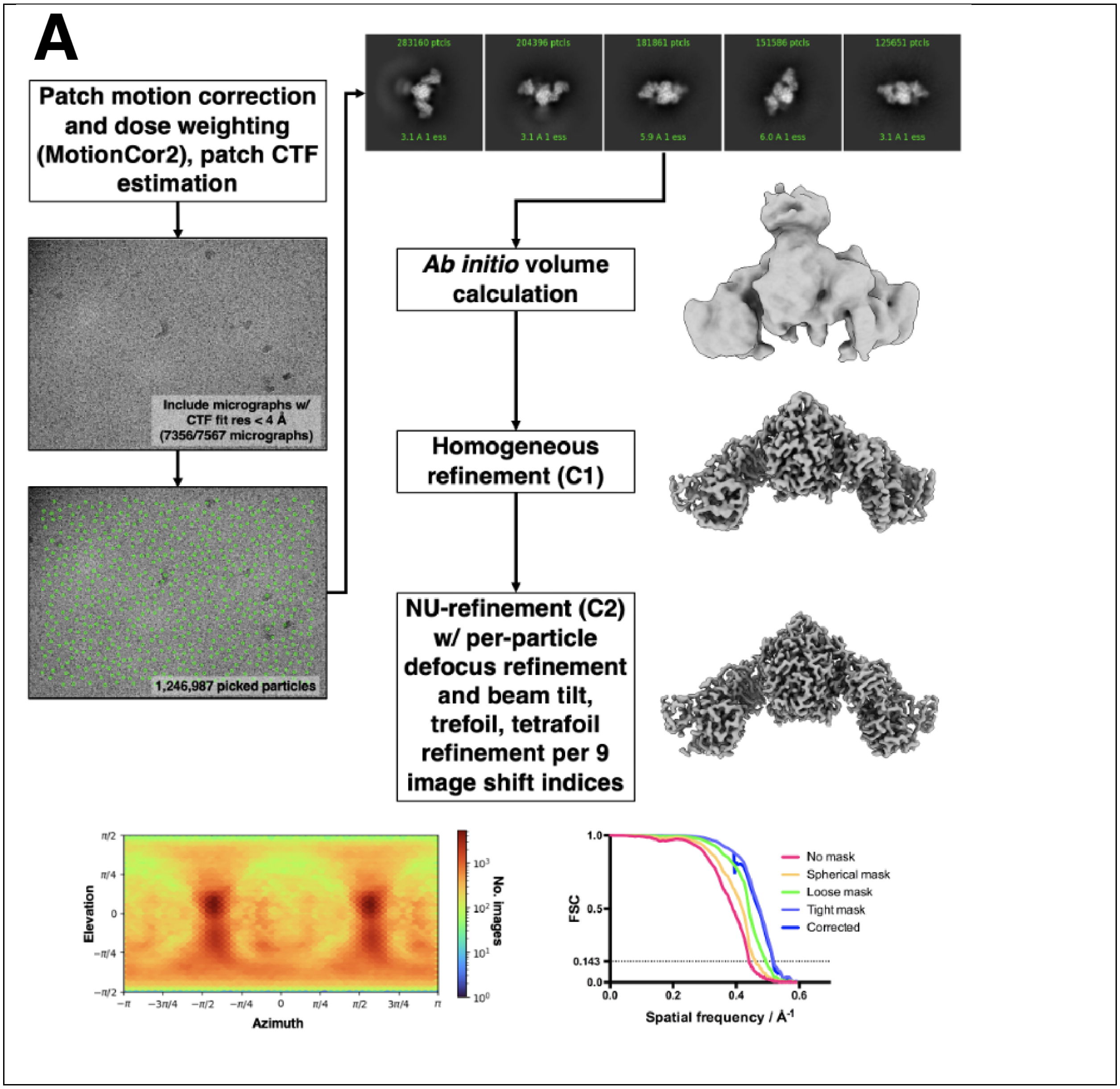
A
Patch motion correction and dose weighting (MotionCor2), patch CTF estimation
Include micrographs w/ CTF fit res < 4 Å (7356/7567 micrographs)
1,246,987 picked particles
283160 ptcls
204396 ptcls
181861 ptcls
151586 ptcls
125651 ptcls
Ab initio volume calculation
Homogeneous refinement (C1)
NU-refinement (C2) w/ per-particle defocus refinement and beam tilt, trefoil, tetrafoil refinement per 9 image shift indices
Elevation
Azimuth
No. images
FSC
Spatial frequency / Å-1
No mask
Spherical mask
Loose mask
Tight mask
Corrected

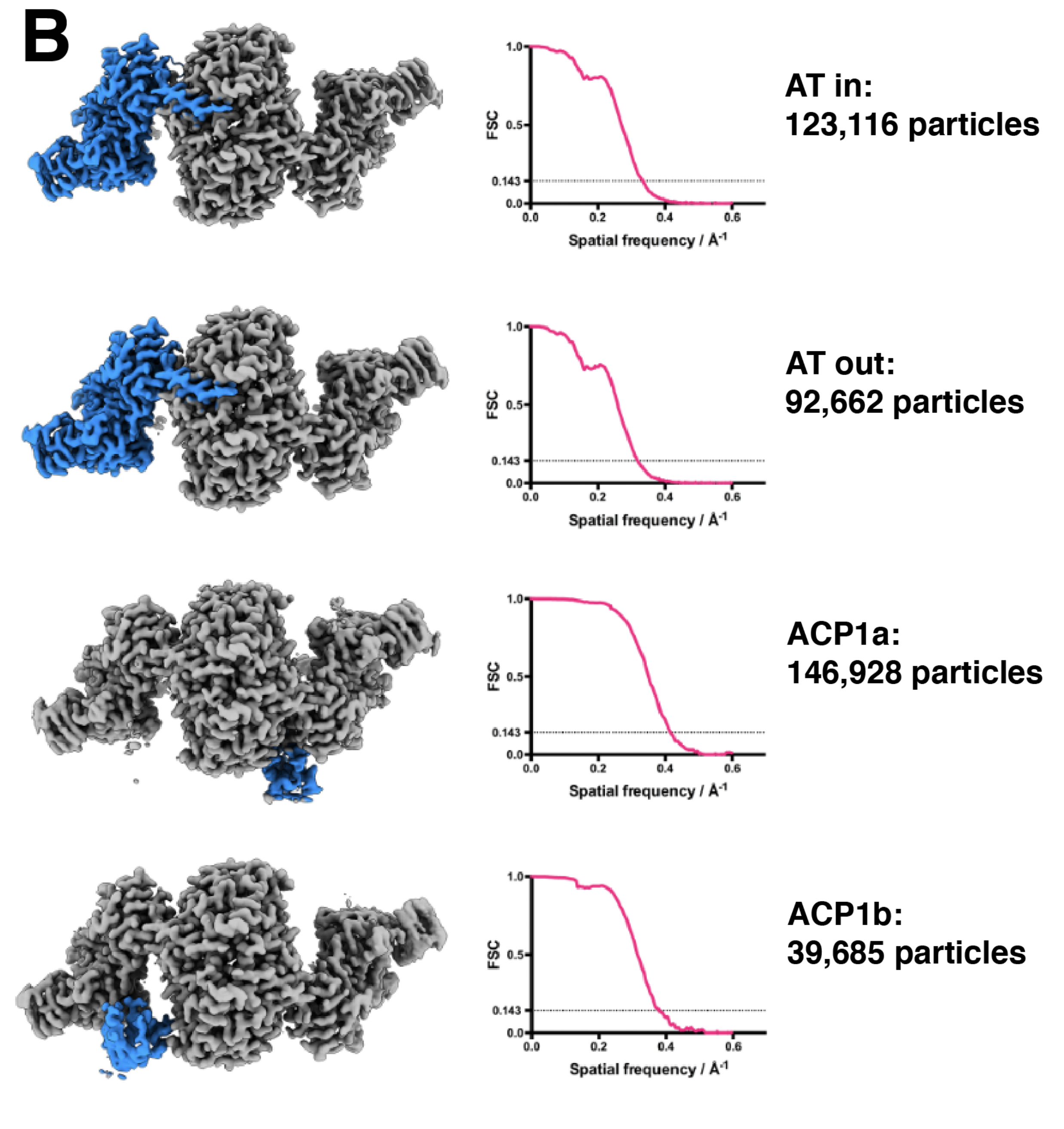

**Fig S3. Single particle analysis processing workflow from movies through consensus C2 refinement and focused refinements.** (A) Image processing workflow done in cryoSPARC for the initial C2 refinement. Particle image pose heat map (displayed as blue to red, low to high numbers of images) showing an isotropic distribution, and complete gold standard (independent half-refinement) Fourier shell correlation plots using the $FSC_{0.143}$

criterion, are included. (B) Symmetry expansion-focused classification done in RELION probing AT and ACP1 domain dynamics. Two conformations of the acyl transferase domain (blue region in both) were identified and used for focused refinement. Two classes were also seen for positions of the ACP1, namely ACP1a (blue), and ACP1b (blue). The number of particles is ~3-4 times higher for ACP1a than for ACP1b indicating a very small energy difference between the two states. The soft masks used for focused classification encompass the blue-colored regions of each map, each dilated ~ 10 Å around the regions of interest. The volumes displayed are viewed down the pseudo-symmetry axis, orthogonal to the viewing direction in (A).

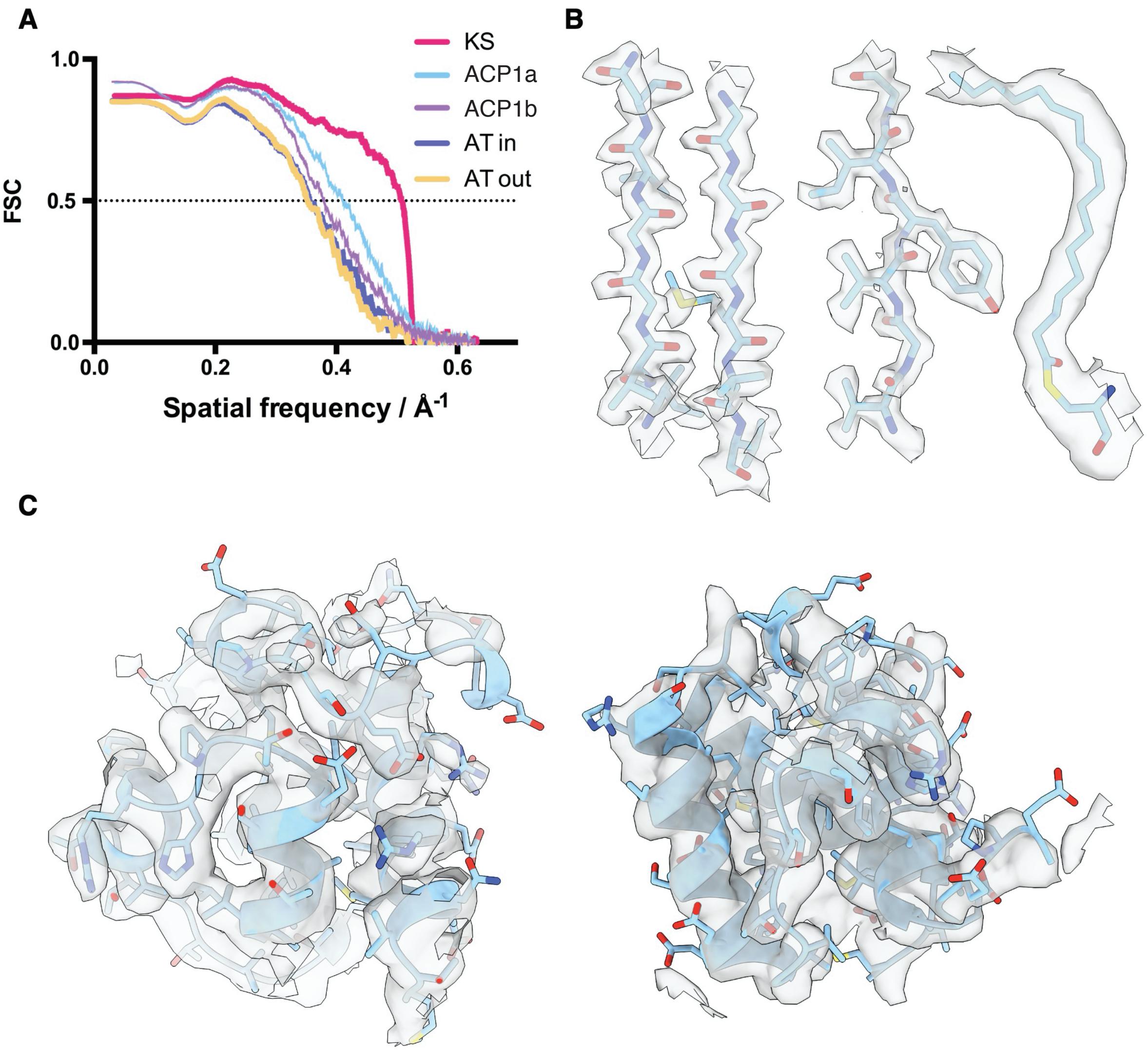

**Fig S4. Map-to-model Fourier shell correlations of focused refinements, and example EM density.** (A) Map-to-model FSC curves for the focused refinements and their cognate models with threshold 0.5. (B) Near-atomic resolution density in the KS-AT core, zoned 2 Å around the model. Two beta strands are shown (left) and lipid density extending upward from the KS active center C267 (right). (C) Density for ACP1a (left) and ACP1b (right), zoned 2 Å around the model.

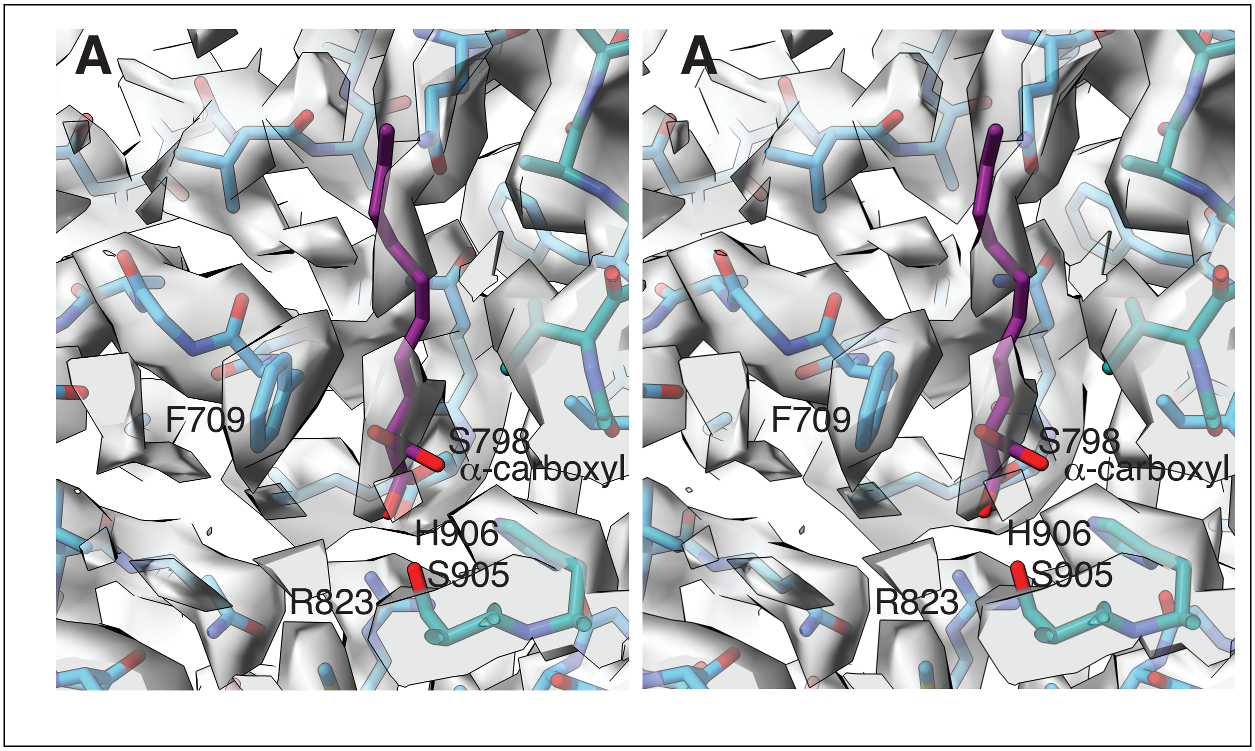
A
F709
S798
α-carboxyl
H906
S905
R823
A
F709
S798
α-carboxyl
H906
S905
R823

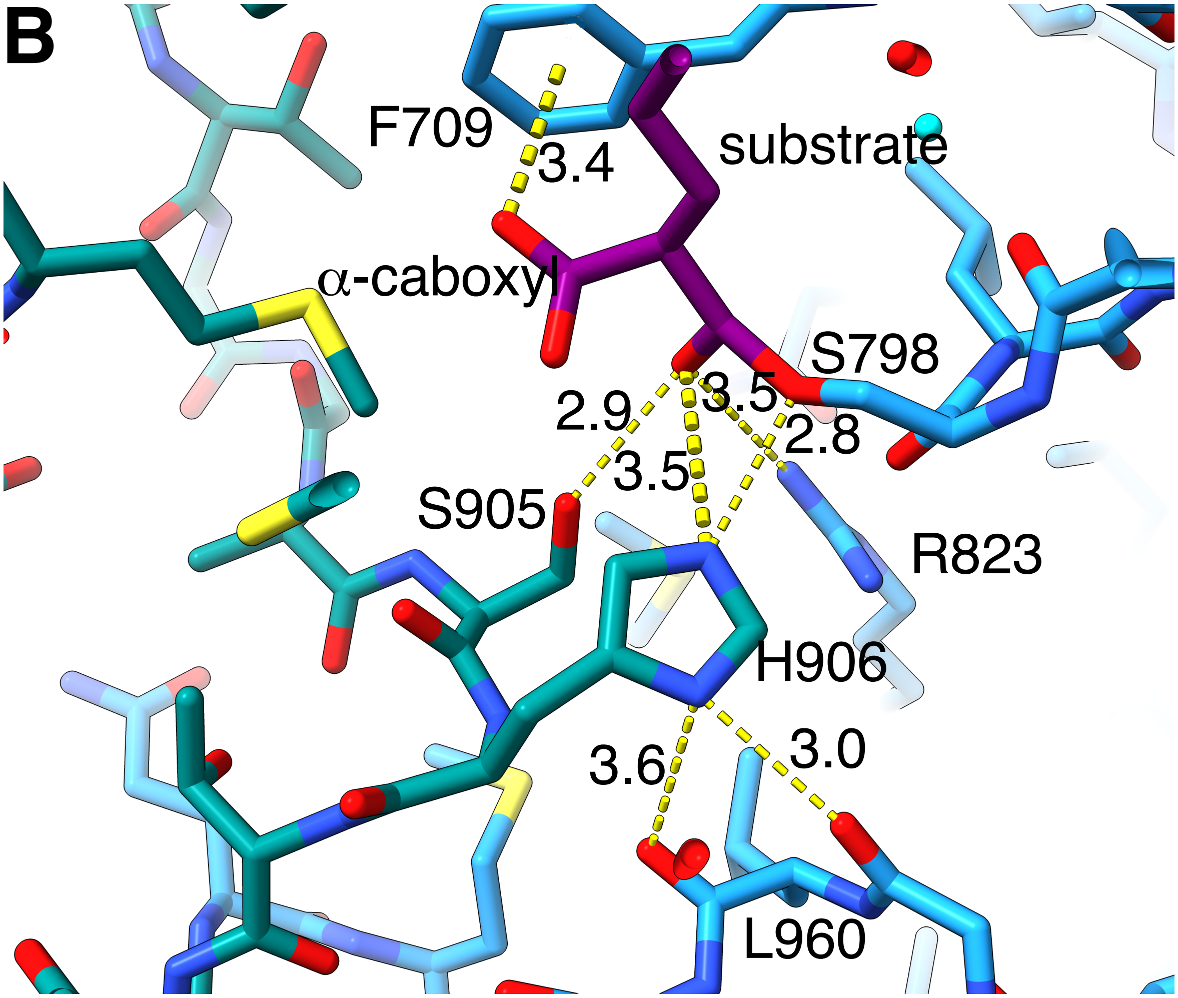

**Fig S5. The active center of the AT domain shows the bound substrate interactions.** (A) (top) Crossed eye stereo image showing the density for the active site of the AT domain with substrate bound. The α-carboxyl group on the bound substrate is stabilized by a π-anion interaction with the phenyl ring of F709. (B) Rotated ~90° around the vertical axis and tilted forward from panel (A) shows interactions between the active center of the AT domain and the bound α-carboxy acyl substrate ($C_{24}$-$C_{26}$) chain attached to S798. Distances between heavy atoms are tabulated in Å. The catalytic base H906 is ideally placed at 2.8Å from the γO of S798. The ester carbonyl on the substrate, and hence the oxyanion intermediate formed at that oxygen during the catalytic reaction is stabilized by R823, H906, S905. The H906 orientation is aligned by hydrogen bonds from the δN-H to the C=O of G959 and of L960.

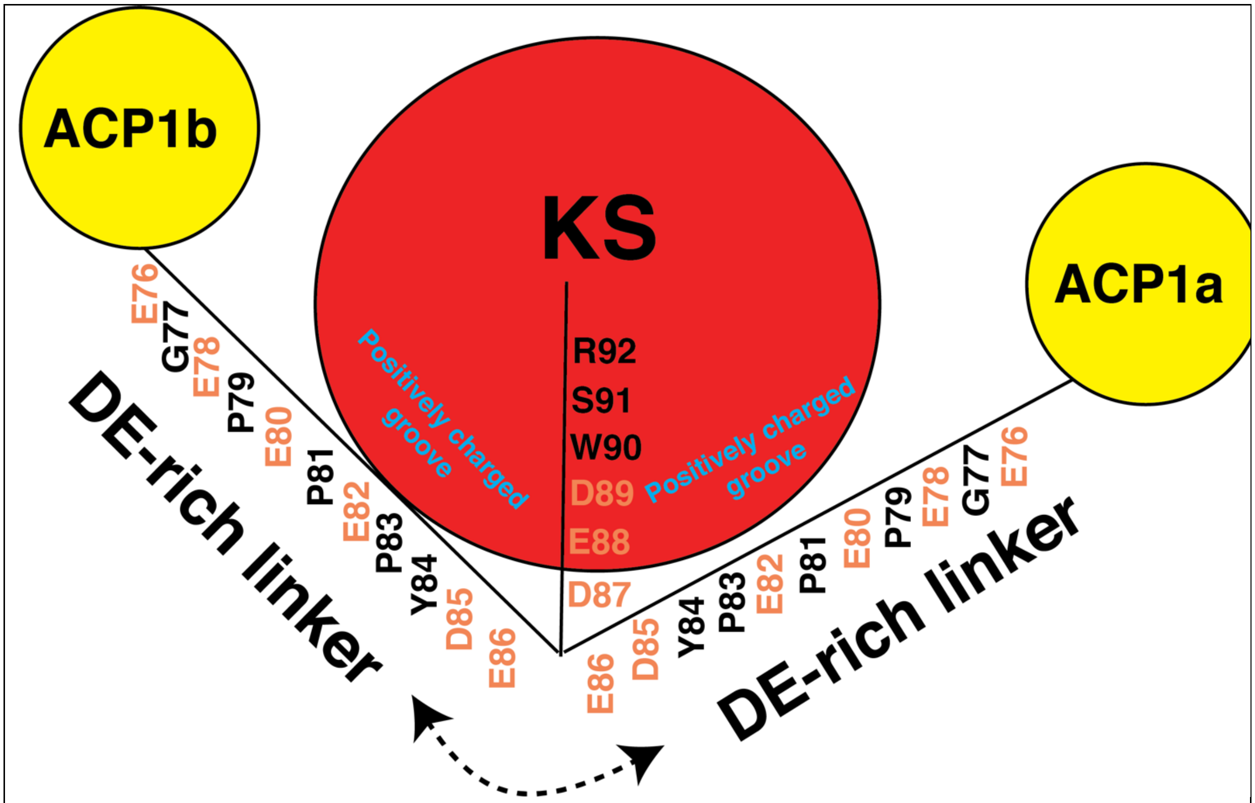

**Fig S6. DE-rich linker.** 17 amino acid linking sequence (E76-92) between ACP1 and KS includes nine acidic amino acids (colored orange) and termed the “DE-rich linker”. First 11 amino acids (E76-E86) occupies either of two positively charged groves on the KS surface.

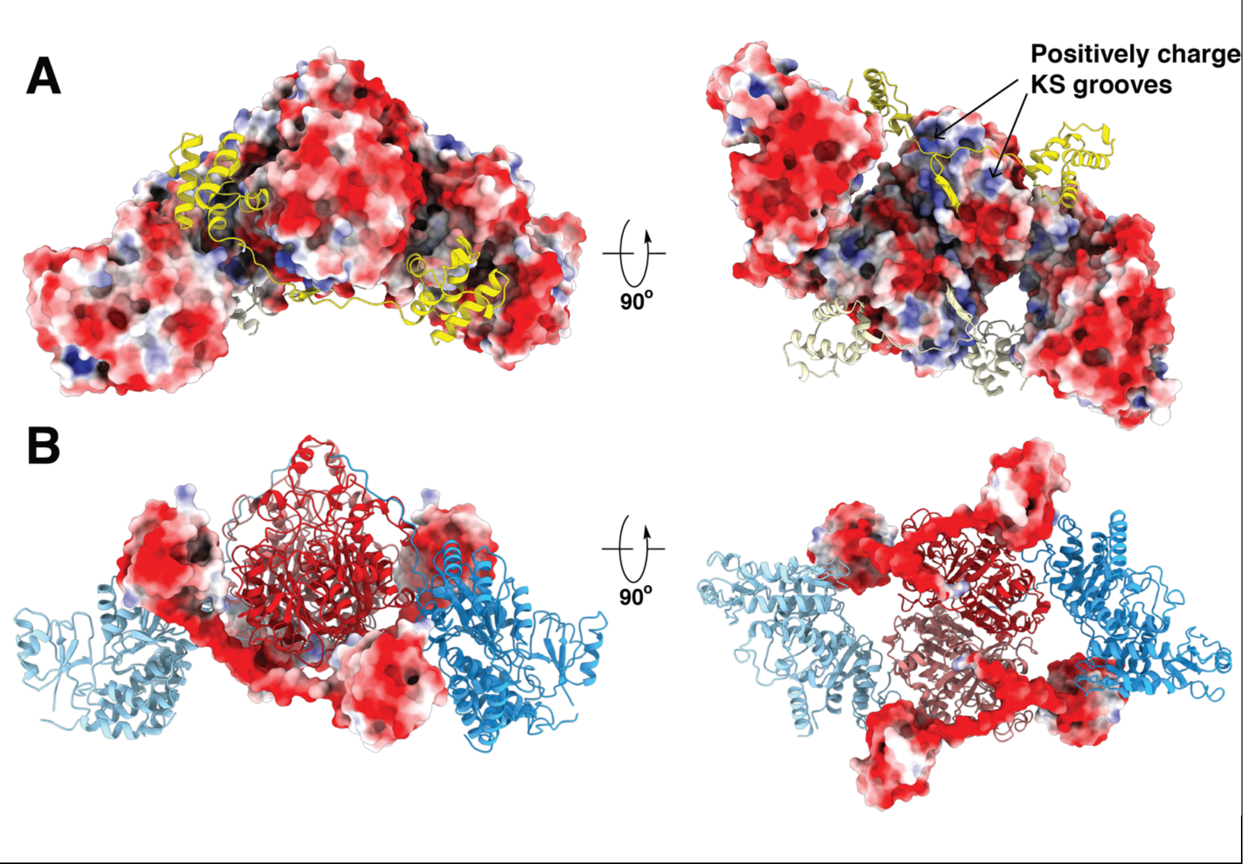

**Fig S7. Electrostatic surface rendering of ACP-KS-AT domains.** (A) The KS-AT surface is colored by electrostatic potential (red: acidic, blue: basic) of the residues. ACP1a and 2 and their ensuing linkers to the KS (yellow and lemon chiffon) are superimposed onto the KS-AT electrostatic potential surface, illustrating the NACP-to-KS linker that docks into either of the two positively-charged grooves on the KS. (B) ACP1a, 2, and the linkers to the KS are colored by electrostatic potential (red: acidic, blue: basic) of the residues. Model of KS-AT are superimposed onto the highly negative electrostatic potential surface.

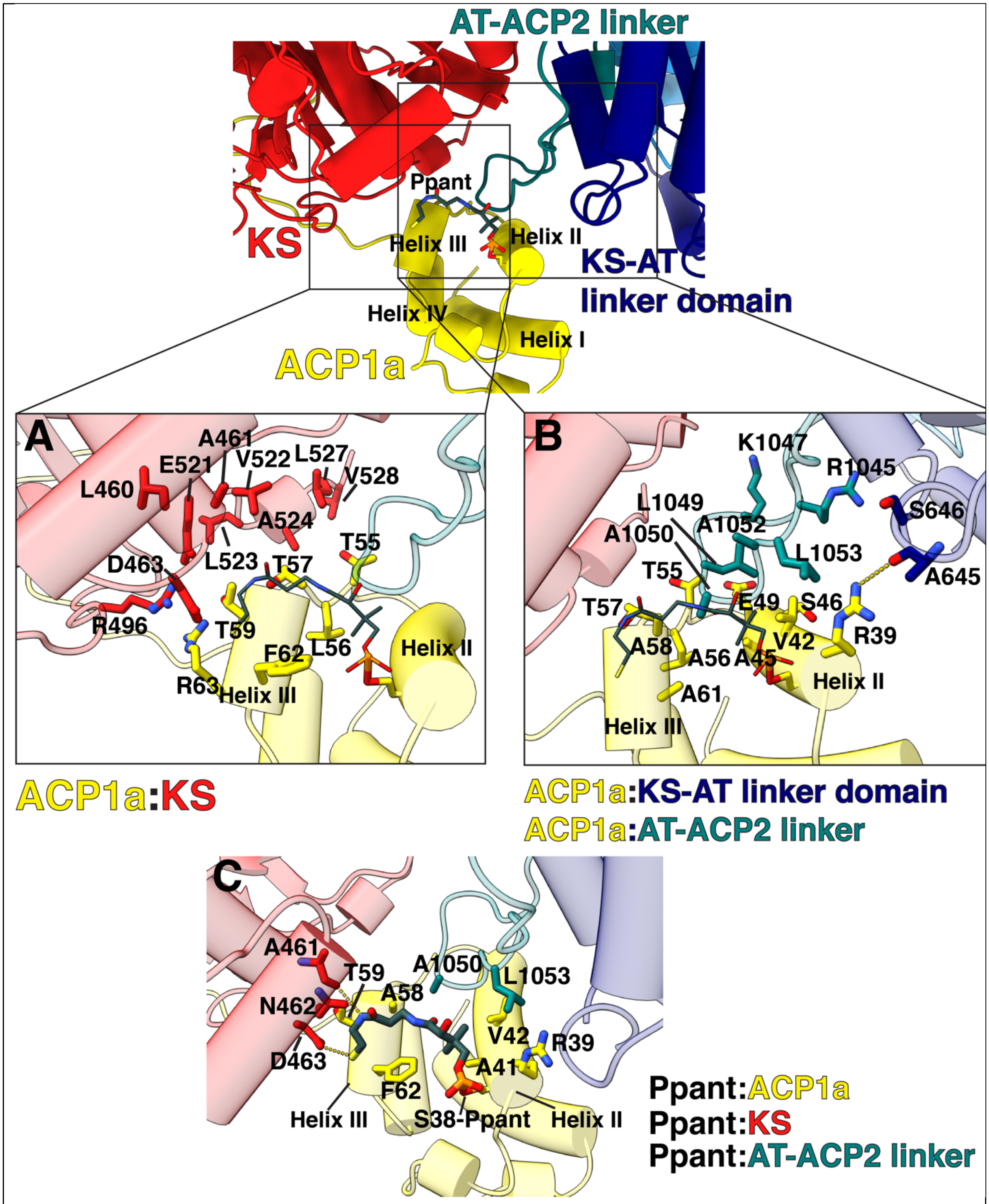

**Fig S8. Holo-ACP1a interaction with KS, KS-AT linker domain, and AT-ACP2 linker.** (A)

Interfacing residues between holo-ACP1a and KS are shown as stick representation and labeled. Ppant arm is shown coming off of ACP1a S38 in helix II. Pi-pi stacking interaction is shown between R63 of ACP1a and R496 of KS. (B) Interfacing residues between holo-ACP1a with KS-AT linker domain and with AT-ACP2 linker are shown. Hydrogen bond between R39 NH and A645 backbone oxygen is labeled with yellow dashed line. (C) Ppant interaction with residues from ACP1a, KS, and AT-ACP2 linker are shown. Two hydrogen bonds formed between Ppant N41 and KS A461 backbone oxygen and Ppant S44 and KS D463 OD1 are labeled with yellow dotted lines.

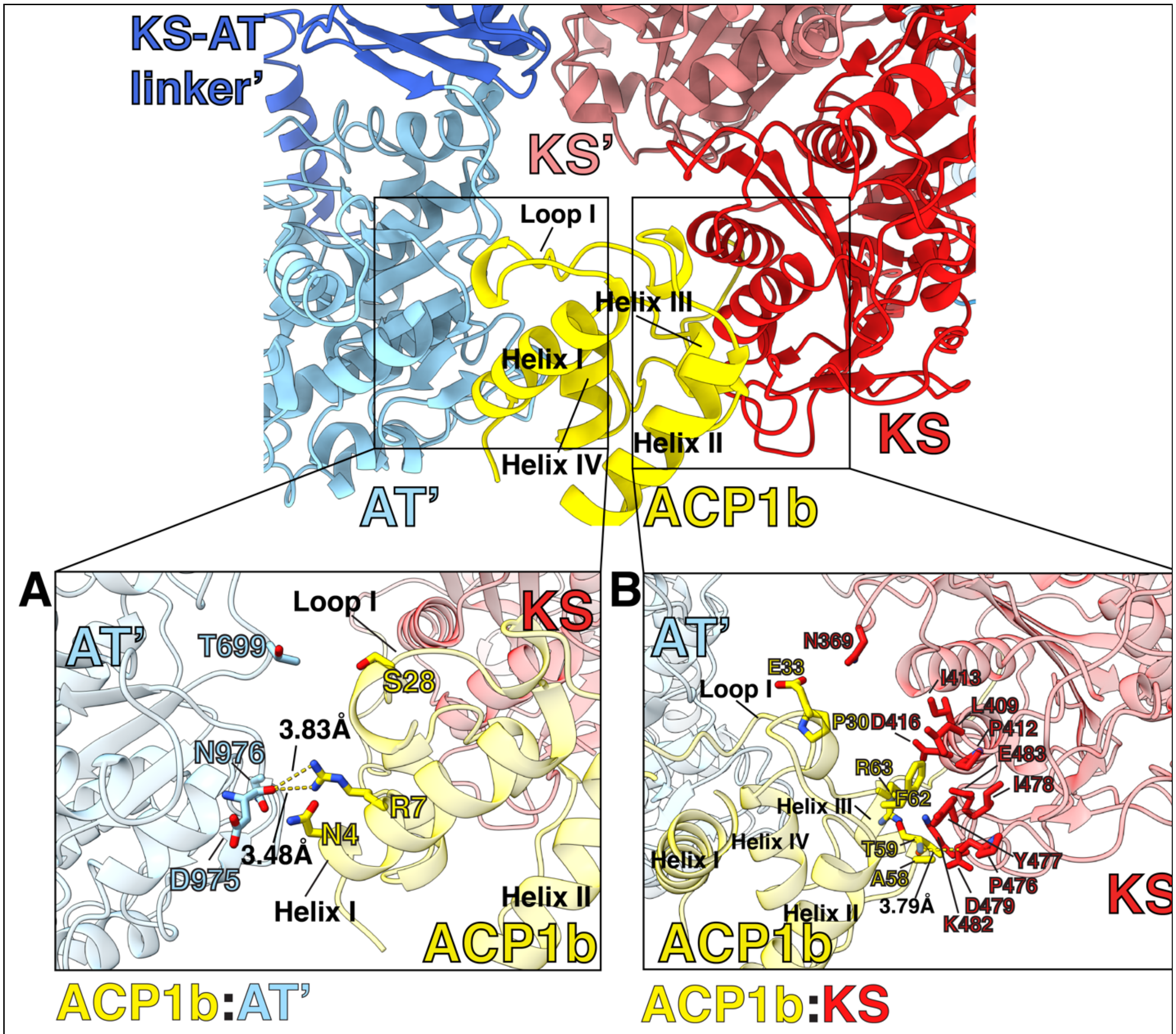

**Fig S9. ACP1b interaction with KS and AT'.** (A) ACP1b:AT' interacting residues are shown and labeled, which includes two hydrogen bonds as indicated with yellow dotted lines between the KS R7 guanidinium group and the backbone carbonyl oxygen of AT' D975. (B) ACP1b:KS interacting residues are shown and labeled, which includes a single hydrogen bond between the backbone amide of T59 and the carbonyl oxygen of P476.

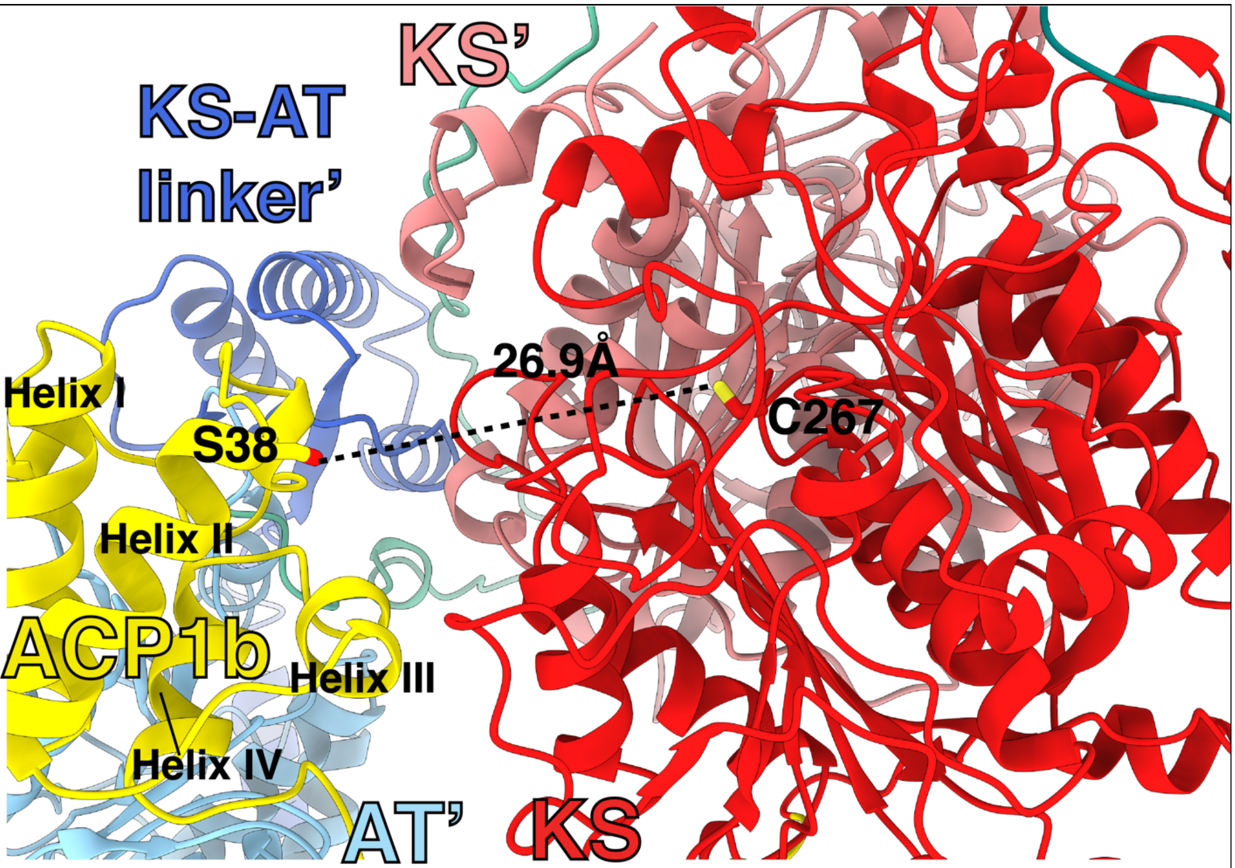

**Fig S10. Distance between ACP1b S38 and KS C267.** ACP1b S38, which received the Ppant arm, is 27 Å away from the KS active site C267 of the same protomer.

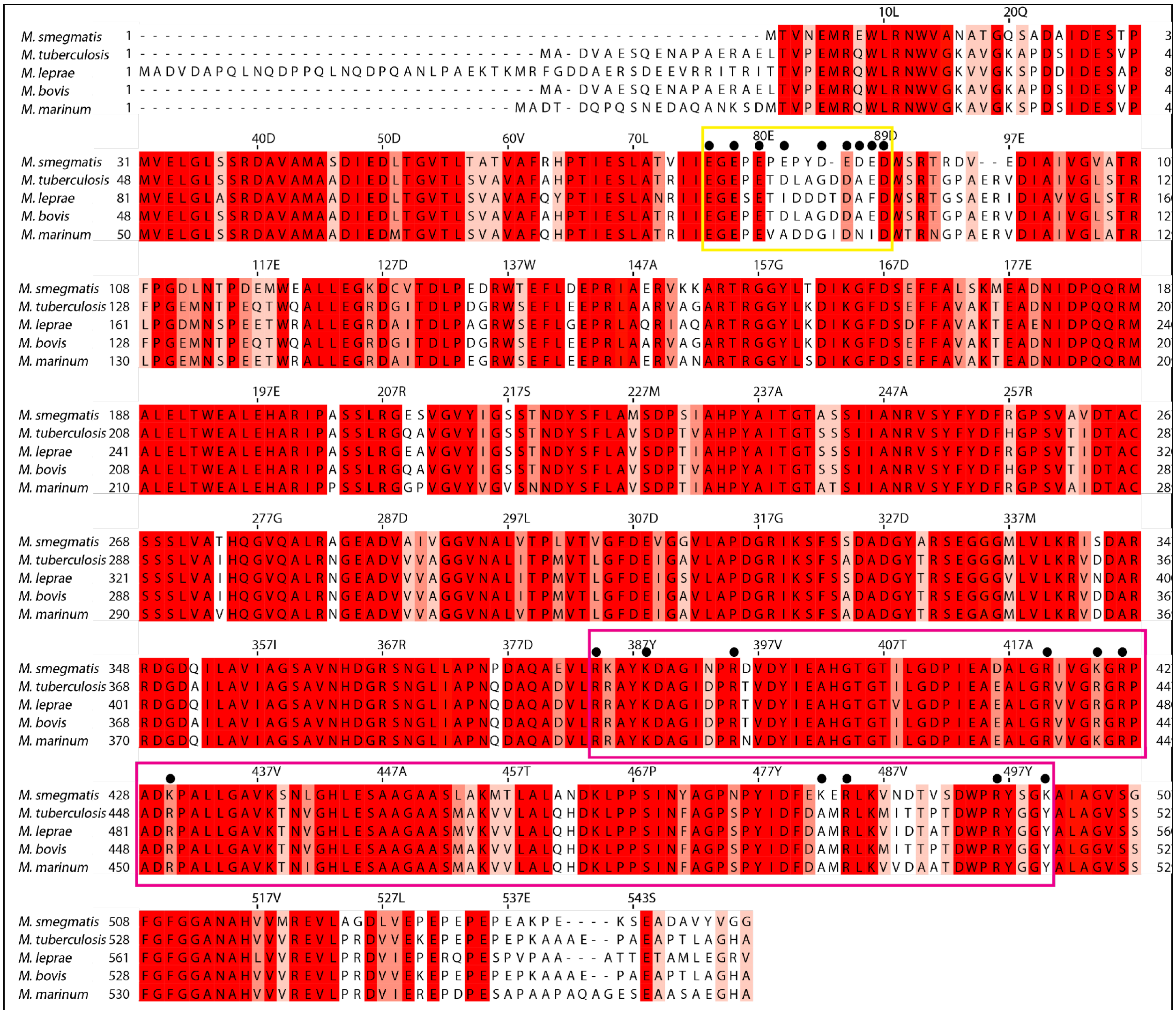

**Fig S11. Sequence alignment of Pks13 ACP1 – KS 'DE-rich linker' region across mycobacterial species.** The DE-rich linker region is boxed in yellow. The corresponding interacting basic region in the KS is boxed in magenta. Pks13 protein sequences from five different mycobacterial species are aligned from N-terminus to the end of the KS domain. Sequence numbering above the sequences is for *Ms.* Residues were colored by conservation with highly conserved residues in dark red and less conserved residues in lighter colors. Acidic and basic residues identified from the structure to make general electrostatic interactions (shown in Fig 1C,D,E) are labeled with black dots. Alignment was carried out using Clustal Omega[86] and annotations were made in Jalview[87] . This interaction is mediated by 773.6Å$^2$ buried surface area. Distance between N-ACP1 S38 which attaches to Ppant, and the KS catalytic residue C267 of the same protomer is 27Å.

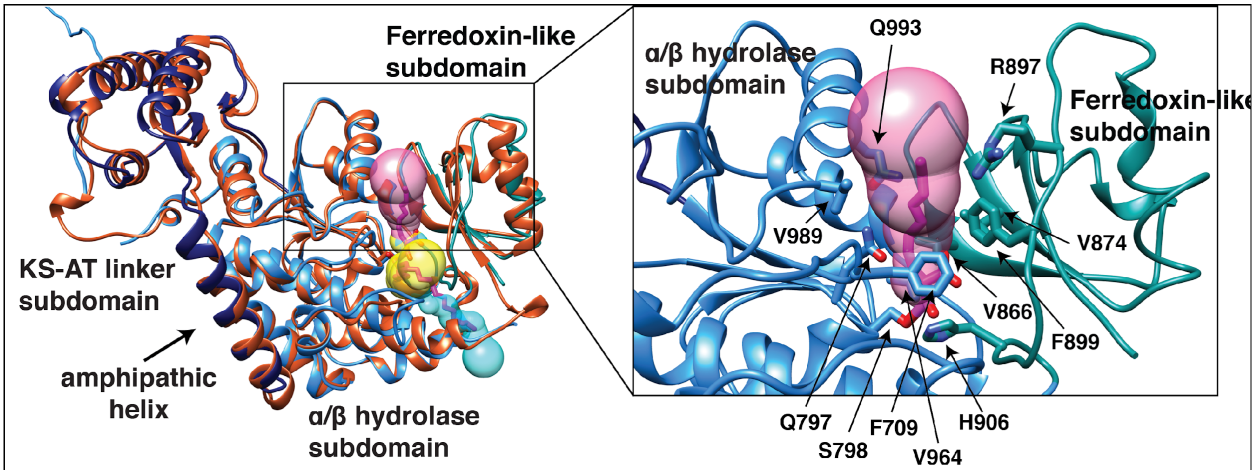

**Fig S12. Comparison of carboxyacyl substrate positions between *Mtb* and *Ms* Pks13 AT domains.** Left: *Ms* Pks13 AT structure (KS-AT linker subdomain; navy blue, αβ-hydrolase subdomain; blue, and ferredoxin-like subdomain; cyan) is overlaid with *Mtb* Pks13 AT fragment structure (PDB:3TZZ[25], orange all throughout), resulting in 1.00 Å RMSD over 449 Cα atoms. The carboxyacyl ligand in two different positions as reported in *Mtb* Pks13 AT fragment[25] lies within the yellow and cyan hydrophobic tunnels. The native carboxyacyl substrate in our *Ms* Pks13 AT structure is seen in a different hydrophobic tunnel colored magenta. In all three cases, the serine-ester carboxyacyl substrate is colored magenta. Inset: Close-up view of the hydrophobic tunnel surrounding the native substrate in *Ms* Pks13 AT. Residues lining the tunnel are labeled. Tunnels were calculated using MOLEonline[78]. The amphipathic helix unique to mycobacterial Pks13 that leads into the KS-AT linker subdomain is labeled in the left panel.

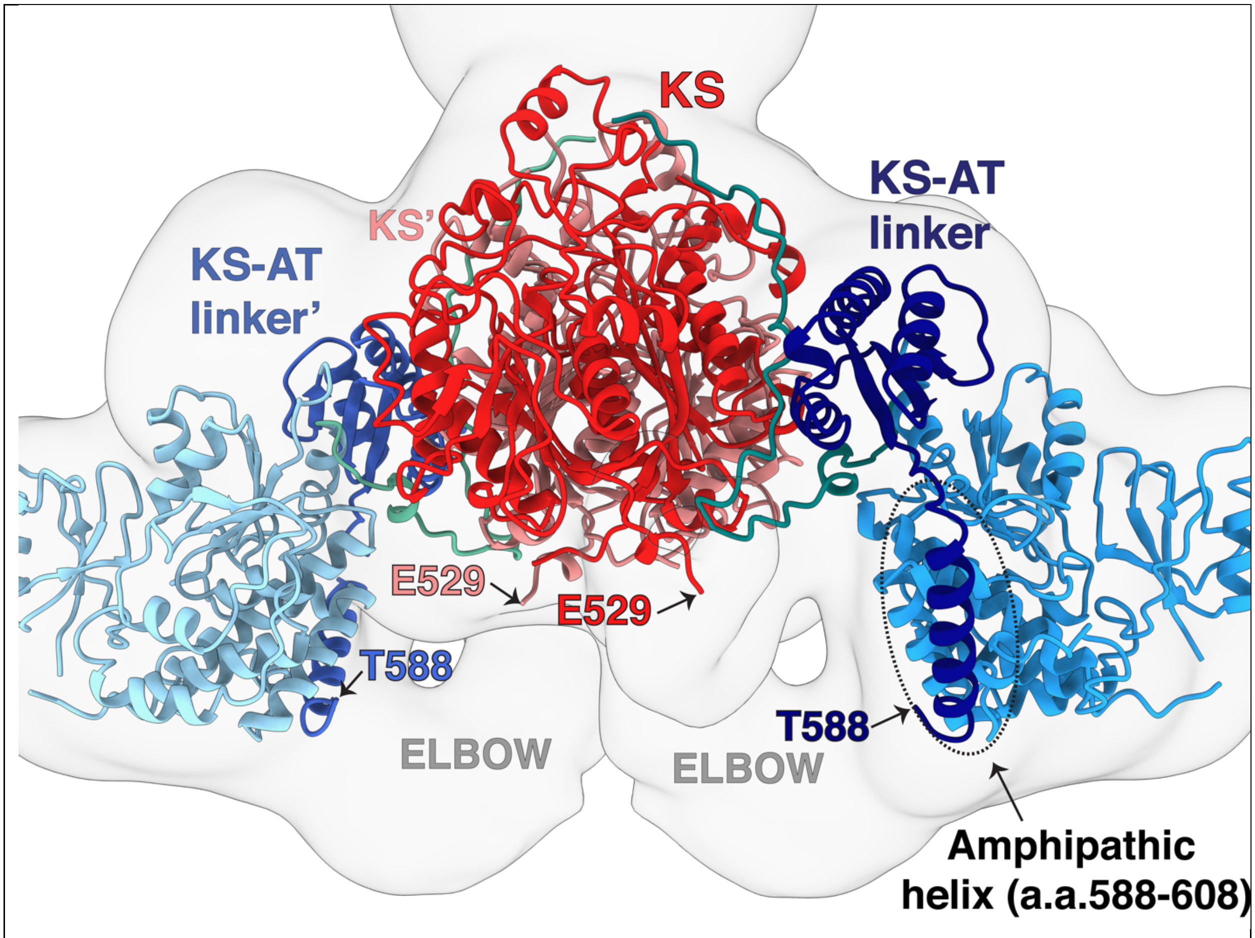

**Fig S13. View of elbow density, and amphipathic helix.** Residues between E529 (KS) and T588 (KS-AT linker) are not modeled due to low resolution density in the region. This region is presumably connected by a flexible “elbow” region shown in the low-resolution composite density map that connects the end of E529 and beginning of T588. In the figure, ACP1 structure is omitted for clarity. An amphipathic helix (588-608) found only in mycobacterial Pks13s leads into the KS-AT linker domain (circled and labeled).

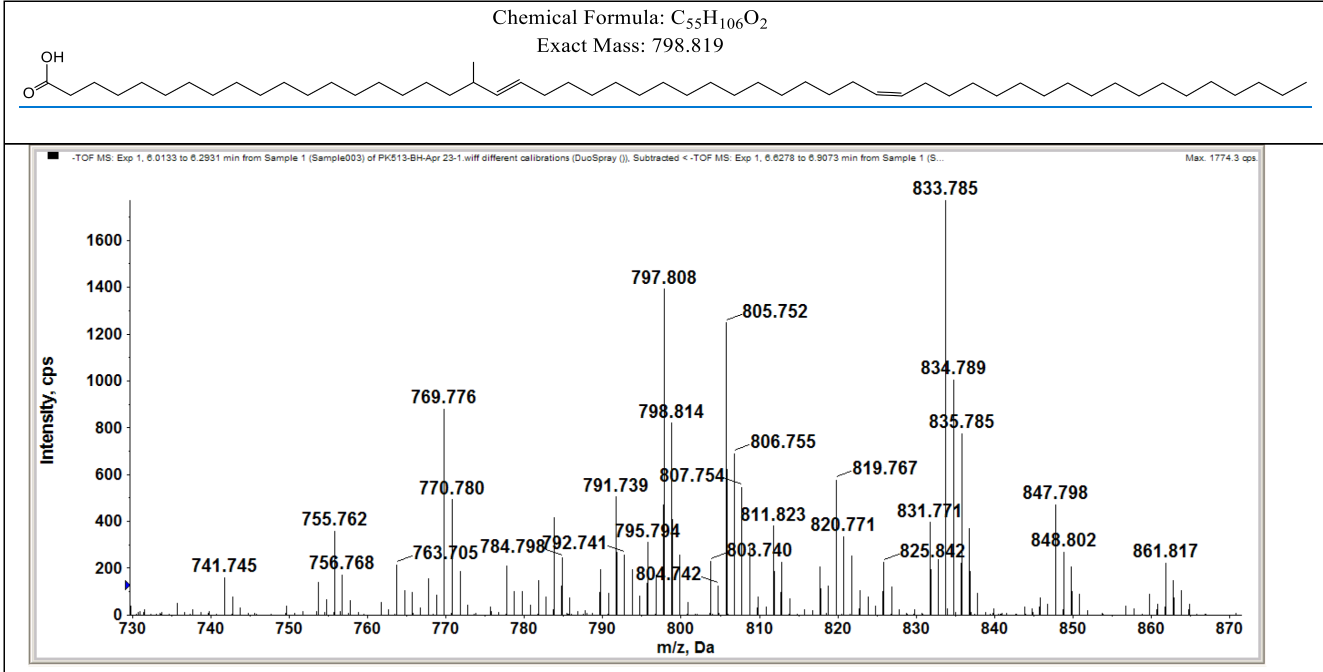

**Fig S14. The extremely long chain, C51-C57 in length, α2-meromycolate** (A) has its major component with the formula $C_{55}H_{106}O_2$ with exact mass 798.819 Da. consistent with the substrate seen attached covalently to the active site of the KS domain in density. (B) Mass spectra define the major species as the $C_{55}H_{106}O_2$. The α-2 chain contains a single methyl branch following a trans double bond, and a single cis double bond[21].

Chemical Formula: $C_{40}H_{78}O_2$
Exact Mass: 590.600

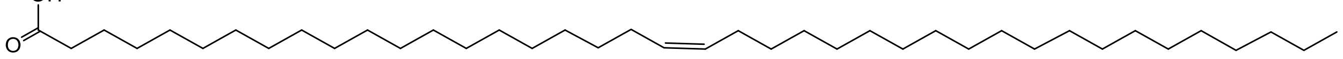

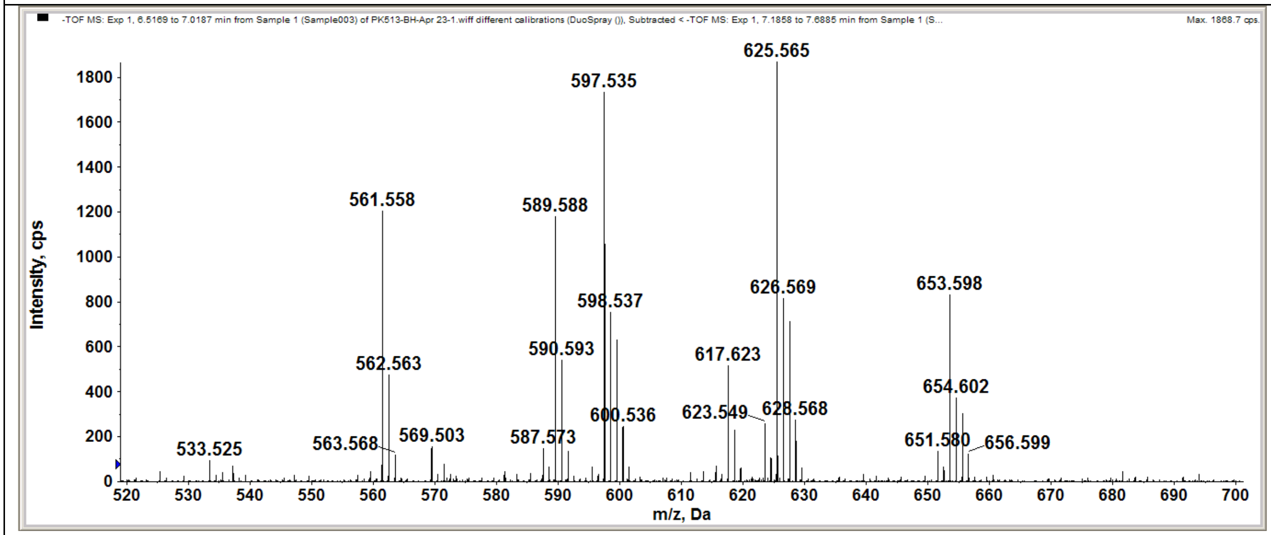

**Fig S15. The C40 chain fatty acid**, (A) C36-C42 chain α'-mycolyl chain has its major component with the formula $C_{40}H_{78}O_2$ and exact mass 590.600, a cis-monounsaturated fatty acid similar in length to tetracontilic acid. (B) Mass spectra for the species are shown. This chain is consistent with the α' mycolate found in *M.smegmatis* but not found in *M.tuberculosis*.

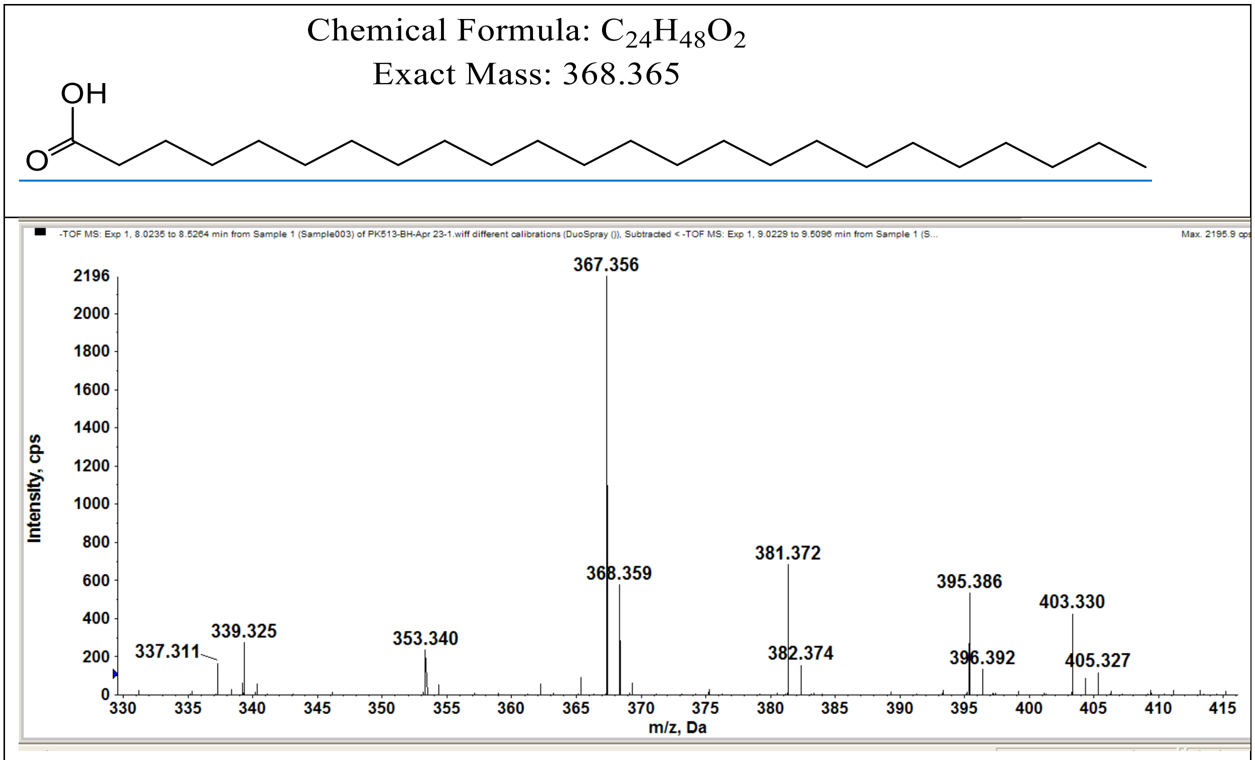

**Fig S16. The α-branch chain** (A) Chains of the expected lengths (C22-C26) for the α-branch, α-carboxy fatty acid as seen attached to S798 of the AT domain. (B) The predominant species is for C24 chain of mass 368.365 Da. The mass of the $[M+Cl]^-$ observed is 403.330 Da. That calculated from the deduced structure indicated above is 403.334

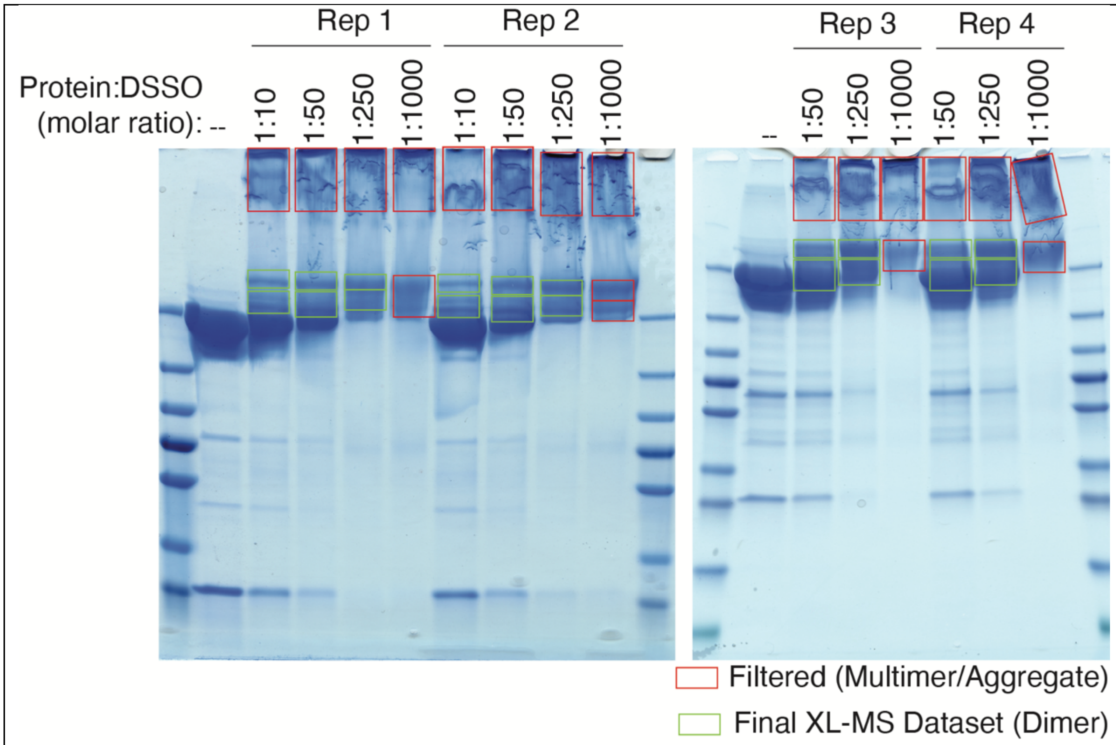

**Fig S17. SDS-PAGE separation of Pks13 cross-linked with increasing amounts of DSSO.** Pks13 was cross-linked with increasing amounts of DSSO for 30min at 37°C (Rep1, 3, and 4) or 4°C (Rep2), separated on a 4-20% SDS-PAGE gel, and stained with MS safe blue stain. All non-homomeric bands were cut as indicated and analyzed by XL-MS[3] (see methods for details). Mapping linkages to the cryoEM structures (Figure S18) demonstrated high levels of distance violations for cross-links that could be mapped. Given that most of these were found in bands associated with higher amounts of cross-linking and potential aggregates, data corresponding to these bands (shown in Red) were filtered out of the Final dataset. Bands that correspond to the lower molecular weight products are shown in green.

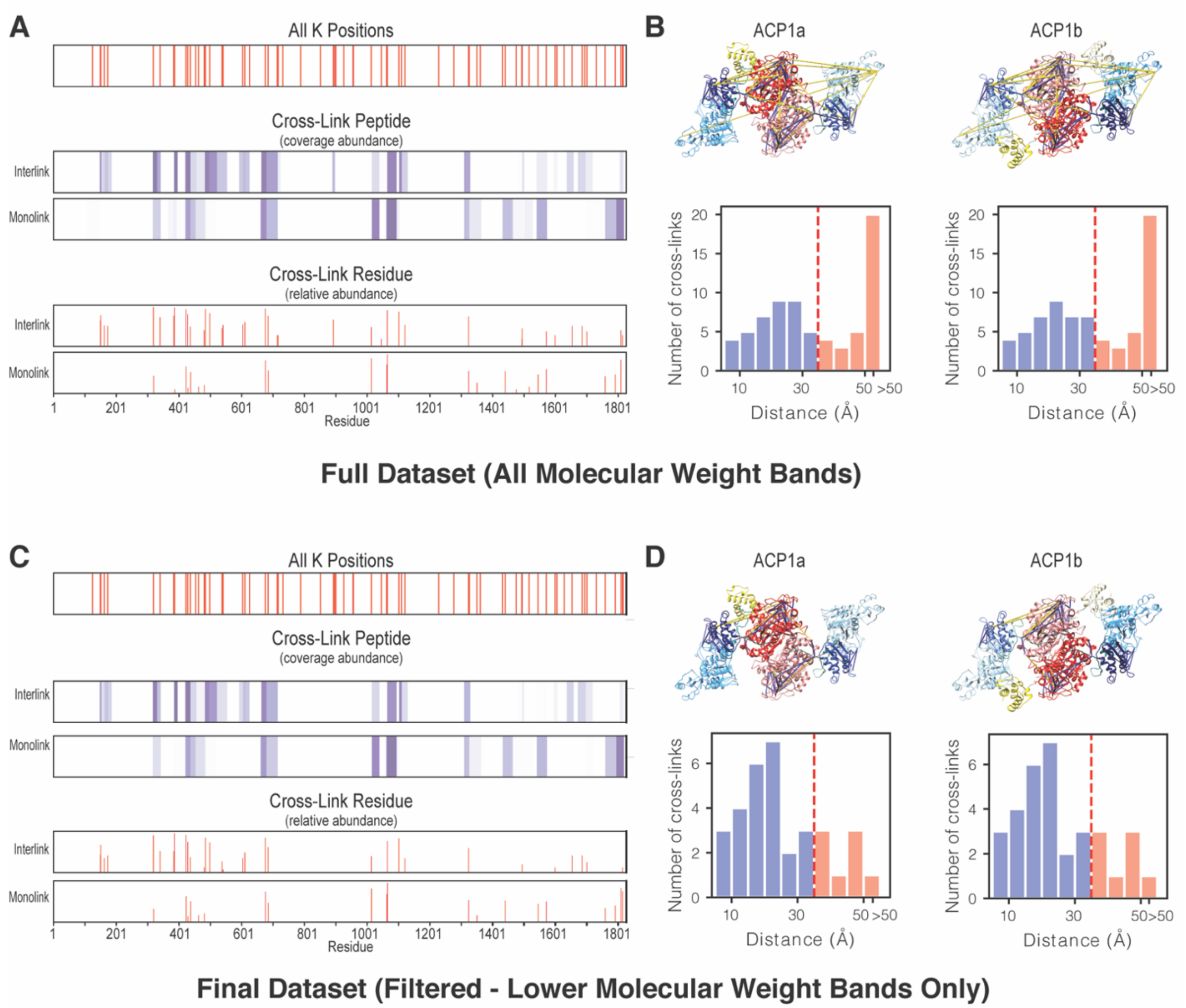

**Fig S18. Summary of Pks13 DSSO cross-links.** (A, C) For the full (A) and filtered (C) data-sets all Pks13 lysine positions (K positions) are plotted on the top graph. The cross-linked peptide coverage is shown for interlinks and monolinks as an intensity plot with darker purple indicating higher spectral count. The corresponding relative abundance for each cross-link by residue is shown as a line graph in the two plots below. (B, D) DSSO interlinked residues are mapped to the two cryo-EM structures (shown on the top) for the full (**B**) and filtered (**D**) data-sets. Linkages that satisfy the expected distance (<35Å) are shown in blue, and linkages that violate this distance are shown in yellow. Histograms that plot the distances for each of the unique linkages are shown for each structure (shown on the bottom).

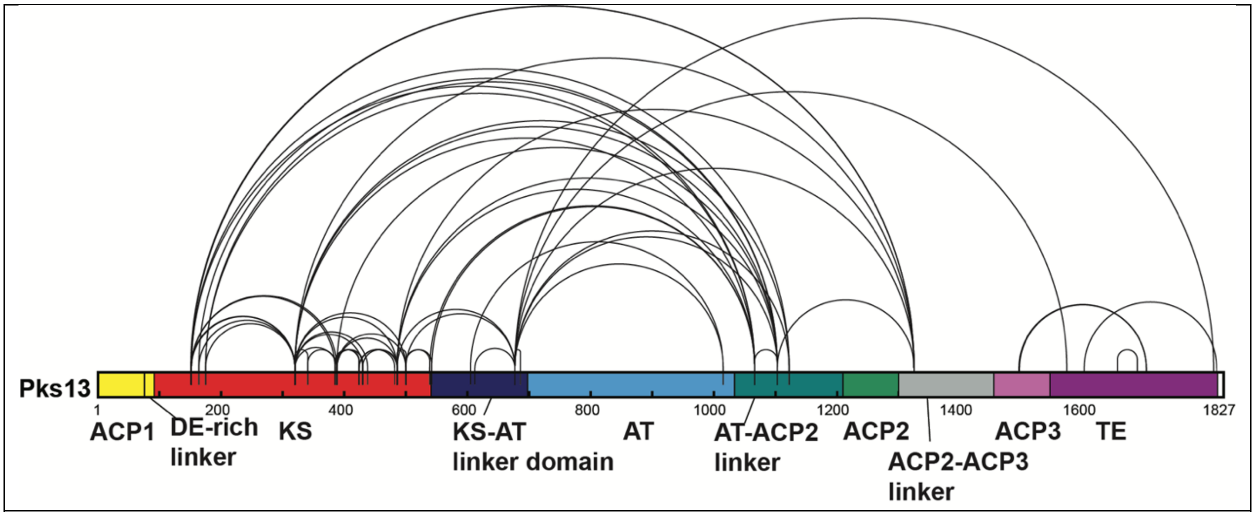

**Fig. S19. Linkage map depicting all unique K-K linkages of DSSO cross-linked Pks13.** The 57 unique K-K inter-linked residues are summarized in this linkage map as black lines connecting two Pks13 lysine residues. Shown is a cartoon diagram of the primary sequence structure of Pks13 with domains labeled in yellow (ACP1), red (KS), blue (AT), green (ACP2), light purple (ACP3), and purple (TE). Labeled also are the linker domains in between them.

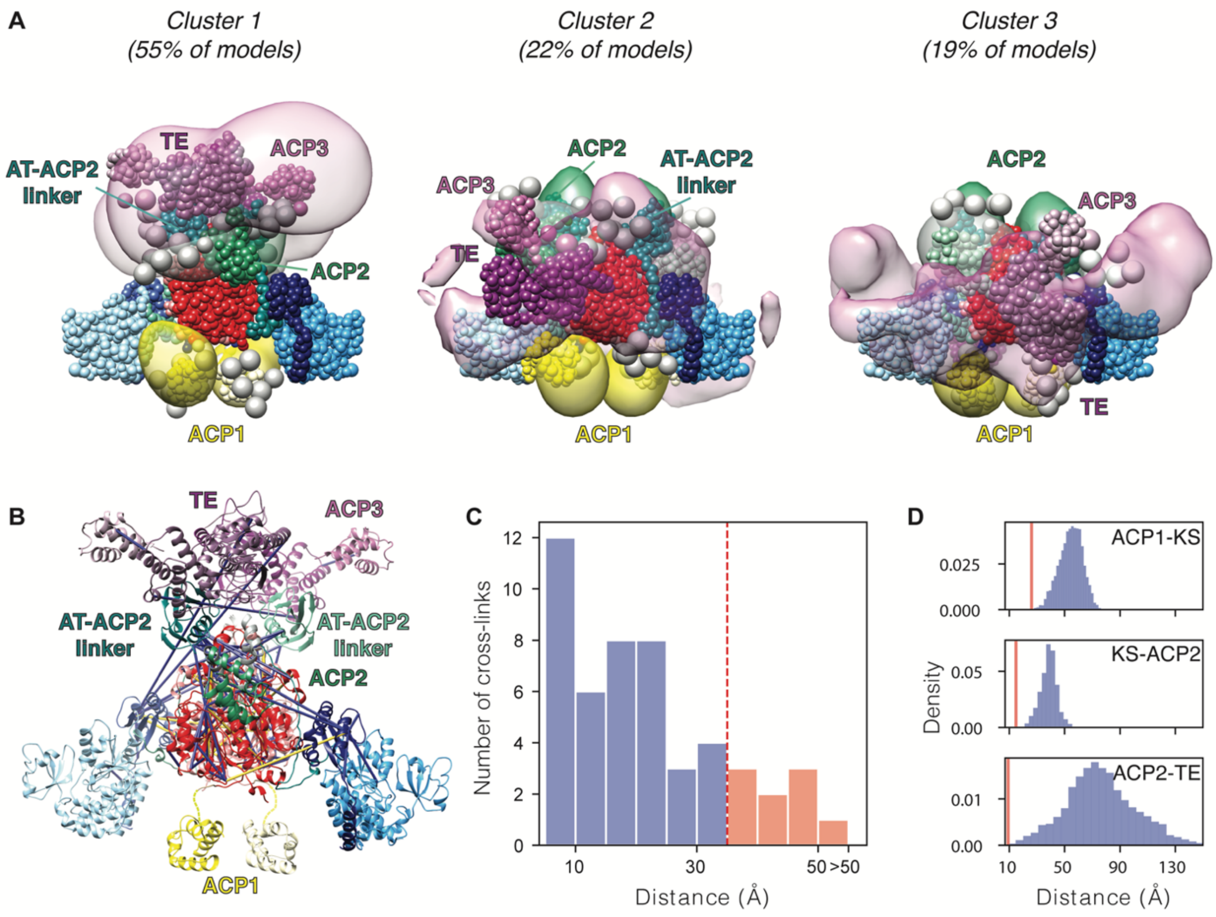

**Fig S20: Integrative structure modeling of the full-length Pks13 dimer.** (A) The localization probability density of the ensemble of structures is shown with representative (centroid) structure from the ensemble embedded within it. The structured and unstructured regions are represented as beads, using the same color scheme as Fig. 1. (B) Detail of cross-links mapped to the centroid structure of the integrative model of the Pks13 dimer. Satisfied and violated cross-links shown in blue and yellow, respectively. A cross-link is classified as satisfied if the Cα–Cα distance spanned by the cross-linked residues in any of the models of the cluster is less than 35 Å. (C) Histogram showing the distribution of the cross-linked Cα–Cα distances in the Pks13 dimer integrative structures. (D) Histogram showing the Cα–Cα distances between the active sites of the ACP1-KS, KS-ACP2, and ACP2-TE domains. The shortest distance (red lines) are 26, 14.7, and 9.2 Å between the ACP1-KS, KS-ACP2, and ACP2-TE domains, respectively.

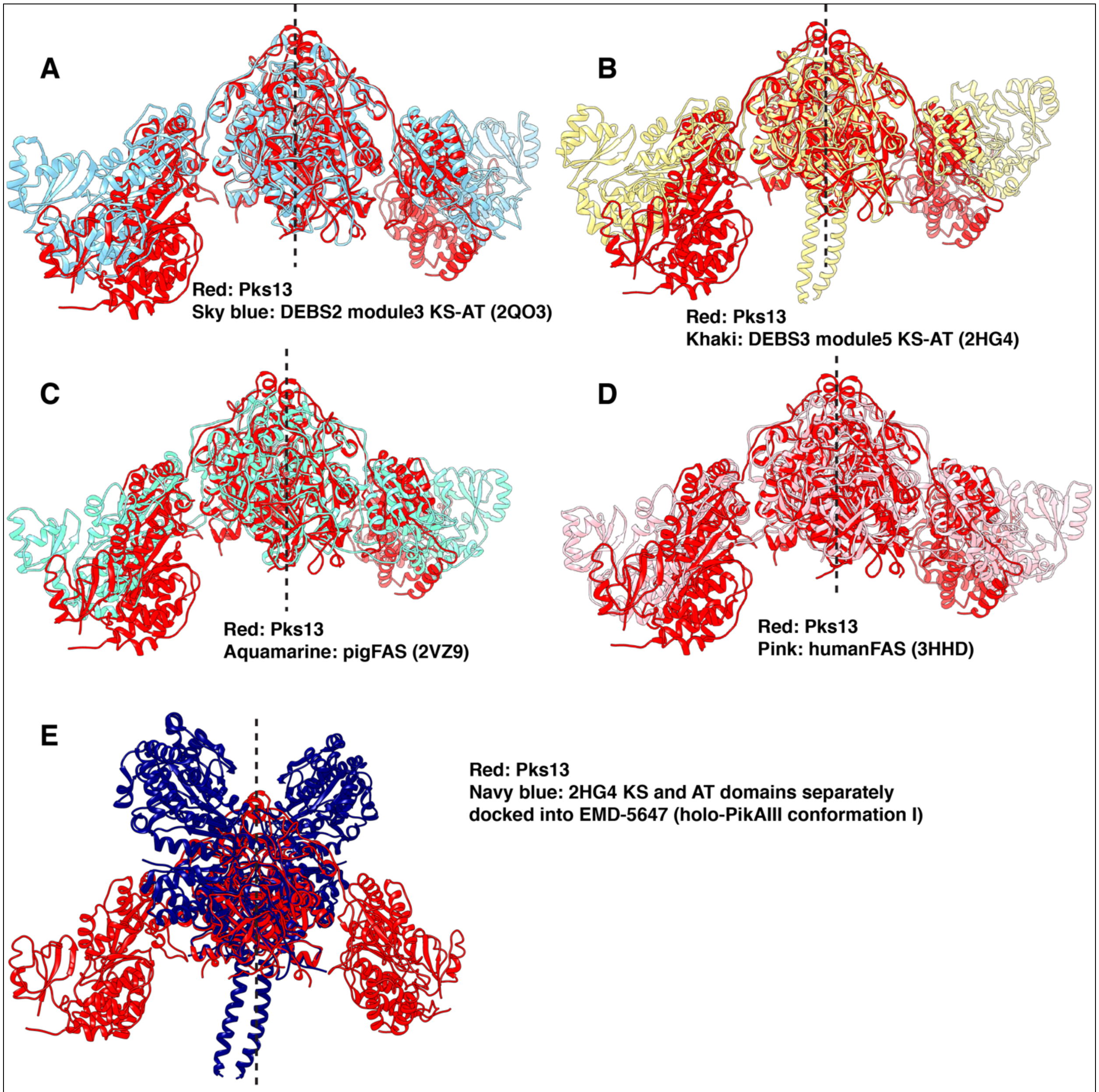

**Fig S21. Pks13 KS-AT comparisons to other structures that include the di-domain KS-AT.** (A) Comparison of Pks13 KS-AT (red) to that in module 3 of DEBS 2 (blue) (PDB: 2QO3). (B) Comparison of KS-AT in Pks13 (red) to that in module 5 of DEBS 3 (khaki) (2HG4). (C) Comparison of KS-AT in Pks13 to that in porcine FAS (Aquamarine) (2VZ9). (D) Comparison of KS-AT in Pks13 (red) to that of human FAS (pink) (3HHD). (E) Comparison of

KS-AT in Pks13 (red) to a structure of PikAIII (blue) obtained by fitting domain homology models into cryo-EM density.

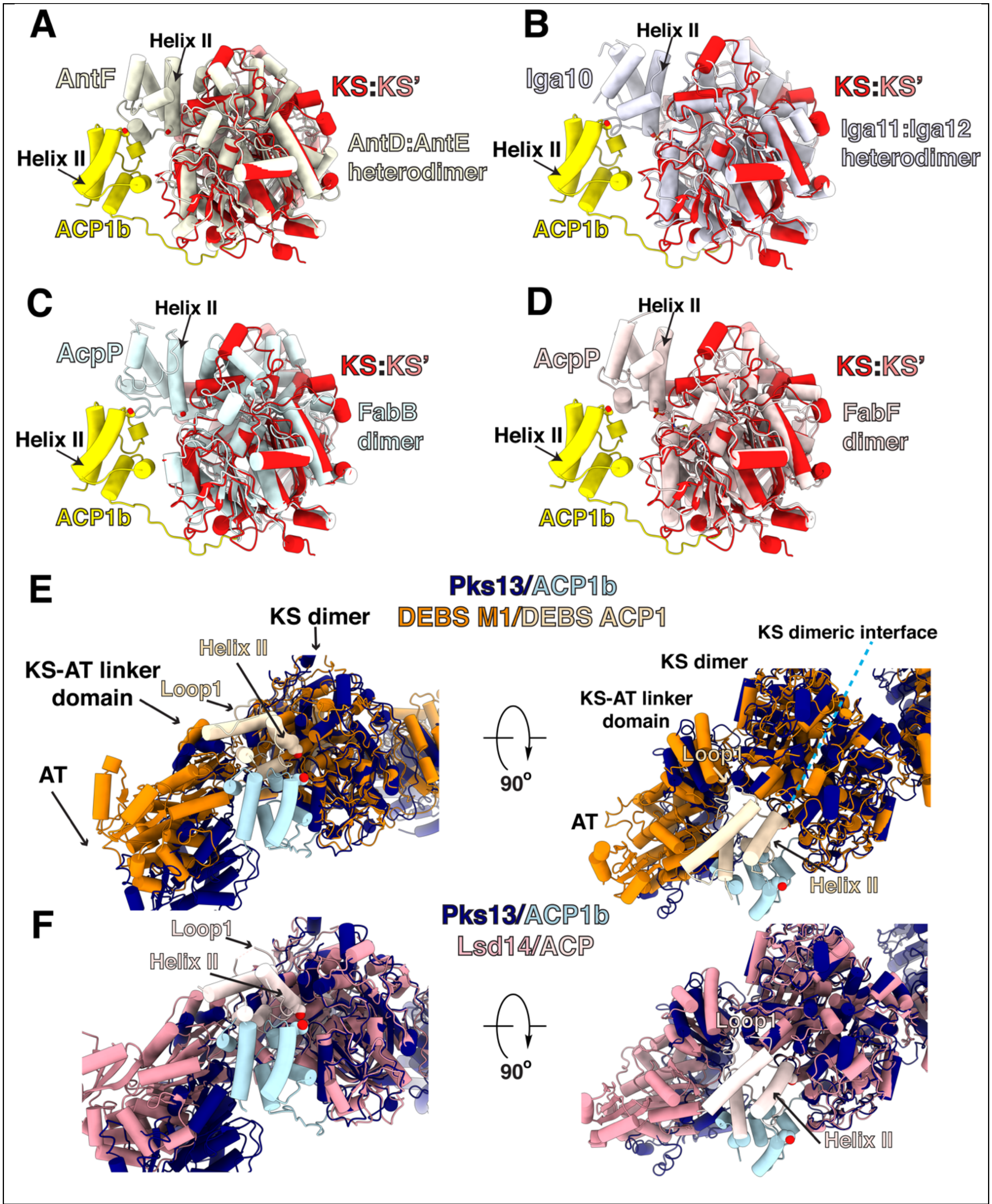

**Fig S22: Comparison of ACP:KS binding modes.** Superposition of Pks13 ACP1b:KS structure with trans-acting ACPs (A) AntF, (B) Iga10, and (C, D) AcpP and their cognate KSs (same color as cognate ACPs). These trans-acting ACPs interact with their KSs similarly via their helix II, with their Ppant-binding serine at the N-terminus of helix II shown in stick representation. Superposition of ACP1b:KS structure with cis-acting ACPs from (E) DEBS module 1and (F) Lsd14 PKSs interacting with their cognate PKS module. KS interacting epitopes loop1 and helix II on the ACP are labeled, and Ppant-binding serine is shown as sphere at the N-terminus of helix II.

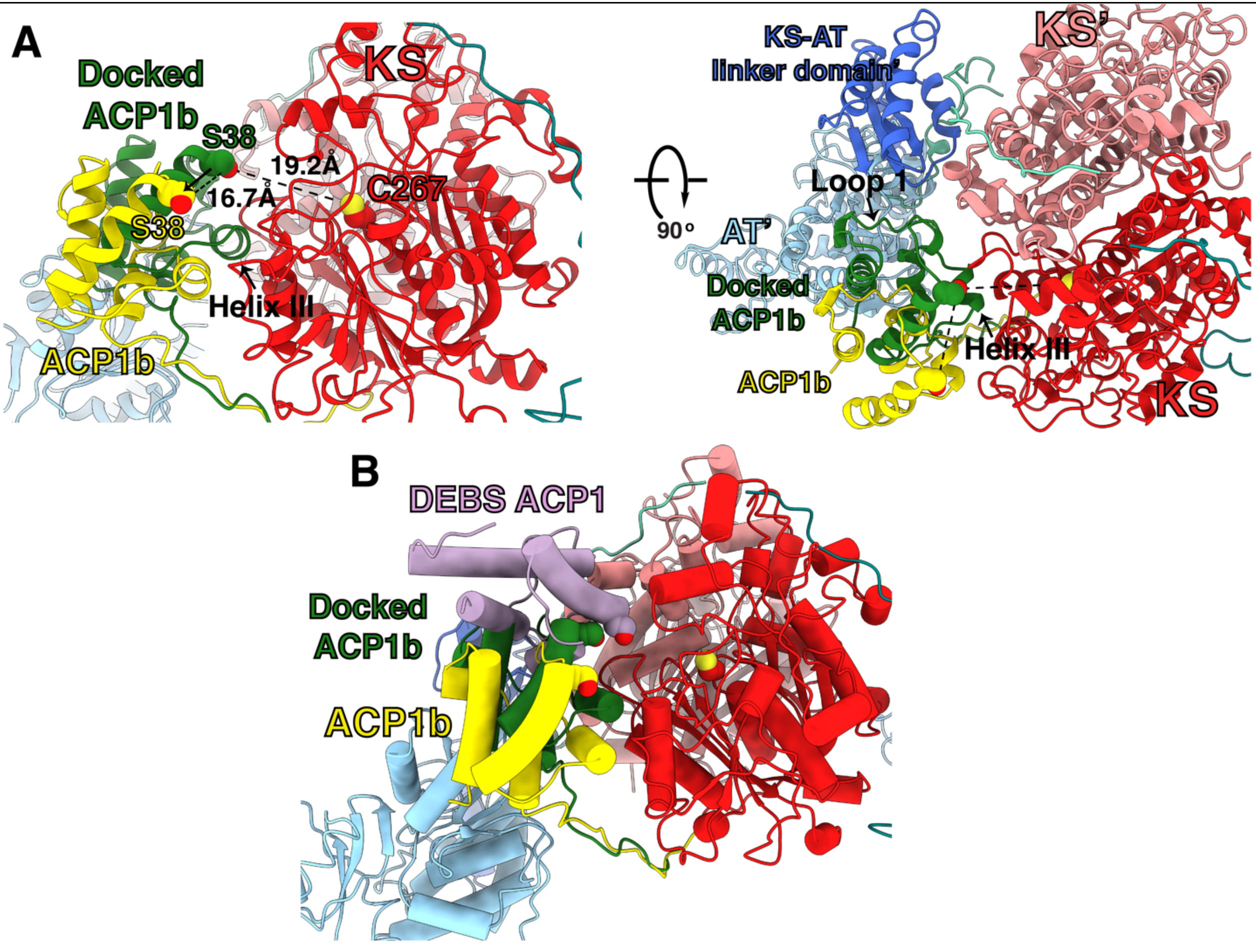

**Fig S23. ACP1b substrate delivery docked position.** (A) S38 of docked ACP1b (green), which is required to deliver the meromycolate to the KS, is removed ~17Å from the S38 of our ACP1b structure (yellow). From the docked ACP1b position (green), S38 is ~19Å from the catalytic C267 of KS. In this docked position, helix III and loop 1 make contacts with the KS-KS' dimer and KS-AT linker' and AT' domains of the opposite protomer. (B) Superposition of ACP1b, ACP1b docked position, and DEBS ACP1. Docked ACP1b orientation is ~90° rotated from the DEBS module 1 ACP binding mode.

**Table S1. Table of microscope parameters**

| **Acquisition parameter** | |
|---|---|
| Microscope | Titan Krios G3 |
| Imaging system | K3 BioQuantum, 20 eV slit |
| Illumination mode | Nanoprobe, parallel |
| High tension | 300 kV |
| Pixel size (super-resolution) | 0.4175 Å $pix^{-1}$ |
| Under focus range | 0.7 - 1.5 µm |
| Accumulated electron dose | 67 $e^{-}$ $Å^{-2}$ |
| Exposure time | 5.9 s |
| Frames per movie | 117 |
| Movies collected | 7567 |
| Acquisition software | SerialEM |

**Table S2. Model refinement statistics.**

| | KS | ACP1a - KS-AT | ACP1b-KS-AT | AT in | AT out |
|---|---|---|---|---|---|
| Resolution based on Q scores | 1.8 | 2.4 | 3.0 | 2.8 | 2.4 |
| Map to model FSC (0.5 threshold, masked/unmasked) (Å) | 2.0/2.0 | 2.5/2.8 | 2.7/2.9 | 2.8/3.9 | 2.8/3.8 |
| **Model composition** | | | | | |
| No. Atoms | 6638 | 15307 | 14775 | 3748 | 3748 |
| No. Residues | 882 | 1944 | 1941 | 484 | 484 |
| No. Water | 0 | 0 | 0 | 0 | 0 |
| No. Ligands | 2 | 3 | 2 | 0 | 0 |
| **Bonds (RMSD)** | | | | | |
| Length (Å) (# > 4 σ) | 0.004 (0) | 0.005 (2) | 0.005 (0) | 0.002 (1) | 0.007 (0) |
| Angles (°) (# > 4 σ) | 0.641 (0) | 0.603 (0) | 0.569 (4) | 0.451 (2) | 0.792 (0) |
| MolProbity score | 1.07 | 1.52 | 1.77 | 1.51 | 1.78 |
| Clash score | 1.60 | 5.85 | 8.70 | 4.69 | 6.43 |
| **Ramachandran plot (%)** | | | | | |
| Outliers | 0.00 | 0.00 | 0.00 | 0.00 | 0.00 |
| Allowed | 2.96 | 2.79 | 4.35 | 3.96 | 6.46 |
| Favored | 97.04 | 97.21 | 95.65 | 96.04 | 93.54 |
| **Rama-Z (Rama. Plot Z-score, RMSD)** | | | | | |
| Whole (N = 866) | 0.22 (0.27) | 0.83 (0.18) | 0.61 (0.19) | 0.46 (0.40) | 1.16 (0.37) |
| Helix (N = 304) | 0.81 (0.27) | 0.25 (0.17) | 0.15 (0.19) | 1.88 (0.36) | 0.13 (0.36) |
| Sheet (N = 128) | 0.81 (0.47) | 0.00 (0.33) | 0.02 (0.35) | 0.09 (0.73) | 0.27 (0.70) |
| Loop (N = 434) | 0.47 (0.28) | 0.88 (0.20) | 0.94 (0.20) | 1.43 (0.42) | 1.92 (0.38) |
| Rotamer outliers (%) | 0.73 | 1.18 | 0.00 | 0.00 | 0.00 |
| Cβ outliers (%) | 0.00 | 0.00 | 0.00 | 0.00 | 0.00 |
| **Peptide plane (%)** | | | | | |

| | | | | | |
|---|---|---|---|---|---|
| Cis proline/general | 0.0/0.0 | 0.0/0.0 | 0.0/0.0 | 0.0/0.0 | 0.0/0.0 |
| Twisted proline/general | 0.0/0.2 | 0.0/0.1 | 0.0/0.0 | 0.0/0.0 | 0.0/0.0 |
| CaBLAM outliers (%) | 1.37 | 1.56 | 1.30 | 2.94 | 3.15 |

**Table S3 Resolution of the map by region and focused refinements based on Q scores**

| Refine | KS-AT dimer | KS substrate | AT focused | AT substrate | AT in | AT out | ACP1a | Ppant ACP1 | ACP1b | ACP1a-KS-AT | KS substrate | ACP1b-KS-AT | KS substrate |
|---|---|---|---|---|---|---|---|---|---|---|---|---|---|
| Res (Å) | 1.8 | 2.4 | 2.0 | 2.9 | 2.8 | 2.4 | 3.6 | 3.5 | 4.6 | 2.4 | 2.4 | 3.0 | 2.4 |

| **Table S4. Exact mass measurement of fatty acids bound to Pks13** | | | | | |
|---|---|---|---|---|---|
| Fatty acids are detected as deprotonated $[M-H]^-$ or chloride adduct $[M+Cl]^-$ ions | | | | | |
| | | $[M-H]^-$ | | $[M+Cl]^-$ | |
| | | Expected *m/z* | Observed *m/z* | Expected *m/z* | Observed *m/z* |
| **C22-C26** | C22 | 339.327 | 339.325 | | |
| | C23 | 363.343 | 363.341 | | |
| | C24 | 367.358 | 367.356 | 403.334 | 403.330 |
| | C25 | 381.374 | 381.372 | | |
| | C26 | 395.389 | 395.385 | | |
| **C36-C42** | C36 | 533.530 | 533.525 | 569.506 | 569.503 |
| | C38 | 561.562 | 561.558 | 597.538 | 597.535 |
| | C40 | 589.593 | 589.588 | 625.569 | 625.565 |
| | C42 | 617.624 | 617.623 | 653.600 | 653.598 |
| **C51-C57** | C51 | 741.747 | 741.745 | 777.726 | 777.724 |
| | C52 | 755.765 | 755.762 | 791.741 | 791.739 |
| | C53 | 769.781 | 769.776 | 805.757 | 805.752 |
| | C54 | 783.796 | 783.793 | 819.772 | 819.767 |
| | C55 | 797.812 | 797.808 | 833.788 | 833.785 |
| | C56 | 811.828 | 811.823 | 847.804 | 847.798 |
| | C57 | 825.843 | 825.842 | 861.819 | 861.817 |

**Table S5. Table with description and scores for overall DSSO cross-linked peptides for all processed files including inter-linked, mono-linked, and single peptides.**

**Table S6. Summary of all unique Pks13 inter-linked residues from the full dataset.**

**Table S7. Summary of unique Pks13 inter-linked residues excluding files from highly cross-linked samples.**

**Table S8. Metadata all DSSO cross-linked Pks13 XL-MS files.**

Table 9: Summary of the Pks13 dimer integrative structure modeling

| | |
|---|---|
| **1) Gathering information** | |
| *Prior models* | 2-fold symmetry derived from cryo-EM structure |
| *Physical principles and statistical preferences* | Excluded volume<br>Sequence connectivity |
| *Experimental data* | 57 DSS0<br>Atomic structure from cryo-EM map; PDB TBD |
| **2) Representing the system** | |
| *Atomic (structured) components* | Pks13: 1-76, 89-228, 229-232, 233-529, 588-835, 836-838, 839-1074, 1078-1173, 1238-1356, 1461-1535, 1539-1816, 1-76, 89-228, 229-232, 233-529, 588-835, 836-838, 839-1074, 1078-1173, 1238-1356, 1461-1535, 1539-1816 |
| *Unstructured components* | Pks13: 77-88, 530-587, 1075-1077, 1174-1237, 1357-1460, 1536-1538, 77-88, 530-587, 1075-1077, 1174-1237, 1357-1460, 1536-1538 |
| *Resolution of structured components* | 1 [R1] residue per bead |
| *Resolution of unstructured components* | 10 [R10] residues per bead |
| *Structural coverage* | 86.77 % |
| *Rigid body (RB) definitions* | RB1: Pks13$_{1-76}$<br>RB2: Pks13$_{89-228}$,Pks13$_{229-232}$,Pks13$_{233-529}$,Pks13$_{588-835}$,Pks13$_{8}$[illegible]<br>RB3: Pks13$_{1078-1173}$<br>RB4: Pks13$_{1238-1356}$<br>RB5: Pks13$_{1461-1535}$<br>RB6: Pks13$_{1539-1816}$<br>RB7: Pks13$_{1-76}$<br>RB8: Pks13$_{89-228}$,Pks13$_{229-232}$,Pks13$_{233-529}$,Pks13$_{588-835}$,Pks13$_{8}$[illegible]<br>RB9: Pks13$_{1078-1173}$<br>RB10: Pks13$_{1238-1356}$<br>RB11: Pks13$_{1461-1535}$<br>RB12: Pks13$_{1539-1816}$ |
| *Resolution of disordered regions* | 10 [R10] residues per bead |
| *Composition (number of copies of Pks13)* | 2 |
| *Spatial restraints encoded into scoring function* | Excluded volume; applied to the R1 representation<br>Sequence connectivity; applied to the R1 representation<br>Cross-link restraints; applied to the R1 representation |
| **3.1) Enumeration of threading of degrees of freedom** | |
| **3.2) Structural Sampling** | |
| *Sampling method* | Replica Exchange Gibbs sampling, based on Metropolis Monte Carlo |
| *Replica exchange temperature range* | 1.0 - 2.5 |
| *Number of replicas* | 8 |
| *Number of runs* | 100 |
| *Number of structures generated* | 2500000 |
| *Movers for flexible string of bead* | Random translation up to 4.0 Å |
| *CPU time* | 22 hours on 80 processors |
| **4.1) Validating the threading models** | |
| **4.2) Validating the Pom152 ring models** | |
| **Models selected for validation** | |
| *Number of models after equilibration* | 2500000 |
| *Number of models that satisfy the input information* | 229554 |
| *Number of structures in samples A/B* | 112886/116668 |
| *p-value of non-parametric Kolmogorov-Smirnov two-sample test* | 0.02 (threshold p-value $> 0.05$) |
| *Kolmogorov-Smirnov two-sample test statistic, D* | 1.0 |
| **Thoroughness of the structural sampling** | |
| *Sampling precision* | 61.37 Å |
| *Homogeneity of proportions $\chi^2$ test (p-value)/Cramers V value* | 0.000/0.067 (thresholds: p-value$>$0.05 OR Cramer's V$<$0.1) |
| *Number of clusters* | 3 |
| *Cluster precisions* | cluster 1 : 55.0 %<br>cluster 2 : 21.7 % |

**Movie S1 caption: Morph of the difference between the AT domain positions relative to the KS domain.** The angular variation is 7° about the region in between the AT' and KS domains. The color code from the manuscript is followed for domains. The two extreme state structures are represented as both AT domains moved toward the viewer, versus AT domains away from the viewer. This angular change may be synchronized with movement of the other domains that might include the ACP1,2 or the TE domains. Scene 1) Showing the density envelope at low resolution to emphasize the domain movement, extrapolated between the first frame (gray) when the AT positions are toward the viewer on both sides, to the last frame (yellow) when the AT domains move away. ACP1 appears better represented in position ACP1b as AT moves away on the left side. Thus it appears that he AT hinges outward and away on the left side relative to the KS as ACP1 docks onto the ACP1b position on that same left side. Scene 2) This is followed by a cartoon showing the 'liquorice' style rendering of the motion of one AT domain relative to the KS dimer from focused alignments, viewed from the side nearest the AT domain with the KS dimer kept constant in position to emphasize the domain shift. This is followed by the same events viewed down the 2-fold axis between the KS domains that illustrates the 7° rotation and twist associated with this motion. Scene 3) The high-resolution density map is superposed onto the KS-AT dimer. The 2-fold axis is vertical. The density for the hinge domain on top of the KS dimer is not interpretable at atomic resolution. This is followed by a low-resolution envelope around the 'cartoon' representation of the -ACP1b-(yellow)-KS (red)–AT (blue). The two -fold axis is vertical. The envelope for the ACP1-KS- linking DE-rich sequence that leads into the KS domain is clearly delineated. The view moves around to the ACP1b location and shows the attachment point of the Ppant arm at S38 (yellow and red (oxygen of the serine) sphere, yellow bonded atoms). The bonded atoms indicate density for atomic positions (yellow spheres and sticks) into a tunnel ('delivery tunnel') into the KS active site. The covalently bound mycolic acid substrate rises vertically from the sulfur of C267 (small red sphere bonded to the substrate yellow carbon atoms) at the far end of this 'delivery' tunnel. The view pulls back to give the relative context of the AT active site serine bound to its $\alpha$-carboxy-fatty acid substrate. Resolved atoms are included in the structure.

Bibliography

1. Harding, E. WHO global progress report on tuberculosis elimination. *Lancet Respir. Med.* **8,** 19 (2020).
2. Dulberger, C. L., Rubin, E. J. & Boutte, C. C. The mycobacterial cell envelope - a moving target. *Nat. Rev. Microbiol.* **18,** 47–59 (2020).
3. Grzegorzewicz, A. E. *et al.* Inhibition of mycolic acid transport across the Mycobacterium tuberculosis plasma membrane. *Nat. Chem. Biol.* **8,** 334–341 (2012).
4. Ioerger, T. R. *et al.* Identification of new drug targets and resistance mechanisms in Mycobacterium tuberculosis. *PLoS One* **8,** e75245 (2013).
5. Zuber, B. *et al.* Direct visualization of the outer membrane of mycobacteria and corynebacteria in their native state. *J. Bacteriol.* **190,** 5672–5680 (2008).
6. Hoffmann, C., Leis, A., Niederweis, M., Plitzko, J. M. & Engelhardt, H. Disclosure of the mycobacterial outer membrane: cryo-electron tomography and vitreous sections reveal the lipid bilayer structure. *Proc. Natl. Acad. Sci. USA* **105,** 3963–3967 (2008).
7. Sani, M. *et al.* Direct visualization by cryo-EM of the mycobacterial capsular layer: a labile structure containing ESX-1-secreted proteins. *PLoS Pathog.* **6,** e1000794 (2010).
8. Sacchettini, J. C., Rubin, E. J. & Freundlich, J. S. Drugs versus bugs: in pursuit of the persistent predator Mycobacterium tuberculosis. *Nat. Rev. Microbiol.* **6,** 41–52 (2008).
9. Portevin, D. *et al.* A polyketide synthase catalyzes the last condensation step of mycolic acid biosynthesis in mycobacteria and related organisms. *Proc. Natl. Acad. Sci. USA* **101,** 314–319 (2004).
10. Gavalda, S. *et al.* The Pks13/FadD32 crosstalk for the biosynthesis of mycolic acids in Mycobacterium tuberculosis. *J. Biol. Chem.* **284,** 19255–19264 (2009).
11. Trivedi, O. A. *et al.* Enzymic activation and transfer of fatty acids as acyl-adenylates in mycobacteria. *Nature* **428,** 441–445 (2004).
12. Léger, M. *et al.* The dual function of the Mycobacterium tuberculosis FadD32 required for mycolic acid biosynthesis. *Chem. Biol.* **16,** 510–519 (2009).
13. Marrakchi, H., Lanéelle, M.-A. & Daffé, M. Mycolic acids: structures, biosynthesis, and beyond. *Chem. Biol.* **21,** 67–85 (2014).
14. Herbst, D. A. *et al.* The structural organization of substrate loading in iterative polyketide synthases. *Nat. Chem. Biol.* **14,** 474–479 (2018).
15. Khosla, C. From active sites to machines: A challenge for enzyme chemists. *Isr J Chem* **59,** 37–40 (2019).
16. Meniche, X. *et al.* Subpolar addition of new cell wall is directed by DivIVA in mycobacteria. *Proc. Natl. Acad. Sci. U. S. A.* **111,** E3243-51 (2014).
17. Aggarwal, A. *et al.* Development of a Novel Lead that Targets M. tuberculosis Polyketide Synthase 13. *Cell* **170,** 249–259.e25 (2017).
18. Pan, H. *et al.* Crystal structure of the priming beta-ketosynthase from the R1128 polyketide biosynthetic pathway. *Structure* **10,** 1559–1568 (2002).
19. Tang, Y., Kim, C.-Y., Mathews, I. I., Cane, D. E. & Khosla, C. The 2.7-Angstrom crystal structure of a 194-kDa homodimeric fragment of the 6-deoxyerythronolide B synthase. *Proc. Natl. Acad. Sci. USA* **103,** 11124–11129 (2006).
20. Krissinel, E. & Henrick, K. Inference of macromolecular assemblies from crystalline state. *J. Mol. Biol.* **372,** 774–797 (2007).
21. Barry, C. E. *et al.* Mycolic acids: structure, biosynthesis and physiological functions. *Prog. Lipid Res.* **37,** 143–179 (1998).

22. Byers, D. M. & Gong, H. Acyl carrier protein: structure-function relationships in a conserved multifunctional protein family. *Biochem Cell Biol* **85,** 649–662 (2007).
23. Robbins, T., Kapilivsky, J., Cane, D. E. & Khosla, C. Roles of conserved active site residues in the ketosynthase domain of an assembly line polyketide synthase. *Biochemistry* **55,** 4476–4484 (2016).
24. Klaus, M. *et al.* Protein-Protein Interactions, Not Substrate Recognition, Dominate the Turnover of Chimeric Assembly Line Polyketide Synthases. *J. Biol. Chem.* **291,** 16404–16415 (2016).
25. Bergeret, F. *et al.* Biochemical and structural study of the atypical acyltransferase domain from the mycobacterial polyketide synthase Pks13. *J. Biol. Chem.* **287,** 33675–33690 (2012).
26. Gande, R. *et al.* Acyl-CoA carboxylases (accD2 and accD3), together with a unique polyketide synthase (Cg-pks), are key to mycolic acid biosynthesis in Corynebacterianeae such as Corynebacterium glutamicum and Mycobacterium tuberculosis. *J. Biol. Chem.* **279,** 44847–44857 (2004).
27. Lucas, X., Bauzá, A., Frontera, A. & Quiñonero, D. A thorough anion-π interaction study in biomolecules: on the importance of cooperativity effects. *Chem. Sci.* **7,** 1038–1050 (2016).
28. George, K. M., Yuan, Y., Sherman, D. R. & Barry, C. E. The biosynthesis of cyclopropanated mycolic acids in Mycobacterium tuberculosis. Identification and functional analysis of CMAS-2. *J. Biol. Chem.* **270,** 27292–27298 (1995).
29. Yuan, Y. & Barry, C. E. A common mechanism for the biosynthesis of methoxy and cyclopropyl mycolic acids in Mycobacterium tuberculosis. *Proc. Natl. Acad. Sci. USA* **93,** 12828–12833 (1996).
30. Piersimoni, L. & Sinz, A. Cross-linking/mass spectrometry at the crossroads. *Anal. Bioanal. Chem.* **412,** 5981–5987 (2020).
31. Kaake, R. M. *et al.* A new in vivo cross-linking mass spectrometry platform to define protein-protein interactions in living cells. *Mol. Cell Proteomics* **13,** 3533–3543 (2014).
32. Kao, A. *et al.* Development of a novel cross-linking strategy for fast and accurate identification of cross-linked peptides of protein complexes. *Mol. Cell Proteomics* **10,** M110.002212 (2011).
33. Merkley, E. D. *et al.* Distance restraints from crosslinking mass spectrometry: mining a molecular dynamics simulation database to evaluate lysine-lysine distances. *Protein Sci.* **23,** 747–759 (2014).
34. Kaake, R. M. *et al.* Characterization of an A3G-VifHIV-1-CRL5-CBFβ Structure Using a Cross-linking Mass Spectrometry Pipeline for Integrative Modeling of Host-Pathogen Complexes. *Mol. Cell Proteomics* **20,** 100132 (2021).
35. Sali, A. From integrative structural biology to cell biology. *J. Biol. Chem.* **296,** 100743 (2021).
36. Saltzberg, D. J. *et al.* Using Integrative Modeling Platform to compute, validate, and archive a model of a protein complex structure. *Protein Sci.* **30,** 250–261 (2021).
37. Saltzberg, D. *et al.* Modeling biological complexes using integrative modeling platform. *Methods Mol. Biol.* **2022,** 353–377 (2019).
38. Kim, S. J. *et al.* Integrative structure and functional anatomy of a nuclear pore complex. *Nature* **555,** 475–482 (2018).
39. Jumper, J. *et al.* Highly accurate protein structure prediction with AlphaFold. *Nature* **596,** 583–589 (2021).

40. Varadi, M. *et al.* AlphaFold Protein Structure Database: massively expanding the structural coverage of protein-sequence space with high-accuracy models. *Nucleic Acids Res.* **50,** D439–D444 (2022).
41. Russel, D. *et al.* Putting the pieces together: integrative modeling platform software for structure determination of macromolecular assemblies. *PLoS Biol.* **10,** e1001244 (2012).
42. Edwards, A. L., Matsui, T., Weiss, T. M. & Khosla, C. Architectures of whole-module and bimodular proteins from the 6-deoxyerythronolide B synthase. *J. Mol. Biol.* **426,** 2229–2245 (2014).
43. Tang, Y., Chen, A. Y., Kim, C.-Y., Cane, D. E. & Khosla, C. Structural and mechanistic analysis of protein interactions in module 3 of the 6-deoxyerythronolide B synthase. *Chem. Biol.* **14,** 931–943 (2007).
44. Khosla, C., Tang, Y., Chen, A. Y., Schnarr, N. A. & Cane, D. E. Structure and mechanism of the 6-deoxyerythronolide B synthase. *Annu. Rev. Biochem.* **76,** 195–221 (2007).
45. Khosla, C. Structures and mechanisms of polyketide synthases. *J. Org. Chem.* **74,** 6416–6420 (2009).
46. Maier, T., Leibundgut, M. & Ban, N. The crystal structure of a mammalian fatty acid synthase. *Science* **321,** 1315–1322 (2008).
47. Pappenberger, G. *et al.* Structure of the human fatty acid synthase KS-MAT didomain as a framework for inhibitor design. *J. Mol. Biol.* **397,** 508–519 (2010).
48. Leibundgut, M., Jenni, S., Frick, C. & Ban, N. Structural basis for substrate delivery by acyl carrier protein in the yeast fatty acid synthase. *Science* **316,** 288–290 (2007).
49. Whicher, J. R. *et al.* Structural rearrangements of a polyketide synthase module during its catalytic cycle. *Nature* **510,** 560–564 (2014).
50. Dutta, S. *et al.* Structure of a modular polyketide synthase. *Nature* **510,** 512–517 (2014).
51. Mindrebo, J. T. *et al.* Gating mechanism of elongating β-ketoacyl-ACP synthases. *Nat. Commun.* **11,** 1727 (2020).
52. Bräuer, A. *et al.* Structural snapshots of the minimal PKS system responsible for octaketide biosynthesis. *Nat. Chem.* **12,** 755–763 (2020).
53. Du, D. *et al.* Structural basis for selectivity in a highly reducing type II polyketide synthase. *Nat. Chem. Biol.* **16,** 776–782 (2020).
54. Milligan, J. C. *et al.* Molecular basis for interactions between an acyl carrier protein and a ketosynthase. *Nat. Chem. Biol.* **15,** 669–671 (2019).
55. Cogan, D. P. *et al.* Mapping the catalytic conformations of an assembly-line polyketide synthase module. *Science* **374,** 729–734 (2021).
56. Bagde, S. R., Mathews, I. I., Fromme, J. C. & Kim, C.-Y. Modular polyketide synthase contains two reaction chambers that operate asynchronously. *Science* **374,** 723–729 (2021).
57. Kao, C. M., Pieper, R., Cane, D. E. & Khosla, C. Evidence for two catalytically independent clusters of active sites in a functional modular polyketide synthase. *Biochemistry* **35,** 12363–12368 (1996).
58. Tsai, S. C. *et al.* Crystal Structure of the Macrocycle-forming Thioesterase Domain of Erythromycin Polyketide Synthase (DEBS TE). (2002).
59. Gokulan, K., Aggarwal, A., Shipman, L., Besra, G. S. & Sacchettini, J. C. Mycobacterium tuberculosis acyl carrier protein synthase adopts two different pH-dependent structural conformations. *Acta Crystallogr. Sect. D, Biol. Crystallogr.* **67,** 657–669 (2011).
60. Keatinge-Clay, A. T. *et al.* Catalysis, specificity, and ACP docking site of Streptomyces coelicolor malonyl-CoA:ACP transacylase. *Structure* **11,** 147–154 (2003).

61. Tsai, S.-C., Lu, H., Cane, D. E., Khosla, C. & Stroud, R. M. Insights into Channel Architecture and Substrate Specificity from Crystal Structures of Two Macrocycle-Forming Thioesterases of Modular Polyketide Synthases[†, ‡]. *Biochemistry* **41,** 12598–12606 (2002).
62. Portevin, D. *et al.* The acyl-AMP ligase FadD32 and AccD4-containing acyl-CoA carboxylase are required for the synthesis of mycolic acids and essential for mycobacterial growth: identification of the carboxylation product and determination of the acyl-CoA carboxylase components. *J. Biol. Chem.* **280,** 8862–8874 (2005).
63. La Rosa, V. *et al.* MmpL3 is the cellular target of the antitubercular pyrrole derivative BM212. *Antimicrob. Agents Chemother.* **56,** 324–331 (2012).
64. Poce, G. *et al.* Improved BM212 MmpL3 inhibitor analogue shows efficacy in acute murine model of tuberculosis infection. *PLoS One* **8,** e56980 (2013).
65. LaCava, J., Jiang, H. & Rout, M. P. Protein Complex Affinity Capture from Cryomilled Mammalian Cells. *J. Vis. Exp.* (2016). doi:10.3791/54518
66. Zheng, S. Q. *et al.* MotionCor2: anisotropic correction of beam-induced motion for improved cryo-electron microscopy. *Nat. Methods* **14,** 331–332 (2017).
67. Punjani, A., Rubinstein, J. L., Fleet, D. J. & Brubaker, M. A. cryoSPARC: algorithms for rapid unsupervised cryo-EM structure determination. *Nat. Methods* **14,** 290–296 (2017).
68. Grant, T., Rohou, A. & Grigorieff, N. cisTEM, user-friendly software for single-particle image processing. *Elife* **7,** e35383 (2018).
69. Zivanov, J. *et al.* New tools for automated high-resolution cryo-EM structure determination in RELION-3. *Elife* **7,** (2018).
70. Pettersen, E. F. *et al.* UCSF Chimera—a visualization system for exploratory research and analysis. *J. Comput. Chem.* **25,** 1605–1612 (2004).
71. Goddard, T. D. *et al.* UCSF ChimeraX: Meeting modern challenges in visualization and analysis. *Protein Sci.* **27,** 14–25 (2018).
72. Sanchez-Garcia, R. *et al.* DeepEMhancer: a deep learning solution for cryo-EM volume post-processing. *Commun. Biol.* **4,** 874 (2021).
73. Terwilliger, T. C., Ludtke, S. J., Read, R. J., Adams, P. D. & Afonine, P. V. Improvement of cryo-EM maps by density modification. *Nat. Methods* **17,** 923–927 (2020).
74. Afonine, P. V. *et al.* Real-space refinement in Phenix for cryo-EM and crystallography. *BioRxiv* (2018). doi:10.1101/249607
75. Kelley, L. A., Mezulis, S., Yates, C. M., Wass, M. N. & Sternberg, M. J. E. The Phyre2 web portal for protein modeling, prediction and analysis. *Nat. Protoc.* **10,** 845–858 (2015).
76. Emsley, P., Lohkamp, B., Scott, W. G. & Cowtan, K. Features and development of Coot. *Acta Crystallogr. Sect. D, Biol. Crystallogr.* **66,** 486–501 (2010).
77. Afonine, P. V. *et al.* Real-space refinement in PHENIX for cryo-EM and crystallography. *Acta Crystallogr. D Struct. Biol.* **74,** 531–544 (2018).
78. Pravda, L. *et al.* MOLEonline: a web-based tool for analyzing channels, tunnels and pores (2018 update). *Nucleic Acids Res.* **46,** W368–W373 (2018).
79. Tan, B. K. *et al.* Discovery of a cardiolipin synthase utilizing phosphatidylethanolamine and phosphatidylglycerol as substrates. *Proc. Natl. Acad. Sci. USA* **109,** 16504–16509 (2012).
80. Chambers, M. C. *et al.* A cross-platform toolkit for mass spectrometry and proteomics. *Nat. Biotechnol.* **30,** 918–920 (2012).
81. Kessner, D., Chambers, M., Burke, R., Agus, D. & Mallick, P. ProteoWizard: open source software for rapid proteomics tools development. *Bioinformatics* **24,** 2534–2536 (2008).
82. Perez-Riverol, Y. *et al.* The PRIDE database and related tools and resources in 2019:

improving support for quantification data. *Nucleic Acids Res.* **47,** D442–D450 (2019).
83. Baker, P. R. & Chalkley, R. J. MS-viewer: a web-based spectral viewer for proteomics results. *Mol. Cell Proteomics* **13,** 1392–1396 (2014).
84. Alber, F. *et al.* Determining the architectures of macromolecular assemblies. *Nature* **450,** 683–694 (2007).
85. Viswanath, S., Chemmama, I. E., Cimermancic, P. & Sali, A. Assessing exhaustiveness of stochastic sampling for integrative modeling of macromolecular structures. *Biophys. J.* **113,** 2344–2353 (2017).
86. Madeira, F. *et al.* The EMBL-EBI search and sequence analysis tools APIs in 2019. *Nucleic Acids Res.* **47,** W636–W641 (2019).
87. Waterhouse, A. M., Procter, J. B., Martin, D. M. A., Clamp, M. & Barton, G. J. Jalview Version 2--a multiple sequence alignment editor and analysis workbench. *Bioinformatics* **25,** 1189–1191 (2009).
